# A brain–behavior generative model links individual brain dynamics to perceptual decisions

**DOI:** 10.64898/2026.10.16.759120

**Authors:** Chaoming Wang, Guochun Yang, Shaonan Wang, Huan Luo

## Abstract

Perceptual decision speed varies across individuals and trials. Explaining this variability requires linking individual brain anatomy, neural dynamics, and behavior, yet whole-brain models typically reproduce brain signals without behavior, whereas behavioral models lack anatomical constraints. Here we introduce a brain–behavior generative model in which whole-brain dynamics on each participant’s structural connectome generate both single-trial electroencephalography (EEG) and reaction times. Fitted to 17 participants performing motion discrimination, the models reproduced individual eventrelated potentials and reaction-time distributions. Joint fitting enabled the models to predict whether held-out trials were relatively fast or slow within each stimulus condition. *In silico* silencing further predicted where, how, and when each person’s brain controls decision speed. Sensorimotor regions affected decision speed consistently across most participants, whereas the same parietal region had opposite effects in different participants. These opposite effects reflected whether the region acted mainly through network dynamics or the decision readout. Temporal analyses suggested that early activity largely set decision speed, whereas late EEG mainly tracked the unfolding decision. These results show that brain–behavior generative modeling can link individual brain dynamics to perceptual decisions and yield testable predictions of why decision speed varies across individuals and trials.

## Introduction

Understanding how individual brains generate behavior remains a fundamental challenge in cognitive neuroscience. Although humans rely on shared computational principles^1^, these computations unfold within neural systems that differ in anatomy, connectivity, and dynamics^2,3^, giving rise to substantial variability in behavior^4^. Perceptual decision-making provides a compelling example^5^: even when sensory inputs are identical, individuals respond at different speeds^6^, and the same individual’s response time varies from trial to trial^7,8^. Such variability suggests that decision speed is not determined solely by external evidence or a fixed computational rule, but emerges from the interaction between task-related computations and the evolving dynamics of each brain^9,10^. Explaining this phenomenon therefore requires a mechanistic account of how an individual neural system transforms sensory information into decisions over time. Crucially, such an account must move beyond identifying neural activity correlated with faster or slower responses^11,12^; it should reveal where, how, and when changes in a person’s brain dynamics causally shape that person’s decision speed.

Existing approaches capture important components of this problem but leave the link between individual brain dynamics and behavior unresolved. Biophysical models of EEG describe how interacting neural populations generate electrophysiological signals and evoked responses^13,14^. When embedded within whole-brain architectures constrained by structural connectomes^15,16^, these models can reproduce resting-state functional organization^17,18^ and even individual-specific stimulation-evoked EEG patterns^19,20^. However, they generally lack a mechanistic link to behavior^21^ and therefore cannot explain why a particular brain produces a particular decision speed. Conversely, cognitive models such as the drift–diffusion model (DDM) successfully explain choices and reaction times as evidence accumulation toward a decision bound^22,23^, but abstract away the neural substrates that implement this computation. Recent artificial-intelligence methods model neural and behavioral dynamics jointly^24^, for example in neurocognitive models that fit EEG and behavior at the single-trial level^25,26^, yet their latent decision variables lack anatomical identity and cannot predict how perturbing a specific brain system would alter behavior. Thus, a generative framework that links individual-specific, anatomy-constrained neural dynamics to both neural signals and behavior remains unavailable.

Here we introduce a brain–behavior generative model (BGM; Fig. 1) that provides this link. The model simulates whole-brain dynamics on each participant’s structural connectome and uses this single generative process to produce both single-trial EEG and decision behavior. Neural activity is mapped onto EEG through a leadfield and onto reaction times through activity in task-relevant decision circuits. By jointly fitting single-trial EEG and reaction times, the BGM allows neural signals and behavior to constrain the same underlying dynamical system, rather than treating them as separate observations requiring independent explanations. This shared generative structure enables EEG to reveal a trial-specific latent variable, while behavioral variability constrains how that variable influences decision formation. Once fitted to an individual brain, the model provides a mechanistic substrate for predicting how targeted perturbations of neural dynamics would alter that person’s decision speed, thereby linking brain anatomy, neural dynamics, and behavior within a unified computational framework.

**Figure 1:**
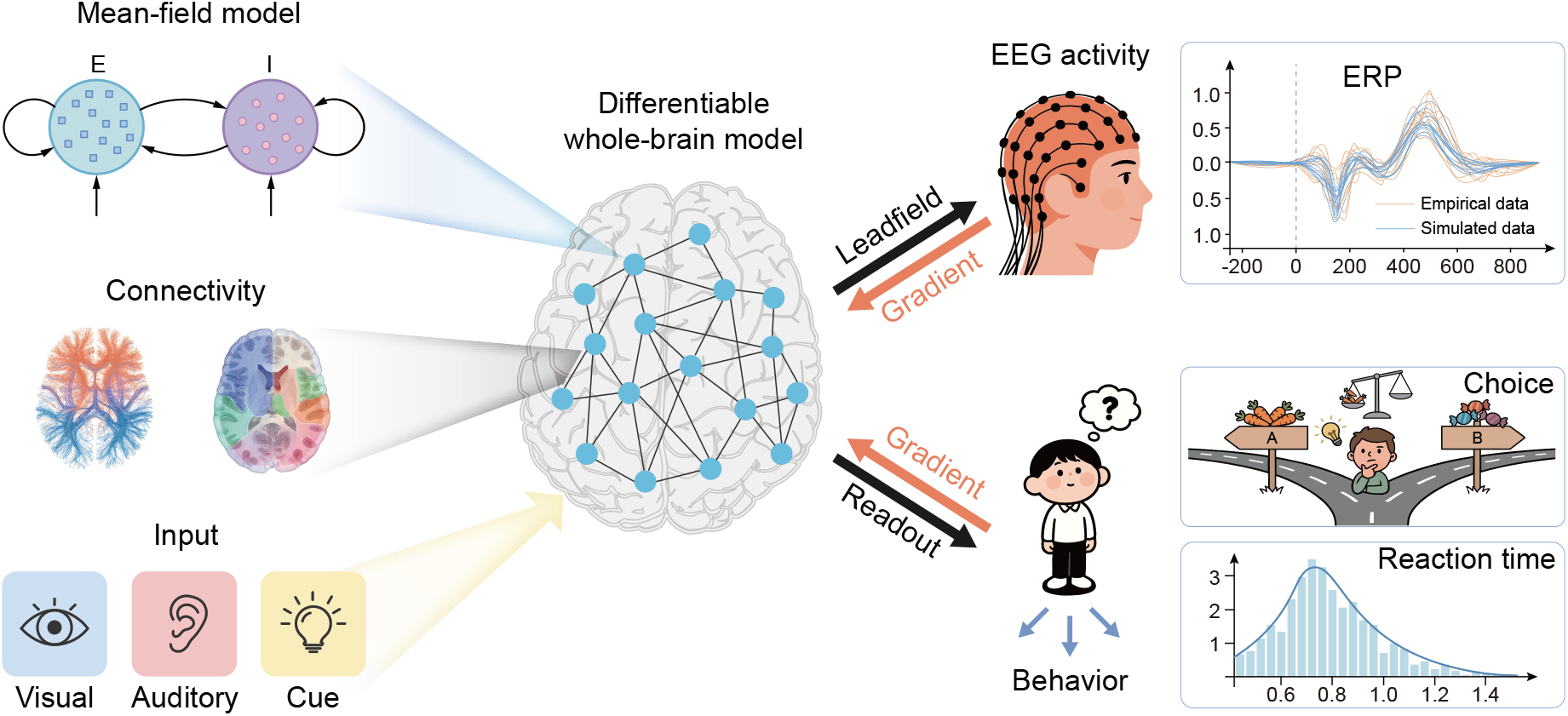
The brain–behavior generative model (BGM). Visual stimulus input drives a whole-brain network of mean-field excitatory–inhibitory (E–I) units coupled on the participant’s own structural connectome, and the regional dynamics are read out twice. A leadfield projects regional currents to scalp EEG, which is compared with that participant’s recorded waveforms, and a readout of activity in identified brain regions produces decision behavior. Gradients from the EEG and behavioral likelihoods (red) propagate back through the same forward computation and update one parameter set in a single end-to-end joint fit.

We applied the BGM to 17 participants performing motion discrimination in the VEPCON dataset^27^. First, each person’s model reproduced that person’s event-related potentials and reaction-time distribution, and its evoked responses depended on that person’s own connectome. Second, the trial-specific latent variable inferred from the whole EEG epoch predicted held-out reaction times within stimulus conditions, but only when reaction times entered the fit. Third, *in silico* silencing predicted where, how, and when each person’s brain sets decision speed. Silencing sensorimotor regions generally slowed decisions, whereas silencing parietal accumulator regions could slow or accelerate them, depending on whether the region acted through the network or through its weight in the decision readout. Silencing restricted to the first 200 ms reproduced the direction of each participant’s strongest effect. Silencing confined to the P3 window^28,29^ barely changed decision speed when it targeted that strongest region, and when it targeted the parietal accumulator regions it changed the late scalp signal more than early silencing did. These findings connect shared decision computations to individual brain dynamics and yield testable, person-specific predictions of how changing neural activity would change a perceptual decision.

## Results

### Each person’s model reproduces their ERP and reaction times

In the VEPCON dataset^27^, participants reported whether a random-dot kinematogram contained coherent motion while 128-channel EEG was recorded. For each of 17 participants, we built one model on that person’s anatomy (Fig. 2a) and asked whether it could produce both that person’s EEG and their decisions. A 200-ms sensory pulse, set by the stimulus condition alone, enters primary visual cortex (V1) and the middle temporal motion area (MT/V5). The pulse drives a Wilson–Cowan network^30^ of excitatory–inhibitory population pairs in the 83 regions of the Lausanne-2018 parcellation^31^. A fixed leadfield, the head model that maps each region’s current onto each electrode, turns this activity into 128-channel EEG. In parallel, a decision pool of 10 regions known to encode and accumulate motion evidence^11,32,33^ forms the pooled evidence, which sets the drift rate of an accumulator rising toward a bound^23^. Accumulator noise, added at every moment, makes two trials with the same drift reach the bound at different times (Fig. 2a). The time it first reaches the bound, plus a non-decision time *t*_0_, is the reaction time. The connectome, leadfield, and conduction delays are calibrated once from that person’s anatomy and never fit. A trial-specific latent variable, introduced in the next section, adds an input that differs between trials of the same condition. EEG and reaction times are fit jointly, in one objective (Methods).

**Figure 2:**
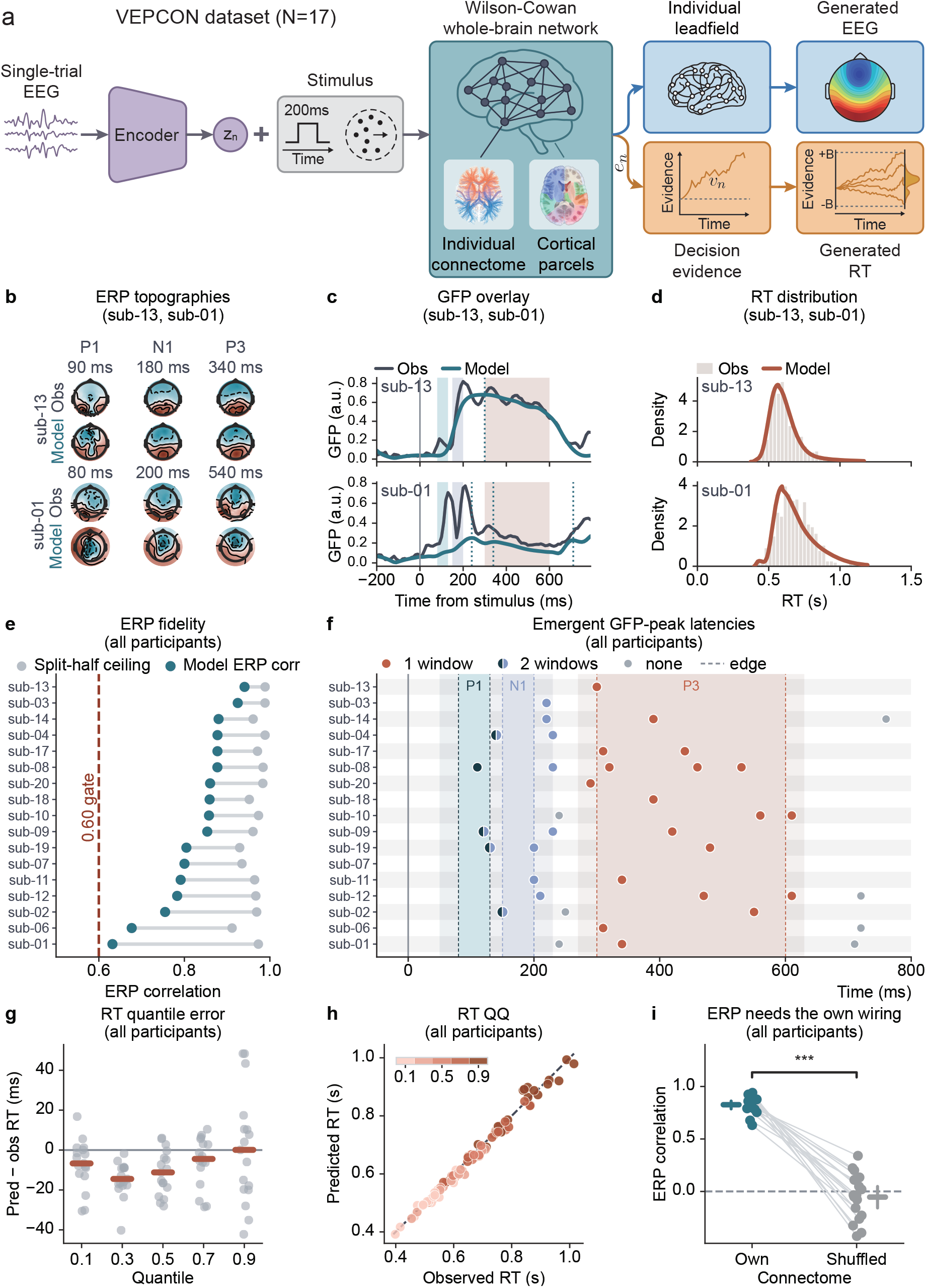
One person’s model generates both that person’s EEG and their reaction times. **a**, BGM forward model. Single-trial EEG is encoded into the trial-specific latent variable *z*_*n*_ (Eq. 12), which is added to a 200-ms sensory pulse and drives an 83-region Wilson–Cowan network on the individual connectome; the leadfield and the feedback-inhibition parameters are fixed. Figure 2: Given *z*_*n*_ the EEG path (leadfield) and the decision path (pooled evidence *e*_*n*_(*t*) setting the drift and so the reaction time, Eqs. 20, 21, and 24) are conditionally independent; both are fit jointly under one evidence lower bound (Eqs. 25 and 26), with choice marginalized out. **b–d**, The best-fitting (sub-13, *r* = 0.941, top) and worst-fitting (sub-01, *r* = 0.632, bottom) participants. **b**, Observed and model scalp topographies at the best-matched time point inside each window (sub-13: 90, 180, and 340 ms; sub-01: 80, 200, and 540 ms); each map is scaled to its own peak, and amplitudes are read from **c. c**, Observed and model condition-mean global field power (GFP) from −200 to 800 ms; dotted lines mark the model’s GFP peaks. **d**, Observed reaction-time histogram with the model’s predicted density. **e**, Each participant’s correlation between model and observed event-related potential (ERP) with its split-half noise ceiling, ordered by correlation. **f**, Latencies of the GFP peaks the model produces on its own, one row per participant in the order of **e.** Each peak is colored by the classical window it falls in within a ±30 ms tolerance, in two colors when it falls in two; the pale band is the tolerance and the darker band the classical window. **g**, Reaction-time quantile error (predicted minus observed) at the 0.1, 0.3, 0.5, 0.7, and 0.9 quantiles. **h**, Predicted against observed reaction-time quantiles (the same five quantiles) with the identity line. **i**, ERP correlation of each participant’s model on its own connectome and on a degree-preserving shuffle of it (Null-DTI), paired per participant; bars, means with bootstrapped 95% CIs; bracket, sign test. Own minus shuffled +0.881 (95% CI [+0.764, +1.001], 17/17, *P* = 1.5 ×10^−5^). The own column is the main fit and the shuffled one a refit on a shorter schedule, so the difference includes that training-budget gap; with both sides refit on the same schedule it is +0.832 ([+0.709, +0.959]; Supplementary Table 1).

Each person’s model was faithful to that person. For the best-fitting participant, sub-13 (ERP correlation *r* = 0.941, Fig. 2b–d top), it reproduced the scalp topographies at 90, 180, and 340 ms, the global field power (GFP), and the reaction-time distribution. For the worst, sub-01 (*r* = 0.632, Fig. 2b–d bottom), its N1 and P3 topographies reached only 0.28–0.63 of the observed amplitude, and its fit to the slow tail of reaction times was the cohort’s poorest. Across the 17 participants, the median ERP correlation was *r* = 0.858 (median ratio to the split-half noise ceiling 0.881), and all 17 exceeded *r* = 0.60 (Fig. 2e). Predicted reaction-time quantiles fell close to the identity line (median error +0.1 ms at the 0.9 quantile; Fig. 2g,h, Supplementary Fig. 1, and Supplementary Table 2). The task is nearly errorless (6.3% errors, with no reaction time recorded on error trials), so the model fits reaction time alone.

The model was not given the ERP components; its network dynamics produced them. Driven by nothing but the 200-ms pulse, 5 of 17 fitted networks produced a P1 peak, 9 an N1 peak, and 14 a P3-like late parietal peak, each within ±30 ms of its classical window (Fig. 2f). These 14, the participants with an emergent P3 peak, are the sample for the P3-window (300–600 ms) test of Fig. 6h.

The evoked response depended on the person’s wiring. We replaced each participant’s connectome with a degree-preserving shuffle (Null-DTI), in which every region keeps its number of connections but the specific wiring is destroyed, and refit the model. The ERP correlation fell in all 17 participants, by 0.881 on average (95% CI [0.764, 1.001], sign test *P* = 1.5× 10^−5^; Fig. 2i), and by 0.832 ([0.709, 0.959]) when both sides were refit on the same schedule (Supplementary Table 1).

### EEG-inferred latent variables predict held-out reaction times when behavior is in the fit

Within a stimulus condition, both the reaction time and the evoked EEG vary from trial to trial^9,10^. If one cause shapes both on a given trial, that trial’s EEG should carry information about its speed. The model has one such cause, the trial-specific latent variable *z*_*n*_: a single number per trial whose deviation from its condition’s mean is added to the input of the 10 decision-pool regions during the 200-ms sensory pulse alone (Eq. 16). It raises or lowers the pooled evidence, and with it the drift and the expected reaction time. The latent variable thus acts early in the trial; the model does not specify its source. It is the only trial-specific cause that the EEG and reaction time share, whereas accumulator noise acts on reaction time alone. EEG and reaction time are therefore independent once the latent variable is known, so any single-trial link between them must pass through it. The latent variable is not observed. An encoder network infers it from the trial’s whole EEG epoch, −200 to 800 ms, so the inference can draw on EEG from any part of the trial, including after the response. We test below whether EEG recorded earlier suffices.

Does the latent variable that the model infers from a trial’s EEG tell whether that trial will be fast or slow for its condition? We split each participant’s trials into five contiguous blocks, refit the whole model on four, and predicted the reaction times of the fifth from its EEG (Methods). Across the 17 participants, the model explained 9.6% of the within-condition reaction-time variance on held-out trials (95% CI [6.9, 12.2]; Fig. 3a). The explained share exceeded zero in 16 of the 17 participants (sign test *P* = 2.7 × 10^−4^; Fig. 3d). Predicted and observed within-condition residuals correlated positively in all 17 (mean *r* = 0.314, 95% CI [0.269, 0.358], sign test *P* = 1.5 × 10^−5^). A poor ERP fit did not prevent the prediction: the largest share, 17.4%, came from sub-01, the participant whose ERP the model fit worst (Fig. 2e), with predicted and observed residuals correlated at *r* = 0.451 (Fig. 3c).

**Figure 3:**
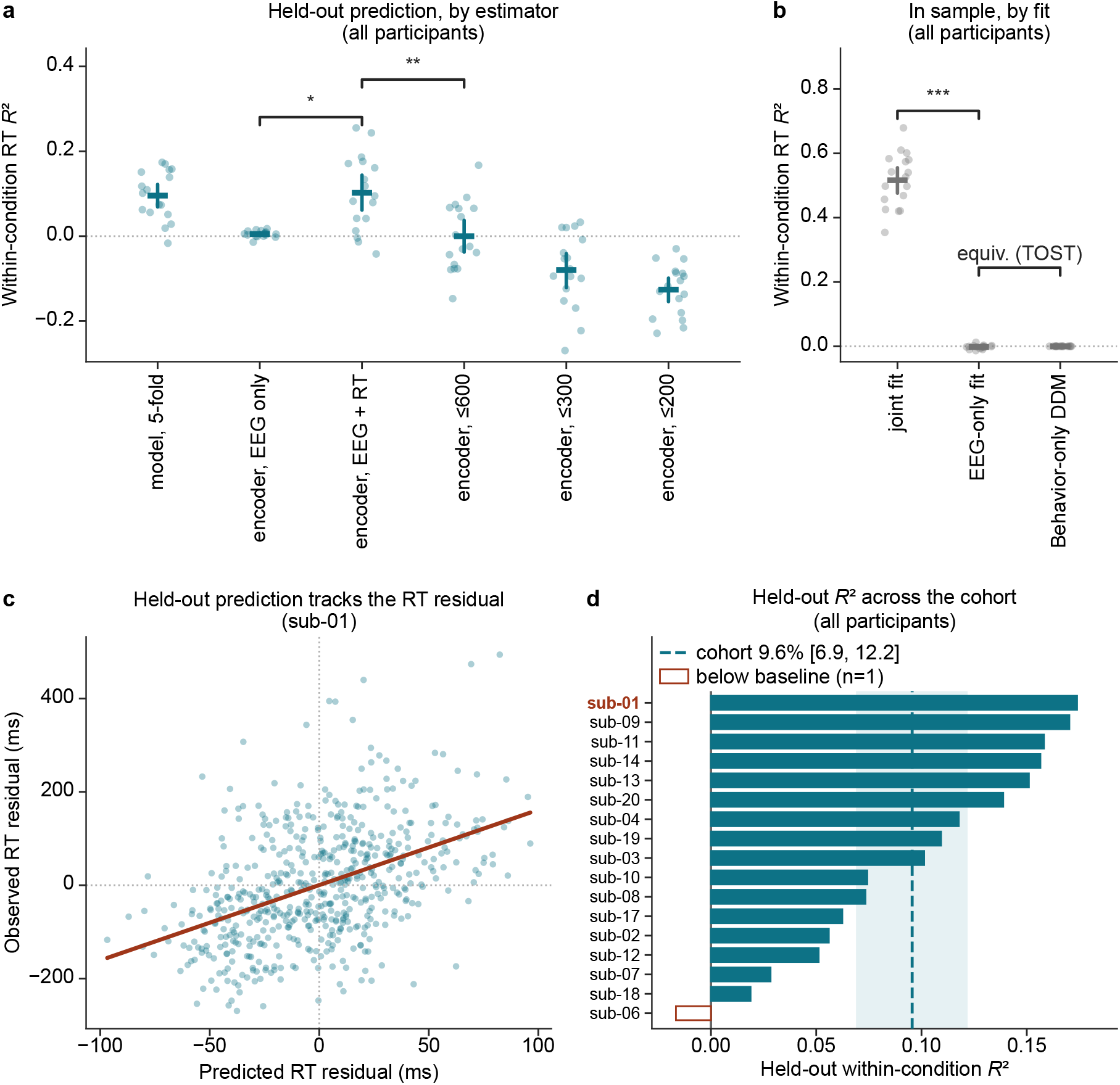
With behavior in the fit, EEG-inferred latent variables predict which held-out trials are fast or slow for their condition. Each estimate is the share of within-condition reaction-time variance explained, after each stimulus condition’s mean is removed; a prediction from the condition means alone scores zero, and a worse one scores below zero. Faint points, participants; bar, the cohort mean with its bootstrapped 95% CI; brackets, paired sign tests across the 17 participants, except the in-sample EEG-only and DDM pair of **b**, which carries a test of equivalence (two one-sided tests, ±2 percentage points). **a**, Held out. *Model, 5-fold*: the whole model refit without each fifth of the trials, so no scored trial entered the fit that predicts it (the estimator of **c** and **d**). *Encoder*: a recognition encoder refit within each fold, with the network and decision parameters of the fit to all trials (Methods); a scored trial’s reaction time never reaches the encoder that infers its latent variable but did enter those parameters. Each encoder was trained on the EEG alone (*EEG only*), on the EEG and reaction time (*EEG + RT*), or on the EEG and reaction time with the EEG cut at 600, 300, or 200 ms after stimulus onset. The median reaction time is 650 ms, so the whole-epoch columns read EEG recorded after the response on most trials (Supplementary Note 1). **b**, In sample, on the trials used for fitting; these values measure fit, not prediction. The joint model, the EEG-only model with its decision readout regressed on reaction time after fitting, and the behavior-only DDM. **c**, sub-01, the participant whose ERP the model fit worst (Fig. 2e) and the largest held-out share in the cohort: held-out predicted against observed reaction-time residuals (ms), each trial predicted by the fold that did not see it (*R*^2^ = 0.174, *r* = 0.451); line, least-squares fit. **d**, Held-out share explained in **a**’s first column, per participant; hollow, below the zero baseline. Dashed line and band, the cohort mean and its CI.

This prediction depended on fitting reaction time together with the EEG. On held-out trials, an encoder refit within each fold explained 10.2% of the within-condition variance when reaction time was in its objective (95% CI [6.2, 14.4]; Fig. 3a). Trained on the EEG alone, the same encoder explained 0.5% ([0.1, 0.8]), 9.7 percentage points less ([5.8, 13.7], 14 of 17, sign test *P* = 0.013; Supplementary Table 3). On the trials used for fitting, where the scores measure fit rather than prediction, the joint model explained 51.6% ([47.6, 55.5]), whereas a model fit to the EEG first, with its decision readout regressed on reaction time afterwards, explained no more than a DDM that never sees the brain (equivalent within ±2 percentage points, two one-sided tests, *P* = 5.5 × 10^−10^; Fig. 3b and Supplementary Table 1).

The encoder trained with reaction time drew this information from late in the trial, on most trials after the response. Given only the EEG up to 200, 300, or 600 ms after stimulus onset, each window including the 200 ms before onset, it explained −12.6%, −8.0%, and 0.0% of the held-out variance, none above the condition means alone (Fig. 3a). Given the whole epoch, to 800 ms, it explained 10.2%, 10.3 percentage points more than with the 600-ms window ([7.2, 13.2], 15 of 17, sign test *P* = 0.002; Supplementary Note 1). The median reaction time was 650 ms, and 81% of responses fell inside the epoch, so on most trials the whole epoch contains EEG recorded after the response. Two features read directly from the observed EEG, the centro-parietal amplitude and the projection on the condition-mean topography, explained −0.3% in the early window (80–200 ms; above a within-condition permutation null in 1 of 17 participants) and 2.0% in the P3 window (300–600 ms; 11 of 17; Supplementary Fig. 2b). The P3 rises to a fixed amplitude just before the response^29^, so the encoder may read activity timed by the response rather than activity that predicts it.

### The latent variable sets which trials are slow; accumulator noise sets the spread

What sets the variance that the latent variable left unpredicted on held-out trials (Fig. 3a)? In the model, a trial’s reaction time has two sources: an expected reaction time, which the latent variable sets through the drift, and accumulator noise, which is drawn afresh on every trial and which nothing in the trial’s EEG or condition predicts. Accumulator noise carried most of the main model’s own reaction-time variance: 89.5% against 10.2% for the latent variable, averaged over the 17 participants (latent variable 2.4–26.4% across participants; law-of-total-variance partition, Supplementary Fig. 3 and Supplementary Note 2). The stimulus condition carried the remaining 0.3%.

The latent variable sets a trial’s expected reaction time early in the trial. It enters with the 200-ms sensory pulse and raises or lowers the pooled evidence (Fig. 4a). The evidence sets the drift, and a higher drift brings the expected first-passage time forward (Fig. 4b). Trials separate during the pulse and converge soon after: in sub-13, the across-trial s.d. of the pooled evidence fell from 1.87 at 150 ms to 0.10 at 300 ms. The accumulator keeps the lead it gained during the pulse, so early activity can shift a much later response, a prediction that time-gated silencing tests below (Fig. 7b).

**Figure 4:**
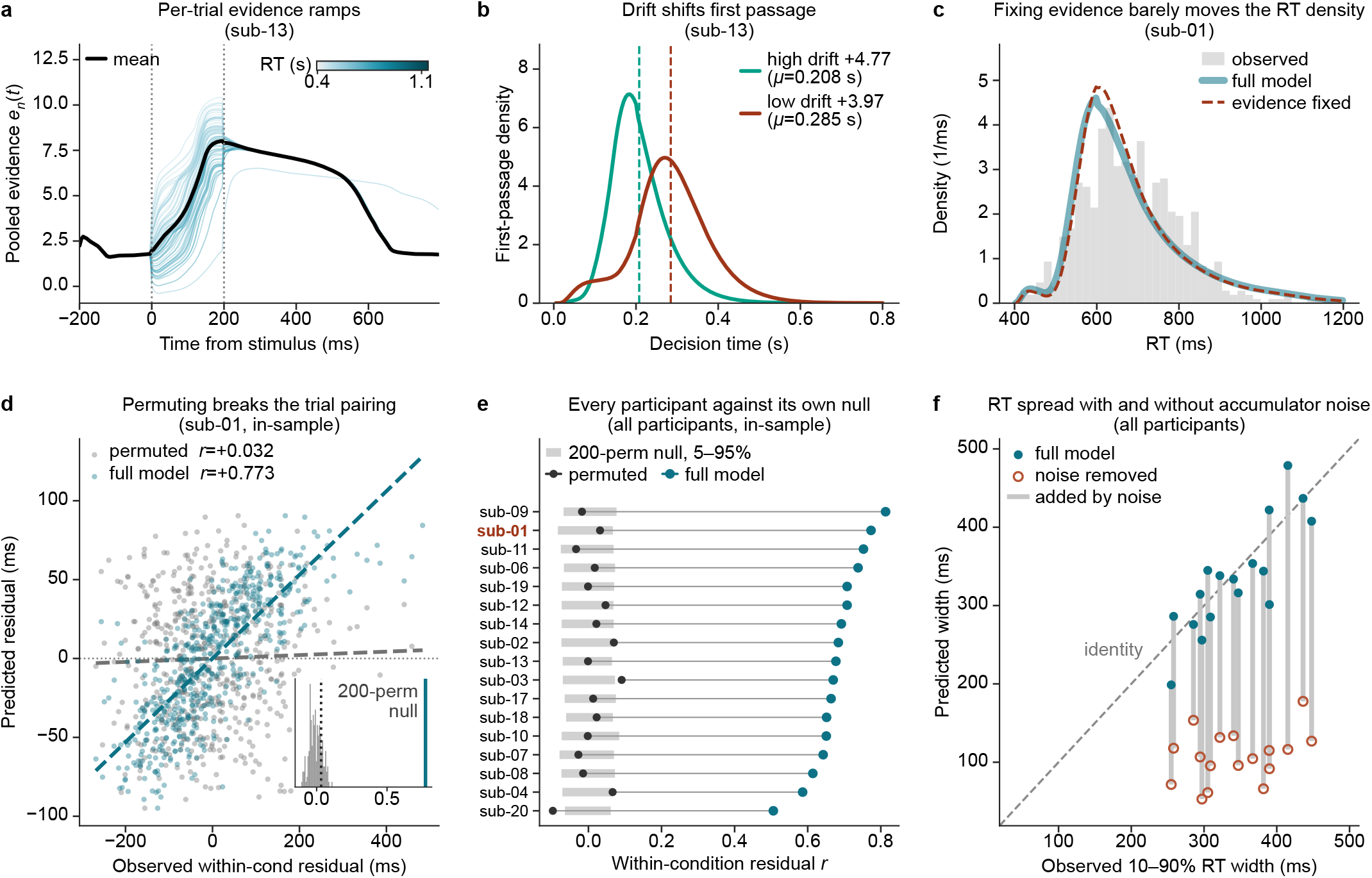
A trial’s latent variable shifts its expected reaction time; accumulator noise supplies most of the spread. **a**, Pooled evidence *e*_*n*_(*t*) of the decision pool for sub-13 from −200 to 795 ms: 70 of 581 trials colored by reaction time and the mean of all 581 in black; dotted lines bound the sensory pulse. **b**, The first-passage density at two drift rates (sub-13), 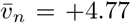 and +3.97, 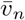 being a trial’s drift *v*_*n*_(*t*) (Eq. 21) averaged over 0–795 ms. Decision time is reaction time minus *t*_0_; dashed lines, the mean decision times (0.208 and 0.285 s). **c**, The predicted reaction-time density of sub-01 with the pooled evidence intact or fixed at its within-condition mean, over the observed histogram. Fixing moves it by a total-variation distance of 0.045; permuting the evidence across trials within condition returns the intact density by construction and is not drawn. **d**, Permuting breaks the trial pairing (sub-01): predicted against observed within-condition reaction-time residuals for the full model and one permutation, with fitted lines; inset, the null from 200 permutations. The decision layer was fitted on these reaction times, so this is an in-sample goodness of fit. The fixed arm is constant within a condition, so it has no correlation and is not drawn. **e**, The cohort version of **d**: each participant’s residual correlation for the full model and one permutation, against the band formed by its own 200 within-condition permutations (5th to 95th percentile); sub-01 in red. The full model lies above the band in 17 of 17 participants and the permuted model outside it in 3 of 17; the band covers 90% of permutations by construction, so about 1.7 are expected by chance. As in **d**, the correlations are in sample and the fixed arm is not drawn (Supplementary Table 4). **f**, Reaction-time spread, the width from the 0.1 to the 0.9 quantile, per participant: observed against predicted by the full model (filled circles) and by each trial’s expected reaction time, its conditional mean first-passage time plus *t*_0_, which keeps the drift and removes accumulator noise (hollow circles). The connector is the width that accumulator noise adds.

The latent variable sets which trials are slow, not how widely reaction times spread. We tested this without refitting. Replacing each trial’s pooled evidence with its within-condition mean removes the latent variable’s trial-to-trial effect, yet the predicted distribution barely moved (cohort median total-variation distance 0.043, where 0 is identical and 1 disjoint; Fig. 4c and Supplementary Table 4). Permuting the evidence across trials within a condition leaves the distribution unchanged by construction but breaks the pairing between each trial’s prediction and its observed reaction time. The correlation between predicted and observed within-condition residuals fell from 0.773 to 0.032 in sub-01 (Fig. 4d) and from a cohort median of 0.677 to 0.013. The full model’s correlation lay above the band of the participant’s own 200 permutations (5th to 95th percentile) in 17 of 17 participants (Fig. 4e and Supplementary Table 4). These correlations are in sample: the encoder was trained with these trials’ reaction times, so they measure fit, not prediction, and exceed the held-out correlation reported above (mean *r* = 0.314).

Accumulator noise sets most of the spread, measured as the width from the 10th to the 90th percentile of reaction time. The full model reproduced each participant’s width (median ratio to the observed width 0.97; Fig. 4f). When each trial instead took exactly its expected reaction time, which keeps the trial’s drift and removes accumulator noise, the width shrank in 17 of 17 participants, to a median 0.285 of the observed width.

### Where speed is set: shared sensorimotor and person-specific parietal effects

Which regions set a person’s speed, and are they the same in everyone? To find out, we silenced a region, re-ran the same trials, and recorded the change in mean predicted reaction time, ΔRT (Fig. 5a, right). Silencing cuts a region’s outgoing connections and, for a decision-pool member, drops it from the pooled evidence; the region itself stays active and still contributes to the EEG. In each participant we silenced each of the 83 regions, one at a time. We call one participant’s silencing of one region a cell of the map, and a cell is robust if its effect exceeds that participant’s noise band, the 95th percentile of the deviation over 100 re-simulations with fresh noise and no silencing. Of the 1,411 cells, 915 were robust (Fig. 5b), but most effects were small: the bands were 0.7–2.7 ms wide and the median robust effect was 4.8 ms (Supplementary Fig. 4a).

**Figure 5:**
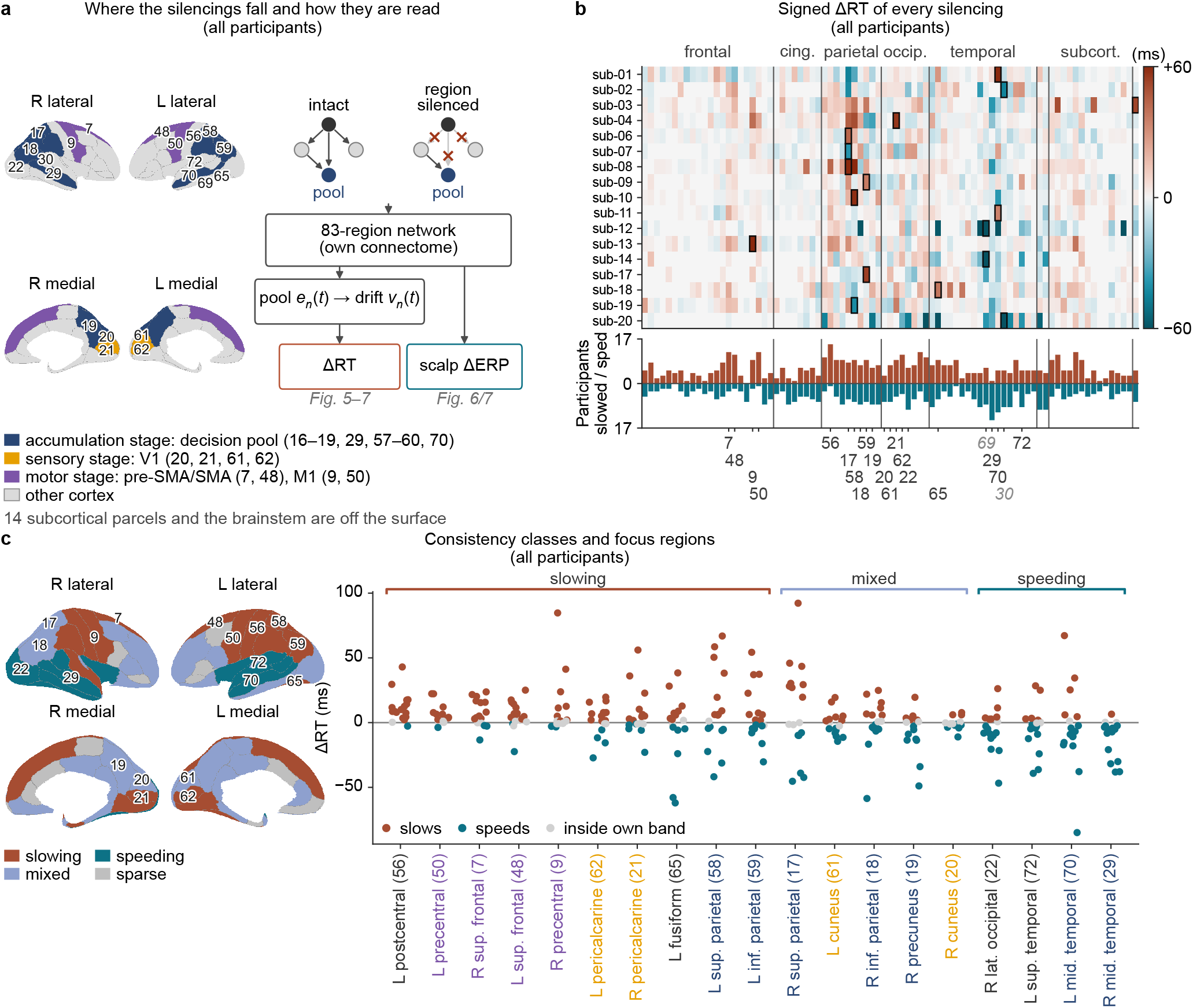
Which regions set a person’s decision speed, and whether the same ones in everyone. All panels are forward re-runs of the main fit on the same trials and seed. Regions are Lausanne-2018 scale-1 parcels, named with their index in parentheses (Supplementary Table 5). **a**, Left, three region sets on sub-13’s inflated surface, colored by stage: the four V1 injection sites, where the stimulus enters (sensory); the decision pool, whose evidence sets the drift (accumulation); and the pre-supplementary motor area (pre-SMA)/SMA (7, 48) and primary motor (9, 50) proxies (motor); index labels mark the focus regions of **c**. Right, the intervention: silencing a region zeroes its outgoing connections and drops it from the pool readout. It is read as the mean shift ΔRT of Eq. 31 and as the scalp change ΔERP (Figs. 6 and 7). **b**, Signed ΔRT (ms; positive slows) of every region silenced in every participant, regions in anatomical order, cells inside the participant’s own reseeded noise band blank, color clipped at ± 60 ms; the outlined cell is the participant’s strongest region. Below, the number of participants each region robustly slowed (up) or sped (down). **c**, Every region’s cohort class on the surface (slowing or speeding, robust in that direction in at least 9 of 17 participants; mixed, at least 3 in each direction; sparse otherwise) and, right, the 19 focus regions (Methods) in class order with every participant’s signed ΔRT (gray, inside the band), names colored by stage as in **a**.

**Figure 6:**
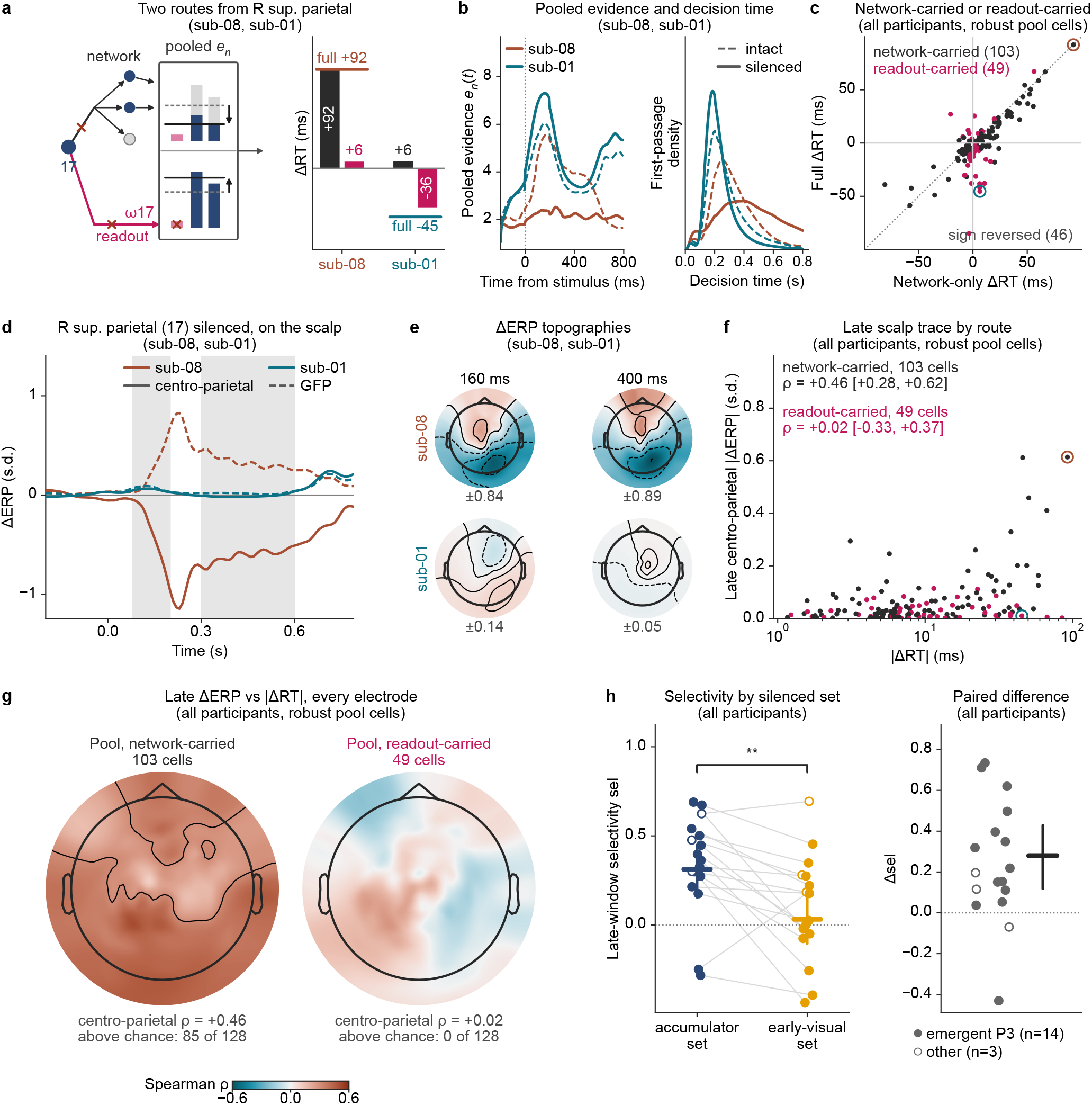
A silenced pool region reaches speed by two routes, and only effects carried by the network scale with the late EEG. All panels are forward re-runs of the main fit on the same trials and seed. Windows as in Fig. 2: early, 80–200 ms (P1–N1); late, 300–600 ms (P3). ΔERP is the change in the predicted condition-mean ERP, in units of the participant’s EEG s.d. (Methods), judged against the participant’s own reseeded band; the centro-parietal amplitude is its mean over nine electrodes around CPz and Pz. **a–e**, One silencing, two routes: the right superior parietal cortex (17) silenced in the two participants with the largest opposite effects of that silencing, sub-08 (+92 ms), which it slows through the network, and sub-01 (−45 ms), which it speeds through the readout. Figure 6: **a**, Left, the two routes from a silenced pool region to the reaction time, drawn for the right superior parietal cortex (17), each cut once (crosses): its outgoing connections, which drive the other pool members (network, black), and its fitted weight *ω*_17_ in the pool readout (readout, magenta; Eq. 32). Both routes end in the pooled evidence *e*_*n*_, the weighted mean (lines) of the members’ evidence (bars, region 17 in magenta), drawn intact (dashed) and silenced (solid) in one row per route: through the network (top), the other members’ evidence falls from its intact height (gray) and the mean falls with it; through the readout (bottom), region 17, whose evidence lies below the mean, leaves it and the mean rises. Right, each route alone for the pair, the full silencing as a line. **b**, The same silencing as dynamics for the pair: trial-mean pooled evidence *e*_*n*_(*t*) and first-passage density, intact (dashed) against silenced (solid). **c**, Every robust decision-pool cell in the cohort (152): full ΔRT against the network route alone, colored by the larger route, the pair ringed; a point whose two coordinates differ in sign is a reversal. **d**, Centro-parietal ΔERP (solid) and the global field power of ΔERP (GFP, the across-electrode s.d.; dashed) over time for the pair, windows shaded. **e**, The pair’s ΔERP topographies at 160 and 400 ms, all four maps on one color scale, each map’s own maximum printed. **f**, The 152 robust pool cells of **c**: late-window centro-parietal |ΔERP| against |ΔRT| , colored by the larger route, the pair ringed. Spearman *ρ* per route with 95% CIs resampling participants. Within participants, the network-carried *ρ* had a median of 0.57 (positive in 14 of the 16 participants with at least five such cells). **g**, The same correlation at every electrode, for the two classes of **f.** The contour on each map is its class’s chance maximum, the 95th percentile of the largest |*ρ*| over the 128 electrodes when |ΔRT| is permuted within participants (1,000 permutations). Beneath each map, the centro-parietal *ρ* and the number of electrodes above chance. **h**, The set-level test: late-window selectivity of Eq. 33 when the accumulator set is silenced against when the early-visual set is silenced, paired per participant, means with bootstrap 95% CIs over the 14 participants with an emergent P3 peak (filled; bracket, sign test). Right, the same pairing read as its per-participant difference Δsel, accumulator minus early-visual, with the mean and its bootstrap 95% CI over the same 14.

**Figure 7:**
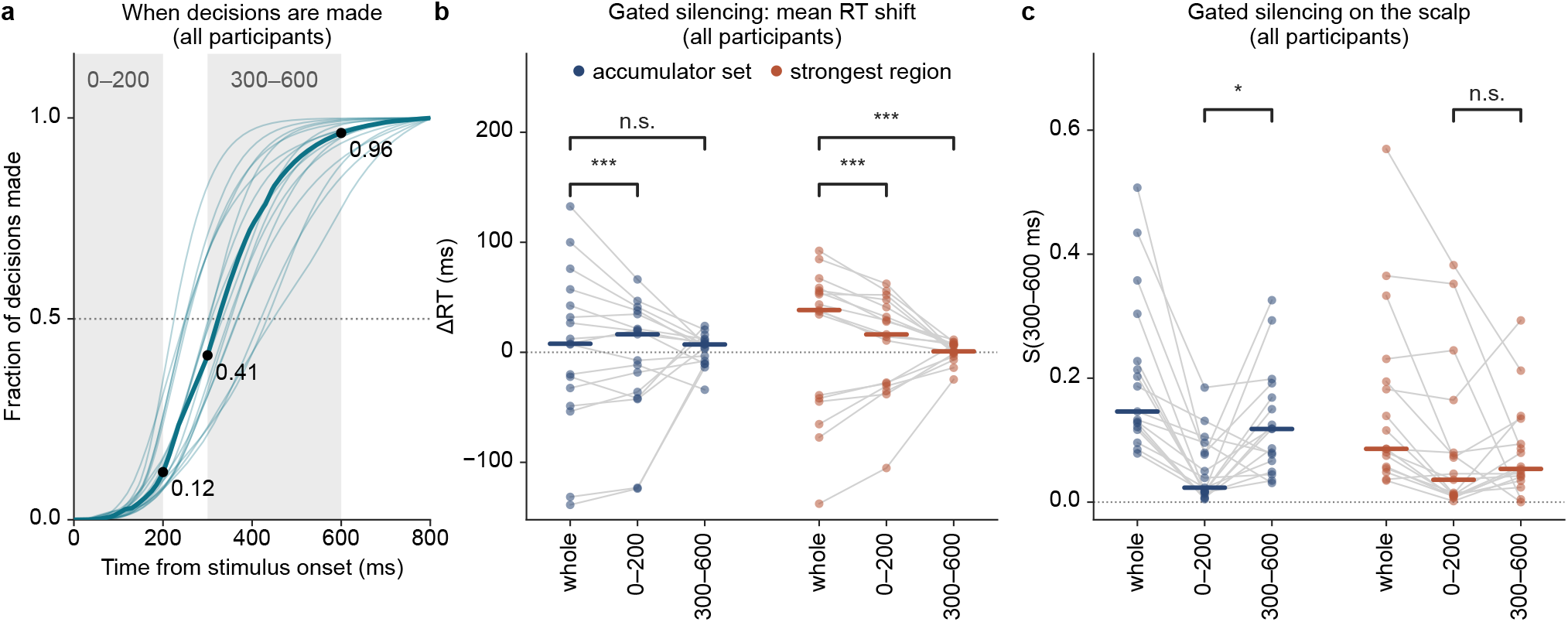
Silencing early reproduces the sign of each participant’s effect on speed; silencing late reaches the scalp. Windows and ΔERP as in Fig. 6. Panels **a** and **b** show when the speed is set, and **c** what reaches the scalp late. **a**, When the intact model’s decisions are made: the fraction of each participant’s decisions that have reached the bound, against time from stimulus onset (light teal, one line per participant; teal, the cohort median), with the two gates of **b** shaded. Dots, the median at the gates’ edges; at 300 ms it is 0.41, below the one half (dotted) fixed in advance as the point past which a small late effect would be uninformative. **b**, The silencing gated in time, for the parietal accumulator set and each participant’s strongest region: applied over the whole trial, during 0–200 ms only, or during 300–600 ms only, on the same trials, per participant with medians. ΔRT; brackets, two-sided sign tests: for 0–200 ms, of whether the gate keeps the whole-trial sign; for 300–600 ms, of whether its |ΔRT| is below a quarter of the whole-trial |ΔRT| . **c**, The same gates as in **b**, read on the scalp: the late-window scalp change *S*_300–600_, the across-electrode s.d. of ΔERP averaged over the window; brackets, the late gate against the early.

Some effects were shared across people. We called a region slowing or speeding when its silencing robustly moved reaction time in that direction in at least 9 of the 17 participants (Fig. 5c; all classes in Supplementary Table 6). The most consistent slowing region was the left postcentral gyrus (56), primary somatosensory cortex, which slowed 15 of 17 participants. Motor regions also slowed responses: silencing the pre-supplementary and supplementary motor area proxies (pre-SMA/SMA; 7, 48) slowed 12 and 11 participants, and silencing primary motor cortex (9, 50) slowed 9 and 12. The speeding regions were temporo-occipital: silencing the right middle temporal gyrus (29), the MT/V5 proxy, sped 14 participants and silencing its left homolog (70) sped 11.

Other effects were person-specific and opposite in sign. The right superior parietal cortex (17), a pool member, was among the eight strongest regions of 11 participants, more than any other region, yet silencing it slowed 8 and sped 6, from +92 ms in sub-08 to −45 ms in sub-01. Neighboring parietal and V1 regions split the same way, and ten different regions were a participant’s strongest region (Fig. 5b and Supplementary Table 7). Averaging over people therefore cancels effects that are large within each: the eight regions with the largest cohort-mean effect miss the strongest region of 7 of the 17 participants (Supplementary Fig. 4b,c).

A participant’s eight strongest regions were stable under trial resampling: 7 or 8 of the eight were retained over 200 resamples (Supplementary Fig. 5a). A participant’s model shared a mean of 1.71 of them with another participant’s model (136 pairs; 0.77 expected by chance) and 2.58 with its own refit on a degree-preserving rewiring of its connectome (five rewirings per participant; Supplementary Fig. 5b). Every entry of the maps is an in-model prediction that no real perturbation has tested.

### How speed is set: network and readout routes

Why would silencing the same region slow one person and speed another? A silenced pool region can change reaction time by two routes (Fig. 6a). Through the network, it stops driving other regions, other pool members among them, and their evidence changes. Through the readout, it leaves the weighted average that forms the pooled evidence, which is retaken over the remaining members; if the region carried less evidence than the average, the new average is higher, the drift faster, and the response earlier. Sub-08 and sub-01, chosen for this contrast, take different routes (Fig. 6a,b). In sub-08 the network route alone reproduces the full +92 ms, against +6 ms for the readout: the other members’ evidence falls, the peak of the pooled evidence drops from 5.6 to 2.5, and the distribution of decision times shifts later. In sub-01 the network route gives +6 ms and the readout −36 of the −45 ms: the region’s evidence lies below the pool’s weighted mean, so removing it raises the pooled peak from 6.0 to 7.3 and the distribution shifts earlier. The same arithmetic explains the speeding middle temporal regions: their network route alone slowed most participants, and their readout route sped them.

Both routes operate across the cohort. Of the 152 robust cells in which a pool region was silenced, the network route was the larger in 103 and the readout route in 49. In 46 of them (29 readout-larger, 17 network-larger), the full effect had the opposite sign to the network route alone (Fig. 6c). Reversals between people thus reflect how each fitted model weights its pool, not simulation noise, so predictions must be made per person.

The scalp signal most closely tied to evidence accumulation is the late centro-parietal positivity, over CPz and Pz in the P3 window (300–600 ms)^12,29^. Do the two routes leave different traces on the scalp in this window? By construction they should: dropping a region from the pooled evidence changes no region’s activity, so only the network route can alter the simulated EEG. From the same simulations we therefore read the change in the predicted condition-mean ERP, ΔERP, robust when it exceeds the participant’s own ERP noise band (Supplementary Fig. 2a). In sub-08, where silencing the right superior parietal cortex (17) acted through the network, the late centro-parietal amplitude (nine electrodes around CPz and Pz, averaged over the P3 window, 300–600 ms) fell by 0.61 s.d.; in sub-01, where the same silencing acted through the readout, it changed by −0.01 (Fig. 6d). Over the whole scalp, sub-01’s change in the P3 window was a twentieth of sub-08’s (GFP 0.017 against 0.33; Fig. 6d,e).

The cohort followed the same rule. Across the 152 robust pool cells, the late centro-parietal |ΔERP| grew with |ΔRT| in the 103 cells where the network route was the larger (Spearman *ρ* = 0.46, 95% CI resampling participants [0.28, 0.62]) but not in the 49 where the readout route was (*ρ* = 0.02, [− 0.33, 0.37]; network minus readout, Δ*ρ* = 0.44, [0.06, 0.82]; Fig. 6f). The two routes separated only in this window: in the early window (80–200 ms), |ΔERP| grew with |ΔRT| in both classes (network *ρ* = 0.45, [0.26, 0.63]; readout *ρ* = 0.37, [0.05, 0.65]). An effect on speed carried by the network leaves a late scalp trace that grows with its size; for one carried by the readout we detected none.

This trace was not confined to the nine centro-parietal electrodes. We compared the correlation at each of the 128 electrodes with a chance maximum, obtained by permuting the effects on speed within participants. Because it is built from the largest correlation over all 128 electrodes, it corrects for testing each one (Methods). In the 103 network-carried pool cells, the correlation exceeded it at 85 electrodes, and in the 49 readout-carried cells at none (Fig. 6g). An effect on speed carried by the network thus leaves a late trace across the scalp.

Does the late change single out the parietal accumulator? In the 14 participants with an emergent P3 peak, we compared silencing the parietal accumulator set, the eight parietal regions of the decision pool (bilateral superior parietal, inferior parietal, supramarginal, and precuneus cortex), with silencing early visual cortex (the four V1 injection sites) by window selectivity: the scalp change in the P3 window minus that in the P1–N1 window (80–200 ms), over their sum (Fig. 6h). Accumulator selectivity was +0.312 ([+0.155, +0.449], 12 of 14, *P* = 0.0129) and exceeded early-visual selectivity by Δsel = +0.280 ([+0.118, +0.429], 13 of 14, *P* = 0.0018). The selectivity is relative: silencing early visual cortex also changed the P3 window more than the P1–N1 window in most participants, only by less.

### When speed is set: early silencing carries the sign, late silencing reaches the scalp

When in the trial does activity set speed, and does the late scalp change drive it? We confined silencing to an early gate, 0–200 ms, when the latent variable is injected, or a late gate, 300–600 ms, the P3 window (Fig. 7a–c and Supplementary Table 8). Because a small late effect would be uninformative if most decisions were already made by 300 ms, we fixed in advance that the late gate would be read only if fewer than half had been made by then. That held: across participants, a median of 12% of decisions had been made by 200 ms, 41% by 300 ms, and 96% by 600 ms (Fig. 7a), so most were made while the late gate was open.

For speed, early silencing carried the sign. Silencing each participant’s strongest region during the early gate alone reproduced the sign of its whole-trial effect in 17 of 17 participants (sign test *P* = 1.5 × 10^−5^), though not its full size (median |ΔRT| 32 ms against 54 ms; Fig. 7b). Silencing it during the late gate alone gave a median of 6.9 ms, less than a quarter of the whole-trial effect in 16 of 17 (*P* = 2.7 × 10^−4^). For the parietal accumulator set, the early gate kept the sign in 16 of 17 participants (*P* = 2.7× 10^−4^), but the late gate’s effect fell below a quarter of the whole-trial effect in only 9 of 17 (*P* = 1.0; median| ΔRT| 42.4, 36.0, and 10.5 ms for the whole trial, the early gate, and the late gate).

On the scalp the order reversed. For the parietal accumulator set, the late scalp change *S*_300–600_, the across-electrode spread of ΔERP averaged over the P3 window, was larger under the late gate than under the early gate in 14 of 17 participants (medians 0.118 against 0.023, sign test *P* = 0.013); for the strongest region it was larger in 12 of 17, not significantly (0.054 against 0.036, *P* = 0.14; Fig. 7c). Late silencing of the strongest region thus barely changed speed, and late silencing of the accumulator set changed the late scalp signal more than early silencing did. In the fitted model, the late centro-parietal change is consistent with a trace of a decision whose speed was mostly set earlier.

## Discussion

Cognitive models describe the computational processes underlying behavior^22,23^, and whole-brain models characterize neural dynamics constrained by anatomical organization^15,16^. These two approaches have largely developed independently^21,34^, leaving unresolved a fundamental question: how do an individual’s brain dynamics shape that individual’s behavior across specific spatial and temporal scales? The BGM addresses this gap by constraining neural activity and behavior within a shared generative whole-brain model. A fitted model can then be used to ask where, how, and when neural dynamics set that person’s behavior.

The BGM changes what a whole-brain model can do. From The Virtual Brain^35^ and dynamic causal modeling^36^ to recent differentiable and simulation-based inference^37,38^, whole-brain models have mostly been asked to reproduce resting-state connectivity^17,18^ or evoked responses^14^, not behavior^21^. The BGM links whole-brain models to decision behavior. Our results show why this matters. Separate fitting of EEG and reaction times provided no advantage over a brain-blind DDM, whereas joint optimization with behavioral likelihood strengthened single-trial coupling both in-sample and on held-out trials. A whole-brain model that reproduces brain signals is therefore not necessarily a model of an individual brain that explains behavior.

Directly probing causal mechanisms in the human brain remains challenging, because available interventions have limited spatial coverage and temporal precision^39,40^. Fitted BGMs offer a complementary approach: they allow simulated perturbations of any region in any time window while preserving each individual’s anatomical constraints^20,41^. Three principles of how individual brain dynamics shape behavior follow. First, the regions that set decision speed differ between individuals, and population averages can obscure them^42^; a fitted BGM identifies candidate intervention targets for each person^43^. Second, a region’s effect depends on the route by which it reaches the decision, not on its anatomical name: the same region can slow one person through its influence on the network and speed another through its weight in the decision readout. Third, the dynamics that set behavior are not necessarily those most visible in EEG. In the fitted models, early activity largely set decision speed^44^, whereas late EEG^12,29^ mainly tracked the unfolding decision and reflected an effect on speed only when the network carried it.

Several limitations remain. First, the person-specific predictions come from a single exploratory cohort of 17 participants and need to be validated in larger cohorts. Second, the model mainly explains decision speed on correct trials and is limited in explaining the mechanisms that underlie errors. Third, the causal predictions from simulated perturbations will need to be tested with direct neural perturbation experiments.

Despite these limitations, the BGM provides a new paradigm for understanding how the brain produces behavior. In principle, it can extend beyond perceptual decision-making to other cognitive domains in which single-trial neural measurements are available, and beyond EEG to other recording modalities such as magnetoencephalography (MEG) and functional magnetic resonance imaging (fMRI). Fitted BGMs could also inform brain–computer interfaces (BCIs) that stimulate in closed loop^45,46^, for example by proposing, for each person, which region to target and in which time window.

## Methods

This section follows the order in which the model is built, fit, and interrogated. We describe the dataset, the frozen anatomical scaffold, and the differentiable forward model. Every participant is fit independently. The logistic function is written *φ*(*x*) = (1+*e*^−*x*^)^−1^. A symbol glossary and a data-flow schematic of the full model are given in Supplementary Note 3, Supplementary Table 9, and Supplementary Fig. 6.

### Dataset and task

We used the VEPCON dataset^27^ (OpenNeuro accession ds003505), in which observers reported whether a random-dot kinematogram, presented centrally in an 8^*°*^ circular field for 300 ms, contained coherent motion. Each trial carried one of three stimulus labels, coherent LEFT, coherent RIGHT, or NONE, on which all dots moved randomly, and per-trial behavior was that binary choice (report coherent vs. report random) together with a reaction time in seconds. Condition means and within-condition residuals throughout are taken over these three labels; coherent LEFT and RIGHT differ only in which hemisphere’s middle temporal cortex receives the motion drive of Eq. 16. Of 20 acquired participants we fit 18, each independently, excluding sub-05 and sub-15, whose motion-task EEG the dataset did not process because of excessive noise^27^. Of the 10,800 trials of the 18, the dataset marks 355 as epochs contaminated by artifacts and, of the rest, 984 as outliers, which include all 980 error trials; the remaining 9,461 trials are usable (9,245 without sub-16). One of the 18, sub-16, contributes 216 usable trials, below the *≥* 250-trial inclusion criterion (this participant inverted the response keys on several trials^27^), and is excluded from every cohort statistic and figure, which therefore report *N* = 17; its face-validity values appear in Supplementary Table 2 only (Supplementary Table 10). Per participant we used the preprocessed 128-channel EEG distributed with the dataset (downsampled to 250 Hz, average-referenced, and cleaned by independent component analysis^27^), the individual diffusion-MRI structural connectome, an EEG leadfield computed with MNE-Python^47^, and the Lausanne-2018 scale-1 parcellation^31,48^ (*R* = 83 cortical and subcortical parcels, named in Supplementary Table 5). Each epoch was cropped to [−200, +800] ms, baseline-corrected on the pre-stimulus interval, resampled to 100 Hz (*N*_obs_ = 100 samples), and divided by the participant’s global s.d. over trials, samples, and channels, so the EEG and ΔERP are in units of that s.d.

### Frozen anatomical scaffold

Three anatomical objects enter as fixed constants: the structural connectome *C*∈ ℝ^*R*×*R*^ (diffusion-MRI streamline counts, symmetrized and divided by the largest entry, so entries lie in [0, 1]), the mean fiber-length matrix *L* ∈ ℝ^*R*×*R*^ (symmetric, zero diagonal), and the leadfield **L** ∈ ℝ^*C*×*R*^ (unit Frobenius norm), with *C* = 128. The leadfield is a three-shell boundary-element forward model (ico-4 surfaces, conductivities 0.3, 0.006, and 0.3 S/m) on the participant’s FreeSurfer surfaces, with an oct-6 cortical source space of fixed surface-normal dipoles and the dataset’s electrode positions; the head-to-MRI transform is MNE-Python’s built-in fsaverage transform, not a per-participant coregistration. Each column is the mean over the dipoles of one parcel, so the 15 subcortical and brainstem parcels have zero columns and do not reach the EEG. Conduction delays are derived once from *L*,

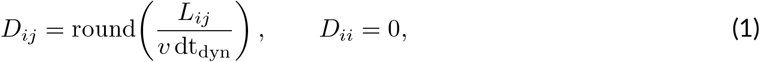

at a fixed conduction speed *v* = 3 m/s and dynamics step dt_dyn_ = 5 ms (*T*_dyn_ = 200 steps per epoch); microstructural estimates support treating conduction velocity as roughly constant across pathways, so that delay scales with fiber length^49^. None of *C, L*, **L** or *D* is updated during fitting. Nothing in the model fixes *R*: the parcellation enters only as the dimension of the state vector and the columns of the leadfield, so another Lausanne-2018 scale can be substituted^31^. Every fit reported here uses scale 1 (*R* = 83).

### Wilson–Cowan node dynamics and feedback-inhibition control

Each parcel *i* carries an excitatory state *E*_*i*_ and an inhibitory state *I*_*i*_ with net input currents

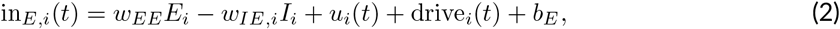

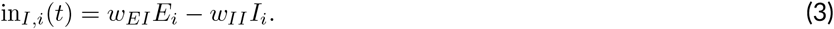

Each population relaxes toward its sigmoidal input–output response with a membrane time constant and additive current noise, so the pair obeys the Wilson–Cowan stochastic differential equations^30^

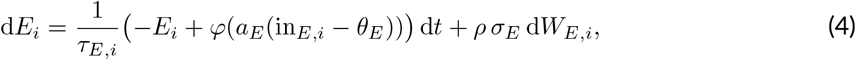

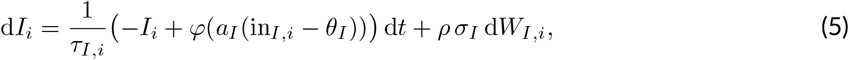

with independent standard Wiener processes *W*_*E,i*_, *W*_*I,i*_ and a noise switch *ρ ∈ {*0, 1*}* (*ρ* = 0 for the deterministic mean forward used in EEG prediction and analysis). Integrating Eqs. 4 and 5 with Euler– Maruyama on the dynamics step dt = dt_dyn_ (drift scaled by dt*/τ* , noise by 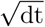 , so the scheme is step-size invariant) gives the discrete update

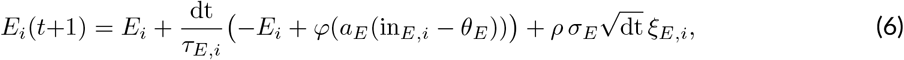

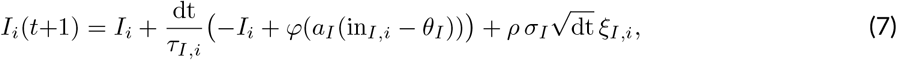

with *ξ*_*E,i*_, *ξ*_*I,i*_*∼ N* (0, 1) drawn per step. The net pyramidal current in_*E,i*_ of Eq. 2 serves as both the EEG source and the decision evidence. Constants were fixed at *w*_*EE*_, *w*_*EI*_, *w*_*II*_ = 1.5, 1.0, 0.5; *a*_*E*_, *θ*_*E*_, *a*_*I*_, *θ*_*I*_, *b*_*E*_ = 1.2, 2.8, 1.0, 1.0, 2.5; *τ*_*E*_, *τ*_*I*_ = 10, 15 ms; *σ*_*E*_, *σ*_*I*_ = 0.05; the tonic background *b*_*E*_ holds the baseline working point in the responsive range (in_*E*_ *≈ θ*_*E*_ *⇒E≈* 0.5). States are zero-initialized and reset at trial onset; all fixed constants are collected in Supplementary Table 11.

−The per-region inhibitory weight *w*_*IE,i*_ is set by offline feedback-inhibition control^50^ and then frozen, never entering the gradient. Calibration is a coupling-aware fixed-point iteration (600 steps, learning rate *η* = 0.05, *w*_*IE*_ clipped to [0.05, 10]) that drives each region’s in_*E,i*_ toward *θ*_*E*_ under the static coupling *W* = *G*_0_*C* (Eq. S2; Supplementary Note 4). At a fixed point the delayed coupling equals the instantaneous coupling 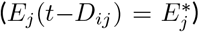, so calibrating on the delay-free system is valid for the delayed one. During training the working point is maintained softly by the penalty *R*_wp_ of Eq. 27; the full substrate, including *τ* , *σ* and *w*_*IE,i*_, is held frozen across the curriculum.

### Effective connectivity, neuromodulation and delayed coupling

In place of a single global coupling scalar, the connectome is reweighted with few degrees of freedom while remaining strictly on the SC support,

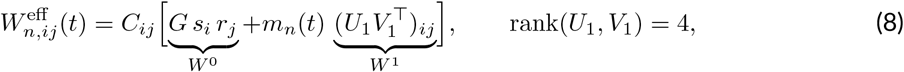

so both terms carry the factor *C*_*ij*_ and *W* ^eff^ vanishes wherever *C*_*ij*_ = 0 (no phantom edges). The neuromodulatory scalar *m*_*n*_(*t*) is a slow first-order filter of mean activity,

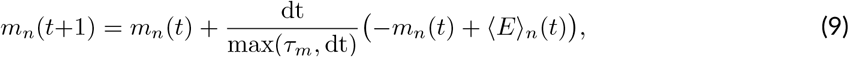

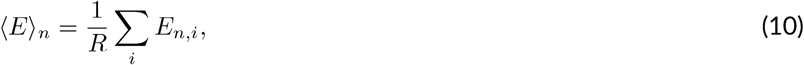

with *m*_*n*_(0) = 0. The delayed coupling current combines Eq. 8 with the delays of Eq. 1,

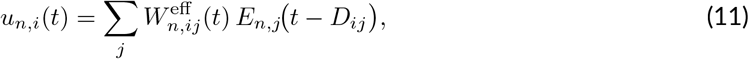

implemented as a differentiable shift buffer initialized at the calibrated set point (so the pre-stimulus interval carries no spurious transient; Supplementary Note 4). The learnable reweighting parameters *s, r, U*_1_, *V*_1_, *τ*_*m*_, *G* are fit during warm-start, frozen for the curriculum, and refit in the re-anchor step (Fitting protocol); *G*_0_ = 0.5 and *G* initialize at 0.5, *τ*_*m*_ at 20 ms, and *U*_1_, *V*_1_ at small random values. The regularizer *R*_ec_ of Eq. 28 anchors *W*^0^ to *G*_0_*C* and shrinks 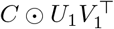.

### The trial-specific latent variable *z*_*n*_

The trial-specific latent variable *z*_*n*_, a single per-trial scalar, is the only trial-specific cause inferred from the data, and it is read from the trial’s EEG alone. Accumulator noise also varies from trial to trial, but it is integrated out in the first-passage density (Eq. 22) and reaches reaction time alone. It carries a standard normal prior *z*_*n*_ *∼ N*(0, 1), *z*_*n*_ ∈ ℝ^1^. The one-dimensional choice is deliberate: the latent is a single shared scalar cause, and because only the product *βλ* of drift gain and injection scale is identifiable from behavior, extra latent dimensions would buy degeneracy rather than explanatory power. The free-bits floor of Eq. 26 keeps even this single dimension from collapsing onto the prior. The latent is not a free parameter fit per trial but a reparameterized sample from an amortized recognition network^51^, so the prior is a variational regularizer, not a fitted distribution. The encoder of the main fit is linear and separable: it projects the single-trial EEG (*T, C*) onto three rank-one space–time patterns, each a channel vector times a time vector, and maps the three projections linearly to *µ*_*ϕ*_ and 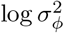 (the latter clipped to [−8, 4]), 692 parameters against the 2 × 10^5^ of a dense layer, so it reads every sample of the −200 to 800 ms epoch. The held-out encoder of Fig. 3a instead flattens the epoch, applies one linear layer and a tanh nonlinearity (width 16), and emits the same two heads. Either gives the posterior, its reparameterized sample and a closed-form KL term,

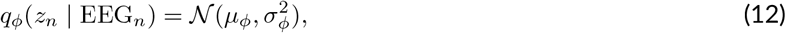

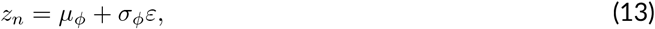

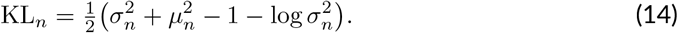

One encoder is shared across a participant’s trials. Its input is EEG only, but its parameters *ϕ* are trained under the joint objective of Eq. 25 in Stage C, which includes the behavioral likelihood, so the coupling between *z*_*n*_ and reaction time inside the main fit is a property of the fit and not a prediction. Out-of-sample prediction either reruns every stage within each cross-validation fold or, for the encoder estimator, refits the encoder alone on the training trials, first with the behavioral term off and then with it on (Held-out prediction and the forward problem), so the held-out trials never enter the encoder’s fit but the encoder is not behavior-blind. Under the encoder estimator, the network and decision parameters still come from the fit to all trials, held-out trials included. The EEG-only column of Fig. 3a keeps the behavioral term off in both passes, and the window-restricted columns multiply the EEG by a time mask before the encoder, so that only the samples up to 200, 300, or 600 ms after stimulus reach it; these refits ran with deterministic GPU operations. At inference we take the posterior mean *z*_*n*_ = *µ*_*n*_ and disable the dynamics noise.

The latent enters the network not as *z*_*n*_ but as its within-condition residual. With *c*(*n*) the condition of trial *n*,

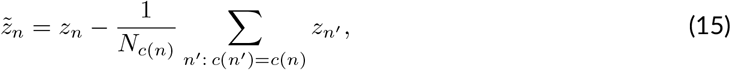

which is exactly zero-mean within every condition. The prior and the KL term of Eq. 14 act on *z*_*n*_ *∼ N*(0, 1); the decision pool receives 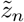 , which is centered by construction and so is not itself standard normal. These are two operations at two points, not a double-centering: the prior regularizes the per-trial posterior, while the residualization is a generative constraint that keeps the latent’s input zero-mean within every condition. The sensory channels of Eq. 16 are *z*-invariant, so the latent adds no condition-specific input, and a within-condition decode of reaction time from *z*_*n*_ cannot reflect the condition effect. The network is nonlinear, so the trial-to-trial spread of 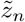 still shifts the condition-mean response; the predicted condition-mean ERP is therefore the mean of the trial-wise forward passes at the encoded latent variables (Fitting protocol).

### Task drive

The drive to parcel *i* on trial *n* is gated by a 200 ms sensory square wave *b*(*t*) = 1[0 *≤ τ* (*t*) *<* 200 ms],

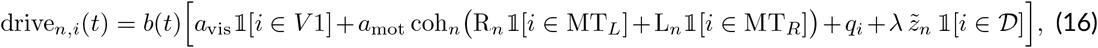

with a visual onset to bilateral V1 (pericalcarine, cuneus), a contralateral coherent-motion term to MT (middle temporal) on coherent trials only, where coh_*n*_ ∈ {0, 1}marks a coherent trial and R_*n*_, L_*n*_ ∈ {0, 1} its rightward or leftward direction, so a rightward stimulus drives left MT, a per-region distributed input field *q*_*i*_, and an additive decision-pool injection of the within-condition residual 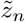 of Eq. 15. The sensory channels (*V* 1, MT, *q*) are invariant to *z*_*n*_, so the latent’s input reaches only the decision pool and carries no condition-specific offset. The drive levers *a*_vis_, *a*_mot_, *q* are fit during warm-start, frozen for the curriculum, and refit in the re-anchor step; the injection scale *λ* is fit in stages A and C. The dataset’s event files list a stimulus duration of 0.2 s, which sets the 200-ms pulse, whereas the dataset description reports 300 ms^27^. The stimulus was central, so the direction-to-hemisphere routing of the motion term is a modeling choice, not a retinotopic mapping.

### EEG observation model

EEG reflects slow dendritic post-synaptic potentials^52^, so the net pyramidal current is low-pass filtered before projection,

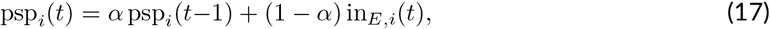

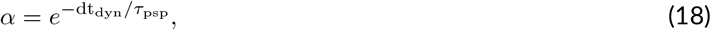

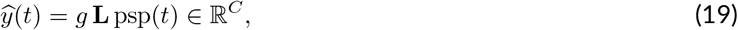

with *τ*_psp_ = 50 ms (frozen) and the PSP initialized at the resting current *θ*_*E*_ for a flat pre-stimulus baseline. Predictions are downsampled to 100 Hz and baseline-corrected per trial and channel by subtracting the mean of the first 20 pre-stimulus samples. The leadfield **L**, connectome *C* and lengths *L* are all frozen; the only free parameter on the observation side is the global gain *g*.

### Decision readout and first-passage likelihood

The decision pool *D* is a fixed scale-1 set of 10 parcels: bilateral MT/V5 together with bilateral superior parietal, inferior parietal, supramarginal and precuneus cortex. These are the scale-1 parcels covering the motion-sensitive and posterior parietal cortex in which motion evidence is encoded and accumulated^11,32,33^; the pool is a parcellation-level superset of those regions, not a set in which each parcel was individually named. The time-resolved pooled evidence is a convex combination of the *pre-sigmoid* net currents over the pool,

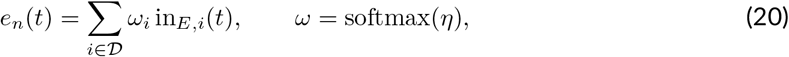

with readout logits *η* learned in the decision stages, initialized at zero so that the readout starts as the unweighted mean; the weights stay positive and sum to one, so the scale of the current is preserved. Using the current rather than the post-sigmoid rate avoids the sigmoid’s compression of trial-to-trial modulation. Because the drive injects 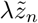 into *D* , the latent enters the evidence directly. The evidence of Eq. 20 sets a time-varying drift on a bounded accumulator through two gains, one on the within-condition mean of the evidence *ē*_*c*(*n*)_(*t*) and one on the residual around it, which is the part the latent moves,

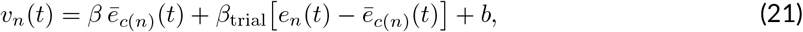

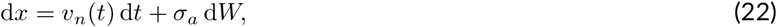

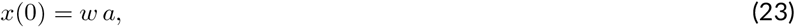

absorbed at *x≤* 0 or *x≥ a*. The accumulator starts at stimulus onset and runs to the end of the epoch, 0.8 s of decision time. Since a time-varying drift has no closed-form Wiener density, the first-passage likelihood is computed by a forward Fokker–Planck grid^53^: the interior (0, *a*) is discretized into *n*_grid_ = 128 points, each step applies a Gaussian transition kernel of mean *x* + *v*_*n*_(*t*) dt and variance 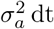, and mass crossing the boundaries is removed as flux by the mirror method; flux per unit time is the first-passage density (Supplementary Note 4). Every configuration of the main fit sets rt_marginal_likelihood=true, which sums the flux at the two boundaries before taking the logarithm. Boundary identity therefore carries no gradient, the likelihood is over reaction time alone, and the fitted model makes no accuracy prediction. The log-likelihood interpolates this marginalized density at rt_*n*_ − *t*_0_ with reaction time

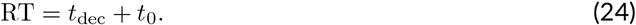

A trial whose rt_*n*_ − *t*_0_ falls beyond the 0.8-s grid (48 of the 9,245 trials of the 17 participants, 0.52%) receives a floor density of 10^−12^. The learnable decision parameters are *β, β*_trial_, *b, a, w, t*_0_ and the read-out logits *η*; the diffusion scale is fixed at *σ*_*a*_ = 1, the diffusion-model gauge, since first-passage times identify only *v/σ*_*a*_ and *a/σ*_*a*_^23^. The pass is fully differentiable to discretization and uses checkpointed scanning to bound memory. A closed-form constant-drift Wiener first-passage density^54^ is also implemented, but only as an optional warm-up approximation; every reported fit uses the time-varying-drift grid.

### Objective: negative ELBO

Each participant’s parameters maximize a single evidence lower bound, written as the loss

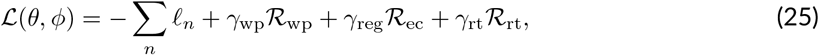

where *l*_*n*_ is the per-trial evidence-lower-bound term,

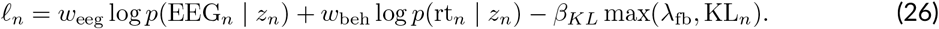

The EEG term is a per-trial Gaussian log-likelihood over the full single-trial waveform with learned observation scale *σ*_obs_ (irreducible variance absorbed into *σ*_obs_); the behavior term is the Fokker–Planck first-passage log-likelihood of the accumulator of Eq. 22. A free-bits floor^55^ *λ*_fb_ = 1.0 replaces the KL_*n*_ of Eq. 14 by max(*λ*_fb_, KL_*n*_), removing the gradient that would otherwise collapse the posterior to the prior, and is active in every stage. The remaining terms anchor the model to its priors,

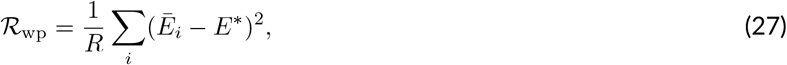

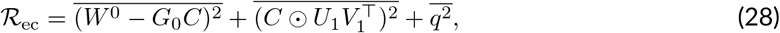

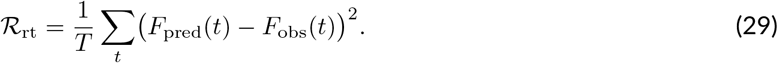

The working-point penalty *R*_wp_ holds mean excitation at *E*^*^ = 0.5; the effective-connectivity regularizer *R*_ec_ anchors the reweighting to the connectome; and the Cramér–von Mises reaction-time term *R*_rt_ pins the diffusion model’s heavy tail to the data (active in the decision stages, *γ*_rt_ = 100).

### Fitting protocol

Each participant is fit on a single GPU in four steps, with the parameter group trained at each step listed in Supplementary Table 11. First, feedback-inhibition control is calibrated and frozen by the fixed-point iteration of Eq. S2. Second, a warm-start places the model in an ERP-fitting basin before any latent or decision parameter is trained. Holding the latent at *z* = 0, it pre-fits only the topology levers (the set TOPO = {*s, r, U*_1_, *V*_1_, *τ*_*m*_, *q, G, a*_vis_, *a*_mot_}; Supplementary Note 4; *λ* is also in the optimizer set but receives no gradient while *z* = 0) to the condition-mean ERP by minimizing

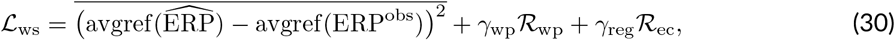

an average-referenced condition-mean ERP MSE under the same working-point and effective-connectivity anchors, Eqs. 27 and 28, as the curriculum, but at strong settings *γ*_wp_ = 0.10, *γ*_reg_ = 5 ×10^−3^ (weaker anchors let the effective connectivity diverge and the ERP correlation oscillate). Because *z* = 0 and condition determines the stimulus, every trial in a condition shares one forward pass, so only the unique conditions need be simulated, an ≈ 190× speedup; we run it at learning rate 10^−2^ for 5,000 epochs. Third, a three-stage curriculum runs with the topology levers and the entire substrate frozen:

- **Stage A (latent)**. Train the encoder *ϕ, λ*, the gain *g* and log *σ*_obs_ to single-trial EEG; weights (*w*_eeg_, *w*_beh_, *β*_*KL*_) = (1, 0, 0*→*1 annealed), regularizers (*γ*_wp_, *γ*_reg_, 0).
- **Stage B (decision)**. Train *β, b, a, w, t*_0_ and the readout logits *η* to the behavior likelihood, with *β*_trial_ held at its initial value until Stage C and *σ*_*a*_ fixed at 1 throughout; (0.1, 1, 1), (0, 0, *γ*_rt_).
- **Stage C (joint)**. Refine the latent/observation and decision parameters together under the full ELBO; (1, 1, 1), (*γ*_wp_, *γ*_reg_, *γ*_rt_).

The warm-start fits the response at *z* = 0, whereas the reported condition-mean ERP averages the responses at the encoded latent variables, and the nonlinear network passes their within-condition spread into that average. A fourth step of two short refits therefore follows the curriculum. The re-anchor refits the topology levers alone to the condition-mean ERP at the posterior-mean latent variable, under Eq. 30 with the same anchors, for 800 epochs at learning rate 10^−2^. The recalibration then refits the Stage B decision parameters under the full objective with *w*_eeg_ = 0.1 and *γ*_wp_ = *γ*_reg_ = 0, for 100 epochs at learning rate 2 × 10^−3^, so that reaction time is refit to the re-anchored dynamics.

The free-bits floor is on in every stage. Defaults were (300, 200, 200) epochs, curriculum learning rate 2 × 10^−3^, *γ*_rt_ = 100, *λ*_fb_ = 1.0, Adam^56^ with gradient-norm clipping at 1.0. Every quantity in the model, its constraint transform and its initial or fixed value are collected in Supplementary Table 11.

### Identifiability and synthetic-data recovery

Two degeneracies are known and shape what we claim. The drift scale *β* and the injection scale *λ* trade off, because only the product *βλ* scales the per-trial evidence; the trade-off is mitigated by pinning *λ* through its KL and EEG imprint and *β* through the behavior likelihood. And a binary stimulus with no recorded error trials makes the bound *a* and the non-decision time *t*_0_ only weakly separable, so we make (*a, t*_0_) claims within participants and not as cross-participant point estimates. Before interpreting any fitted parameter we ran a synthetic-data recovery test: trials generated from the frozen model were refit by the same training curriculum. The test used an earlier configuration of the model: a single drift gain *β* on the whole pooled evidence, a nonlinear encoder, a uniform pool readout, and a free diffusion scale *σ*_*a*_, not the configuration of the main fit. The test was run in two participants, sub-14 and sub-09, chosen before the test by the size of the latent variable’s forward imprint on the predicted ERP, each swept over an 11 × 11 grid of *β*× *λ* working points, giving 121 full refits per participant. Two criteria were fixed before the test was run: latent recovery, the correlation between the generating and the recovered *z*_*n*_, against a 0.20 threshold; and *β*/*λ* separability, by bootstrap partial correlation of each recovered gain with its generating value. Each criterion was then repeated under ten fresh master seeds. The recovered values, the reseeding, the log–log recovery slope that the partial correlation cannot see, and the recovery of the decision parameters are set out in Supplementary Note 5 and Supplementary Table 12.

### Face-validity checks

Behavioral claims rest on face-validity checks computed per participant and fixed before they were run. The primary check (a) is the element-wise Pearson correlation between predicted and observed condition-mean ERP (average-referenced, pooled over condition, time and channel), required to exceed 0.60. Because that correlation is bounded by how much of a participant’s own ERP is reproducible at all, we also compute a split-half noise ceiling from that participant’s trials and report the model correlation as a fraction of it (Supplementary Table 2). The ceiling averages the half-to-half correlation over 20 random within-condition splits at a fixed seed, each half average-referenced and pooled over condition, time and channel, then rescales it to full-sample reliability *ρ* by the Spearman–Brown correction 2*r/*(1 + *r*), and takes the ceiling as 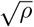 , the expected correlation of a perfect model with the noisy data. The check was evaluated on all 18 fitted participants; every cohort statistic below is over the 17 retained. A secondary diagnostic compares single-trial EEG *r*^2^ against the condition-mean baseline; the cohort median is −0.047, so the model does not beat that baseline trial by trial. Check (b) compares reaction-time quantiles (0.1–0.9) for shape and tail agreement; check (c) looks for emergent global-field-power peaks not templated into the model, scored separately in the fixed P1 (80–130 ms), N1 (150–200 ms), and P3 (300–600 ms) component windows with a ±30 ms tolerance. Each window is tested independently, so a peak landing inside two tolerance-widened windows counts for both, and the per-window hit counts can sum to more than the number of participants. The same window definitions are reused by the stage-signature and counterfactual-EEG analyses, the latter pooling P1 and N1 into one early window (80–200 ms) against the late P3 window (300–600 ms); the participants with an emergent P3 peak define its analysis subset. Per-participant outcomes across the cohort are shown in Supplementary Fig. 1. The within-condition decode (d) is a block-cross-validated linear decode of reaction time from the latent variable: both *z*_*n*_ and reaction time have their training-fold condition means removed, so a positive held-out *R*^2^ cannot reflect the condition effect. This residualization matches the condition-residual injection of Eq. 16 and is the operational definition of the single-trial link.

### Constraint and joint-fit tests

We tested the individual connectome with a degree-preserving (Null-DTI) shuffle^57^ that preserves the degree sequence but destroys the specific wiring, holding the leadfield, fiber lengths, injection sites, and decision pool fixed. The topology levers and the EEG observation parameters were copied from an own-connectome refit and frozen, and only the encoder, the injection scale *λ*, and the decision parameters were refit, on five shuffles per participant. We report own-minus-shuffle differences in ERP correlation and in within-condition coupling *z*_within_, with participant-level confidence intervals and minimum detectable effects. The joint objective was tested against an EEG-only model, refit with the behavioral weight set to zero and its decision readout then regressed on reaction time under the same protocol, and against a behavior-only DDM baseline, both scored by *z*_within_: a linear decode of the trial’s latent variable to reaction time over five contiguous cross-validation blocks, with each fold’s per-condition means estimated on its training trials and subtracted from both. The decode is held out but the encoder is not, since it comes from the fitted model and that fit saw reaction times, so *z*_within_ measures coupling inside the fitted model. Out-of-sample prediction is tested separately, by refitting within each fold (Fig. 3a,c,d). The test has two parts: the joint model must exceed the post-hoc readout by a sign test, and the post-hoc readout must be shown *equivalent* to the baseline by two one-sided tests against a ± 0.02 bound fixed in advance, so that the second part is an active demonstration rather than a failure to reject (Supplementary Note 2; Supplementary Table 1).

### Held-out prediction and the forward problem

A model can match the condition-mean ERP and the reaction-time distribution without knowing which trial is which, so the link between a trial’s latent variable and its reaction time is tested on held-out trials. Held-out prediction is scored first on the whole pipeline. Each participant’s trials were split into five contiguous blocks, every stage of the fit was rerun on four, and the model predicted the reaction times of the fifth. Every condition mean the prediction uses came from the training blocks: the mean latent variable that *z*_*n*_ is residualized against, the mean pooled evidence that sets the condition part of the drift in Eq. 21, and the mean observed and predicted reaction times the residuals are taken against. The five held-out blocks were pooled, so every trial is scored once, and the score is the out-of-sample *R*^2^ = 1−Σ (*y* − *ŷ*)^2^/Σ *y*^2^ on those residuals with the slope fixed at 1: a prediction that knows only the training-block condition means scores zero, and a worse one scores below zero. We also report the Pearson correlation *r* between predicted and observed residuals, which tests the direction of the prediction but not its scale. Contiguous blocks make the score conservative, since slow drift across the session separates training from test trials.

The encoder estimator regresses reaction time on the EEG-inferred latent variable within condition under the same blocks. The EEG-only and window-restricted variants were run with this estimator only; the whole-model refit was not repeated for them. Inside each fold it refits only the encoder, the injection scale *λ* and the trial drift gain on the training trials, with the behavioral likelihood in its objective, so that a held-out trial’s reaction time never reaches the encoder producing its latent variable; the network and the decision parameters (bound, drift gain, *t*_0_) are those of the fit to all trials. The condition means subtracted from the state and from reaction time are estimated on the training trials of the fold, and the permutation null shuffles trials within condition. This encoder has a two-dimensional recognition latent, one component of which is a slow state carried from trial to trial as a first-order autoregressive process, so unlike the main fit’s *z*_*n*_ its state also reads the EEG of preceding trials. The ablations and the variance partition below read the main fit’s one-dimensional *z*_*n*_ in sample. The two latents are reported side by side and no quantity is carried between them (the encoder columns of Fig. 3a against Fig. 4). Two ablations then ask what the latent does. *Fixed-z* replaces the per-trial decision evidence with its within-condition mean; *within-condition permutation* keeps every trial’s evidence and shuffles only its assignment to trials. Both run one forward Fokker–Planck pass on the already-fitted model with no re-training, and each is scored twice, by total-variation distance between predicted reaction-time densities and by the correlation between predicted and observed within-condition residuals. These ablations are descriptive: no criterion was fixed in advance for them, and they produce no pass or fail. For the forward problem we partitioned each participant’s single-trial reaction-time variance into stimulus, recoverable latent-variable (*z*_*n*_) and irreducible diffusion components by the total-variance law, reporting shares rather than absolute variances because the diffusion term inherits a first-passage grid truncation bias; share robustness to that truncation is judged separately. We also correlated the fitted decision parameters (*a, t*_0_, *β, b, w*) and the decision-pool quantities with reaction-time summaries between participants; that table is provided with the analysis outputs (Data availability).

### Sensitivity maps

Sensitivity maps are computed by *in silico* silencing of the fitted network, the counterfactual-lesion approach that connectome-based whole-brain models allow^41^, implemented as output disconnection: silencing region *i* sets every entry of column *i* of the structural connectome to zero, self-coupling included, so the region keeps evolving under its own input and keeps projecting to the scalp but drives no other region; a decision-pool region is also dropped from the pool readout. “Silencing” throughout means this output disconnection plus the readout drop. Each of the 83 regions is silenced in turn, with reaction time re-simulated on the same trials and seed to give a per-target ΔRT, the change in the trial-mean predicted reaction time,

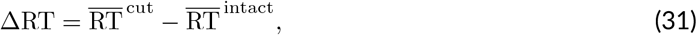

with the overline the mean over that participant’s trials. The pool readout under a silencing is the fitted-weight average of the remaining pool regions,

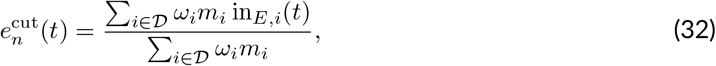

with *ω* the fitted readout weights of Eq. 20 and *m*_*i*_ ∈ {0, 1} the keep mask; it reduces to the intact readout when nothing is silenced. Because the denominator renormalizes the remaining weights to one, a silenced pool region leaves the average rather than contributing zero, so a region whose evidence lies below the weighted mean raises the pooled evidence when it leaves: the readout route of Fig. 6a. Every number in this paper is from the scan under these fitted weights. For decision-pool regions the full silencing is decomposed, without further forward runs, into the network route alone (column zeroed, readout kept) and the readout drop alone (intact forward, region removed from the readout); a region outside the pool has no readout weight, so its full silencing is its network route.

A target counts, and a silencing is robust, only when its |ΔRT| exceeds that participant’s own noise band, the 95th percentile of the absolute deviation from the median of the same readout over 100 re-simulations with fresh noise and no silencing; the existence criterion, fixed in advance, asks for at least one region or connection target (a single directed connection removed by zeroing its entry) beyond the band, and all 17 participants had region targets (Supplementary Table 7). Target stability is the inclusion frequency, the fraction of *B* = 200 resamples that keep a target among the participant’s eight strongest, scored over a frozen top-8 so that no participant’s count can exceed eight. The stability criterion, fixed in advance, counts participants: the number retaining a target at inclusion frequency ≥ 0.8 must exceed a frozen threshold of 8.5, more than half of the 17. A region’s cohort class is slowing or speeding when its silencing moved at least 9 of 17 participants robustly in that direction, mixed when at least 3 moved robustly in each direction, and sparse otherwise. The focus set comprises every region in at least four participants’ eight strongest regions together with the injection sites and the MT/V5, superior parietal, pre-SMA/SMA, and primary motor pairs, and the mechanism pair is the mixed decision-pool region in most top-8 sets with its largest robust slowing and largest robust speeding participant. We report each participant’s strongest region, the region of largest |ΔRT| (Supplementary Table 7) and cohort tables by region and for the focus regions (Supplementary Tables 6 and 13).

### Counterfactual EEG

The counterfactual-EEG test silences a set of regions in the forward model and reads the resulting ΔERP rather than ΔRT, summarized by the window selectivity

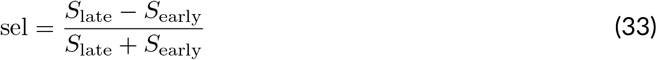

over the frozen P1–N1 and P3 windows, for the parietal accumulator (the decision pool with MT removed) and for early visual cortex (the bilateral V1 injection sites of Eq. 16, pericalcarine and cuneus cortex).

The paired readout of Fig. 6 takes the behavioral and the scalp effect of each region’s silencing from one forward run per region, on the same trials and seed. *S*_*W*_ is the time mean over window *W* of the across-channel standard deviation of ΔERP, and the scalp magnitude is the post-stimulus mean of that standard deviation, judged against the participant’s reseeded ERP band; the signed centro-parietal amplitude is the window mean of ΔERP averaged over nine electrodes fixed in advance around CPz and Pz (A2, A3, A4, A5, A19, A20, A32, B2, and D16 of the 128-channel montage). A cell is one participant’s silencing of one region. Across cells, |ΔRT| is related to the late-window| ΔERP| by Spearman correlation, with 95% CIs resampling participants and keeping all of a drawn participant’s cells (10^4^ draws). Per electrode, the chance maximum of a class is the 95th percentile, over 1,000 permutations of| ΔRT| within each participant, of the largest |*ρ*| over the 128 electrodes, whose positions are those of the cohort-mean montage on which the centro-parietal electrodes were fixed. The observed-EEG feature decode (Supplementary Fig. 2b) uses the folds and scoring of the held-out decode of Fig. 3, with two features per trial and window, the centro-parietal amplitude and the projection of the window-mean topography onto the training-fold condition-mean topography, centered within condition on the training fold and fit by ordinary least squares; its null shuffles reaction time within condition on the training folds (1,000 shuffles). These readouts are descriptive, and no criterion was fixed in advance for them.

### Time-gated silencing

The time-gated silencing replaces the connectome on the dynamics grid by *C* + *g*(*t*) (*C*_cut_ −*C*) with *g*(*t*) ∈ {0, 1} and gates the readout mask in the same way; the gates are [0, 200) ms, [300, 600) ms, and the whole trial, and an all-ones gate reproduces the whole-trial silencing exactly. The fraction of decisions made by time *t* is the intact model’s per-trial first-passage density integrated up to *t* of decision time, the absorption time minus *t*_0_, summarized by its median over trials; the accumulator starts at stimulus onset, so decision time runs on the gates’ clock, and the value at 300 ms is the condition fixed in advance for reading the late gate. These readouts are descriptive; apart from that condition, no criterion was fixed in advance for them.

### Statistical analysis

The criteria were fixed before the analyses were run, but the planned held-out confirmation was not carried out, so every *P* reported here is exploratory and none is corrected for the number of tests in the chain. These criteria were written in internal analysis plans and were not publicly registered. Cross-participant comparisons are two-tailed at *α* = 0.05. Sign tests are always two-sided, and a result counts as significant only when the two-sided *P* is below 0.05 *and* the majority of participants move in the predicted direction; choosing a one-sided alternative would halve every reported *P* , so all reported sign tests call the same routine. Where a paired contrast was fixed in advance as a sign test and the ranks carry information the counts do not, we report a Wilcoxon signed-rank *P* beside it. In figures a bracket marks a paired test between the two columns it spans, labeled * for *P <* 0.05, ** for *P <* 0.01, ***for *P <* 0.001, and n.s. for *P* ≥ 0.05; equiv. marks an equivalence test (two one-sided tests) that passed. Cohort summaries are reported as a mean or a median with a bootstrapped 95% confidence interval, the two never mixed within an analysis; those intervals are bootstrapped at the participant level (10^4^ iterations, fixed seed) unless noted as trial-level. The resampling unit is the cohort of the analysis in question: *N* = 18 for the face-validity checks and *N* = 17 for every cohort statistic and figure, sub-16 having too few usable trials to enter them, and *N* = 14 for the counterfactual-EEG test, whose subset is the participants with an emergent P3 peak. Within-condition and within-participant effects use permutation nulls that respect condition structure. For every null result we report the minimum detectable effect, (*z*_1−*α/*2_ + *z*_0.8_) s.d.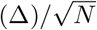 for paired differences Δ at *α* = 0.05 and 80% power. Where no smallest effect size of interest was set in advance, a negative result is not read as evidence of absence, and the minimum detectable effect beside it is descriptive.

## Data availability

The VEPCON dataset analyzed in this study is publicly available from OpenNeuro under accession code ds003505^27^. The per-participant anatomical inputs that fix the model scaffold are derived from that dataset by the preprocessing pipeline (see Code availability): the diffusion-MRI structural connectome, the EEG forward leadfield, and the inter-regional fiber-length delays in the Lausanne-2018 scale-1 parcellation^31^. Source data for the main and supplementary figures will be released with the code upon acceptance of the manuscript.

## Code availability

The model was implemented in Python with the BrainX ecosystem^58–62^. The model code, the fitting and analysis pipelines, the synthetic-data generator used for the identifiability test and the trained per-participant models will be made publicly available upon acceptance of the manuscript.

## Acknowledgements

We thank Professor Chi-Chung Hui (University of Toronto) for the selfless supervision and in-depth discussion. We thank Xinzhu Liao (Guangdong Institute of Intelligence Science and Technology) for help with visualization. This work was supported by the General Program of the National Natural Science Foundation of China (No. 32671468, CMW) and the Young Scientists Fund of the National Natural Science Foundation of China (No. 32400937, CMW).

## Author contributions

Conceptualization, methodology, analysis, visualization, writing — original draft: CMW. Writing — review and editing: CMW, GCY, SNW, and HL.

