## Supplemental Information for "A brain–behavior generative model links individual brain dynamics to perceptual decisions"


#### Contents


**1 Held-out reaction-time prediction by EEG window** **S3**
1.1 Motivation. . . . . S3
1.2 Encoder and protocol. . . . . S3
1.3 Results by window. . . . . S3
1.4 Effect of the training objective. . . . . S3
1.5 Interpretation. . . . . S3

**2 Variance budget, baselines, and analysis provenance** **S4**
2.1 The main model's variance budget . . . . . S4
2.2 Baseline and null models . . . . . S4
2.3 Analysis provenance . . . . . S5

**3 Notation, model overview, and data flow** **S5**
3.1 Model overview and data flow . . . . . S5

**4 Forward-model numerics and parameters** **S6**
4.1 Feedback-inhibition control (FIC) calibration. . . . . S6
4.2 Fokker–Planck first-passage solver. . . . . S6
4.3 Differentiable conduction-delay buffer. . . . . S7
4.4 Parameters: fixed, calibrated, and learned . . . . . S7

**5 Identifiability and synthetic-data recovery** **S7**
5.1 Degeneracies tested. . . . . S7
5.2 Test and outcome. . . . . S7
5.3 Reseeding. . . . . S7
5.4 Partial correlation and recovery slope. . . . . S8

6 Supplementary Tables S9

7 Supplementary Figures S26

Supplementary References S34

### 1 Held-out reaction-time prediction by EEG window

The main text reports only the outcome of this test (EEG-inferred latent variables predict held-out reaction times when behavior is in the fit); this note gives the encoder, the windows it was allowed to see, the numbers behind Fig. 3a and Supplementary Fig. 2b, and what the outcome does and does not show.

#### 1.1 Motivation.

Each trial's EEG epoch runs from $-200$ to $800$ ms around the stimulus, 200 steps of $5$ ms (Supplementary Table 11). The median reaction time is $650$ ms over $9{,}245$ trials, and $81\%$ of trials end before the epoch does, $55$–$96\%$ per participant. A reader given the whole epoch therefore sees EEG recorded after the response on most trials, so it could be reading the response rather than anything that preceded it. Cutting the window back is the way to find out.

#### 1.2 Encoder and protocol.

The reader is the joint encoder itself, refit within each fold with reaction time in its objective, which is the *encoder* estimator of Fig. 3a, with its input cut at $600$, $300$, or $200$ ms. It uses the five contiguous folds and the within-condition residualization of the held-out test (Methods Held-out prediction and the forward problem): each fold takes its condition means from its own training trials, so a negative held-out $R^2$ means the encoder does worse than predicting the training-fold condition mean. Every window starts at the start of the epoch; only its end moves.

#### 1.3 Results by window.

Supplementary Table 3 gives the ladder. The joint encoder falls below zero in the short windows, to $-12.6\%$ at $200$ ms, and reaches $+10.2\%$ on the whole epoch, beating its own $600$-ms window by $10.3$ points. Two hand-built readouts of two features each, centro-parietal amplitude and the projection on the training-fold condition-mean topography, agree (Supplementary Fig. 2b). The early window, $80$–$200$ ms, gives $-0.3\%$ and passes the permutation null in 1 of 17 participants; the P3 window, $300$–$600$ ms, gives $2.0\%$, passes in 11 of 17, and carries the negative slope in 16 of 17; the two windows together give $2.4\%$.

#### 1.4 Effect of the training objective.

The window is not the only thing that decides whether a reader works. Refit on the whole epoch but without reaction time in its objective, the encoder falls to $0.5\%$ ($[0.1, 0.8]$), $9.7$ points below its score with reaction time ($[5.8, 13.7]$, 14 of 17, $P = 0.013$). A whole-epoch window is necessary for the effect and not sufficient for it.

#### 1.5 Interpretation.

EEG up to and including the response predicts held-out reaction time, and no window ending before the typical response does. The test cannot separate evidence accumulation from the response itself, because every window that carries the effect contains both, and it cannot locate the information within the late window. What it does rule out is that the encoder finds usable held-out information in the first $300$ ms of recorded EEG.

### 2 Variance budget, baselines, and analysis provenance

#### 2.1 The main model's variance budget

The budget of The latent variable sets which trials are slow; accumulator noise sets the spread is a forward partition of the main fit's own reaction-time variance, not a refit and not a prediction score. For trial $n$ with drift trajectory $v_n(t)$, let $\mu_n = \mathbb{E}[\mathrm{RT} \mid v_n]$ and $s_n^2 = \mathrm{Var}[\mathrm{RT} \mid v_n]$ be the conditional moments of the first-passage density. By the law of total variance,

$$\mathrm{Var}(\mathrm{RT}) = \underbrace{\overline{s_n^2}}_{\text{diffusion}} + \underbrace{\textstyle\sum_c \frac{N_c}{N}\left(\bar{\mu}_c - \bar{\mu}\right)^2}_{\text{stimulus}} + \underbrace{\mathrm{Var}_n(\mu_n) - \textstyle\sum_c \frac{N_c}{N}\left(\bar{\mu}_c - \bar{\mu}\right)^2}_{\text{latent variable}}, \tag{S1}$$

with overlines means over trials, $\bar{\mu}_c$ the mean of $\mu_n$ over condition $c$'s $N_c$ trials, and $t_0$, a constant, dropping out. The three terms sum to the model's variance exactly. Averaged over participants, diffusion carries $89.51\%$, the latent variable $10.22\%$, and the stimulus $0.27\%$ (Supplementary Fig. 3 and Supplementary Table 4). The diffusion term is drawn afresh on every trial, and nothing in that trial's EEG or its condition predicts the draw: even with a trial's drift trajectory fixed exactly, the accumulator wanders on its way to the bound, so its crossing time stays random. The latent-variable share exceeded the stimulus share in every participant (17/17, median ratio $122$), ranging from $2.40\%$ in sub-10 to $26.40\%$ in sub-09.

#### 2.2 Baseline and null models

The constraint tests of the main text (Each person's model reproduces their ERP and reaction times and EEG-inferred latent variables predict held-out reaction times when behavior is in the fit) compare the joint model against three reduced or shuffled models, all sharing its frozen anatomy and forward pass.

##### 2.2.1 EEG-only baseline.

The EEG-only baseline asks whether fitting EEG and behavior jointly, rather than EEG alone followed by decoding, is what loads trial-level behavioral information into the shared latent. It is fit exactly like the joint model through warm-start and Stage A, but with the behavior weight set to zero ($w_{\mathrm{beh}} = 0$) and the decision parameters held frozen, so only the encoder $\phi$, the injection scale $\lambda$, the EEG gain $g$ and $\log \sigma_{\mathrm{obs}}$ are trained, to single-trial EEG. The within-condition $z \to$RT readout is then computed *post hoc* by block-cross-validated linear decoding on the frozen encoder, never having entered the fit. The joint model differs only by adding Stages B and C, where the behavior likelihood shapes the same latent.

##### 2.2.2 Behavior-only baseline.

The behavior-only baseline is a per-condition constant-drift diffusion model with no per-trial covariate: a drift $v_c$ for each condition with a shared boundary $a$, relative start $w$ and non-decision time $t_0$, the diffusion scale fixed at $\sigma_a = 1$ (the diffusion-model gauge[1]), fit by maximizing the same Fokker–Planck first-passage likelihood. Because it has no trial-level latent, its predicted reaction time is constant within a condition, so its within-condition $z \to$RT $R^2$ is identically zero. It is the baseline against which the joint and EEG-only within-condition recoveries are judged.

##### 2.2.3 Null-DTI degree-preserving shuffle.

The Null-DTI null replaces a participant's connectome with a degree-preserving shuffle while keeping all other anatomy ($L$, leadfield, region sites) fixed. The shuffle is a Maslov–Sneppen double-edge swap[2] on the symmetric weighted SC: random edge pairs $(a, b), (c, d)$ are rewired to $(a, d), (c, b)$ whenever that creates no self-loop or duplicate, carrying their weights, repeated $10\times$ the edge count; this preserves each node's binary degree exactly but destroys the specific wiring. The topology levers and the

EEG observation parameters are copied from an own-connectome refit and frozen, so the model cannot re-weight its connections around the wrong wiring; only the encoder, the injection scale $\lambda$, and the decision parameters are refit, on five shuffles per participant. This isolates whether the individual topology carries EEG-relevant structure. The region-silencing sensitivity maps (Where speed is set: shared sensorimotor and person-specific parietal effects) reuse the saved intact and shuffled fits.

#### 2.3 Analysis provenance

Every criterion in this paper was fixed before the analysis it governs was run. Two analyses carry provenance details that a reader needs in order to cite their numbers correctly, and they are collected here so that each appears once and in full; the main text points to this note at the claim it qualifies rather than restating it. Supplementary Table 14 first says which fitted model produced which reported number, because the paper uses several and a number read against the wrong one is misread.

##### 2.3.1 The strict held-out test.

The strict test scores the five-fold refits, fitted without each fold's trials, with three statistics taken from the training folds: the condition means against which the latent variable is residualized, the condition-mean evidence that sets the drift's baseline, and the condition means on which predicted and observed reaction times are centered. It pools the five folds so that each trial is scored once, and fixes the slope at $1$ against the training-mean baseline. The scoring rule was written, and its regression tests passed, before the cohort was scored, and the main text reports the outcome as it came out: $9.6\%$ ($95\%$ CI $[6.9, 12.2]$), above zero in 16 of 17 participants, with predicted and observed residuals correlated in all 17 (mean $r = 0.314$, $[0.269, 0.358]$).

##### 2.3.2 Reseeding the identifiability test.

The identifiability statistic fixed in advance is defined on a single base cell per participant. The whole measurement was re-drawn under ten fresh master seeds per participant, resetting both the synthetic keys and the curriculum's random state. Those draws are a readout of the sampling distribution and do not replace that statistic; we report both. The base cell cleared the $0.20$ threshold 9 times of 10 for sub-14 and 9 of 10 for sub-09 (Supplementary Table 12).

### 3 Notation, model overview, and data flow

Supplementary Table 9 collects the indices, tensors, and frozen anatomical operators used throughout the model. Region indices run over the $R = 83$ Lausanne-2018 scale-1 parcels[3,4] and channel indices over the $C = 128$ electrodes of the VEPCON recording[5]; one epoch covers $[-200, +800]$ ms, integrated at $\mathrm{dt}_{\mathrm{dyn}} = 5$ ms ($T_{\mathrm{dyn}} = 200$ dynamics steps) and observed at 100 Hz ($N_{\mathrm{obs}} = 100$ samples), so the observation stride is $T_{\mathrm{dyn}}/N_{\mathrm{obs}} = 2$ (the full time grid and all fixed constants are in Supplementary Table 11).

#### 3.1 Model overview and data flow

Each participant has one joint generative model $p_\theta$ paired with an amortized recognition network $q_\phi$: a variational autoencoder[6] whose decoder is a connectome-constrained neural-mass model. The trial-specific latent variable $z_n$ is the single shared cause of that trial's EEG and behavior. It is inferred from the EEG alone and then forced to explain both the EEG and the reaction time through one differentiable forward pass (Supplementary Fig. 6):

$$\mathrm{EEG}_n \xrightarrow{q_\phi} z_n \longrightarrow \mathrm{drive}_n(t) \xrightarrow{\text{WC+FIC+delays}} \{E_i(t), I_i(t)\} \implies \begin{cases} \mathrm{psp}(t) \xrightarrow{g\mathbf{L}} \widehat{\mathrm{EEG}}_n & \text{(EEG readout)} \\ e_n(t) \xrightarrow{v_n(t)} \text{first passage} & \text{(behavior readout)} \end{cases}$$

Given $z_n$ the two readouts are conditionally independent, so every single-trial correlation between EEG and behavior is carried by $z_n$.

Three properties make that latent interpretable. First, every anatomical forward operator (leadfield $\mathbf{L}$, connectome $C$, fiber lengths $L$ and the derived delays $D$) and the feedback-inhibition working point $w_{IE,i}$ are frozen, leaving only a few regularized degrees of freedom (Supplementary Note 4). Second, the latent enters as the within-condition residual $\tilde{z}_n$ and only into the decision pool, so it adds no condition-specific input; the sensory channels ($V1$, MT, $q$) are $z$-invariant. Through the nonlinear network it can still shift the condition-mean response, which the re-anchor step refits (Methods Fitting protocol). Third, the encoder's input is EEG alone, but its parameters are trained under the joint objective, so the single-trial coupling between $z_n$ and reaction time inside the main fit is a property of the fit; prediction is tested with encoders refit within each cross-validation fold on the training trials, behavior included, and scored on the held-out trials. These are the properties the Results exploit and stress-test: the frozen scaffold and the residualized single-trial link (EEG-inferred latent variables predict held-out reaction times when behavior is in the fit), the reaction-time variance that link accounts for (The latent variable sets which trials are slow; accumulator noise sets the spread), and the labeled decision pool that the sensitivity maps perturb (Where speed is set: shared sensorimotor and person-specific parietal effects).

### 4 Forward-model numerics and parameters

This note expands three numerical components that the main Methods state in one line each, then lists every parameter and constant of the model.

#### 4.1 Feedback-inhibition control (FIC) calibration.

The per-region inhibitory weight $w_{IE,i}$ (Methods Wilson–Cowan node dynamics and feedback-inhibition control) is fixed before training by feedback-inhibition control[7] and never enters the gradient. Calibration is a deterministic, delay-free fixed-point iteration on the static coupling $W = G_0 C$: from an initial $w_{IE,i}$ the network is relaxed and, at each of 600 steps, every weight is nudged

$$w_{IE,i} \leftarrow \mathrm{clip}(w_{IE,i} + \eta\,(\mathrm{in}_{E,i} - \theta_E),\ [0.05, 10])\,, \qquad \eta = 0.05, \tag{S2}$$

raising inhibition where a region's excitatory input current overshoots $\theta_E$ and lowering it where it falls short, until $\mathrm{in}_{E,i} \to \theta_E$ (so $E_i \to 0.5$). Calibrating on the delay-free system is valid for the delayed network: at a fixed point each region's delayed presynaptic activity equals its instantaneous value, $E_j(t - D_{ij}) = E_j^*$, so the delayed coupling term collapses to the instantaneous one and the calibrated $w_{IE,i}$ continues to hold once conduction delays are restored. During training the working point is maintained softly by the penalty $\mathcal{R}_{\mathrm{wp}}$ (Methods Objective: negative ELBO).

#### 4.2 Fokker–Planck first-passage solver.

Because the drift of Eq. 22 varies in time, the first-passage density has no closed form and is obtained by forward integration of the Fokker–Planck equation on a grid[8] (Methods Decision readout and first-passage likelihood). The interior $(0, a)$ is discretized into $n_{\mathrm{grid}} = 128$ bins and the density is initialized as a point mass at $x(0) = w\,a$. Each dynamics step convolves the density with a Gaussian transition kernel of mean $x + v_n(t)\,\mathrm{d}t$ and variance $\sigma_a^2\,\mathrm{d}t$; probability mass crossing $0$ or $a$ is removed by the mirror (method-of-images) construction and accumulated as boundary flux, and the flux per unit time at each boundary is the first-passage density for the corresponding choice. The log-likelihood interpolates this density at $\mathrm{rt}_n - t_0$, and the scan over time steps is gradient-checkpointed to bound memory. A small fraction of mass leaks past the grid edges within the integration window (cohort mean $1.59\%$, median $0.81\%$, range $0.04$–$4.49\%$, $N = 17$; Supplementary Table 4); the leak falls almost entirely on the within-trial diffusion term and shifts absolute variances slightly without changing the variance *shares* of Supplementary Note 2.

#### 4.3 Differentiable conduction-delay buffer.

Inter-regional coupling reads each source at its own fiber-length lag (Methods Effective connectivity, neuromodulation and delayed coupling), $u_{n,i}(t) = \sum_j W^{\text{eff}}_{n,ij}(t)\, E_{n,j}(t - D_{ij})$. The delayed activity is supplied by a shift buffer of regional excitatory states whose pre-stimulus lags are initialized at the calibrated working point $E^* \approx 0.5$ rather than at zero, so the buffer fills without injecting a spurious onset transient. The buffer is advanced each step and the delayed current is formed by a differentiable gather over per-edge lags, so gradients flow back through the delayed coupling.

#### 4.4 Parameters: fixed, calibrated, and learned

The model keeps the number of free parameters small and bounded. Anatomy enters frozen; the feedback-inhibition working point is calibrated offline and frozen; a handful of effective-connectivity and stimulus levers are fit during warm-start, frozen for the curriculum, and refit in the re-anchor step; and only the encoder, the injection scale, the observation gain, and the six decision parameters move during the joint curriculum, with the decision parameters refit once more in the recalibration step. Supplementary Table 11 is the full ledger of every quantity: the step at which it is trained, its constraint transform, and its initial or fixed value, with the fixed constants and fitting hyperparameters in its last two blocks.

### 5 Identifiability and synthetic-data recovery

A good fit does not mean that the model's parts are pinned down. This note gives the synthetic-data recovery test in full: what could fail, what came back, how much of the outcome is a single draw, and what the test's criterion cannot see. The design is in Methods Identifiability and synthetic-data recovery.

#### 5.1 Degeneracies tested.

The latent variable reaches behavior through two scales in series: the injection scale $\lambda$, which adds $z_n$ to the decision pool, and the drift scale $\beta$, which turns pooled evidence into drift. Only their product $\beta\lambda$ scales a trial's evidence, so the data can fix the product while leaving each factor free to slide. The other failure is simpler: the latent variable's imprint on the EEG could be too faint for any encoder to read $z_n$ back at all. Either one would strip the basis from every absolute magnitude in the Results.

#### 5.2 Test and outcome.

We tested both on data whose true answer we already knew. Trials were generated from the frozen model at set values of $\beta$ and $\lambda$, then refit from scratch by the same training curriculum, and we asked how much of the truth came back. Two participants, sub-14 and sub-09, were chosen before any recovery ran by the size of the forward EEG imprint of their latent variable, measured as $\|\partial\, \mathrm{ERP}_n / \partial \tilde{z}_n\| / \sigma_{\text{obs}}$, and each was swept over an $11 \times 11$ grid of $\beta \times \lambda$ working points, 121 refits per participant, with the test statistic read at the base cell of that grid. The recovered latent variable tracked the true one at $|r| = 0.340$ (sub-09) and $0.233$ (sub-14), against a $0.20$ threshold on a statistic whose information ceiling is $\approx 0.46$, and $\beta$ and $\lambda$ came back separately in both participants (Supplementary Table 12). Two further participants, sub-10 and sub-18, whose latent variables leave a smaller EEG imprint, were swept on the same grid as reference points on the recovery floor; they are not part of the test.

#### 5.3 Reseeding.

Each of those numbers is a single draw, so the outcome could be luck. We re-ran the whole measurement under ten fresh master seeds per participant, each resetting the synthetic data and the curriculum's random state. The threshold was cleared 9 times in 10 for sub-14 and 9 in 10 for sub-09 (Supplementary Fig. 7a and Supplementary Table 12), so the original outcome was not a matter of which seed we drew. The statistic fixed in advance remains the single base cell, and these draws sit beside it rather than replace it (Supplementary Note 2).

#### 5.4 Partial correlation and recovery slope.

Clearing the criterion is not the same as pinning $\lambda$ down, and the criterion cannot see the difference. It is a partial correlation, which asks only whether the estimate moves in the right direction when the truth changes; it is silent on how far the estimate moves. The log–log recovery slope asks that second question, and across the same $11 \times 11$ sweep the two answers diverge. The $\lambda$ partial correlations are $+0.400$ for sub-14 and $+0.533$ for sub-09, so both clear the criterion; but over a grid covering a $4.00\times$ range in $\lambda^*$, their slopes reproduce only $5\%$ and $31\%$ of that variation. In sub-09 the compensation is explicit: $\hat{\lambda}$ responds more strongly to $\beta^*$ ($-0.410$) than to its own $\lambda^*$ ($+0.310$), so an error in one scale is absorbed by the other along a line of constant $\beta\lambda$, which is the trade-off the test was built to detect. We therefore claim that $\beta$ and $\lambda$ each carry identifiable information, not that either is pinned. The two criteria are drawn side by side in Supplementary Fig. 7b. With 121 cells a bootstrap interval excludes zero above $r = 0.18$, so the interval and the $0.20$ threshold fixed in advance both bind here.

One property of the sweep limits how the test statistic may be summarized. Latent recovery improves systematically with $\lambda^*$ across the grid ($\mathrm{d}\log|r|/\mathrm{d}\log\lambda^* = +0.93$ and $+0.39$ in the two participants), so the statistic is read at the base cell rather than as a median over the whole grid: that median would be taken mostly over cells with $\lambda^* \neq 1$, which are the easier ones.

The two scales are the hard case, and the rest of the decision parameters recover unevenly. Pooled over the two participants and 121 grid cells, the recovered-to-true ratios for $\beta$, the starting point $w$, the non-decision time $t_0$, and the diffusion noise $\sigma$ have intervals covering $1$, with medians $0.91$, $0.76$, $0.92$, and $0.71$. The diffusion noise can be recovered here because the configuration this test used leaves $\sigma_a$ free, whereas the main fit fixes it at $1$ (Methods Identifiability and synthetic-data recovery). The bound $a$ is under-recovered across its whole interval (median $0.75$, $2.5$–$97.5\%$ band $[0.17, 0.83]$), and the drift bias $b$, which is non-positive and near-degenerate, is only weakly identifiable (Supplementary Fig. 7c, $n = 242$ refits). Bound and non-decision time are compared within participants rather than as cross-participant point estimates in any case, for a reason given before these numbers were measured (Methods Identifiability and synthetic-data recovery).

### 6 Supplementary Tables

Supplementary Table 1: **Connectome-specificity and joint-fit tests.** Paired cohort contrasts, $N = 17$. $\overline{\Delta}$ is the own-minus-comparison mean, $N^+$ counts participants moving in the predicted direction, $P$ is the two-sided sign test, and MDE is the minimum detectable effect at that $N$. The upper block replaces the individual connectome with a degree-preserving (Null-DTI) shuffle and reads two channels: ERP reconstruction and the within-condition single-trial recovery $z_{\mathrm{within}}\ R^2$. With the topology levers and observation parameters frozen and only the encoder, $\lambda$, and the decision parameters refit, ERP reconstruction collapses in every participant. The behavioral channel cannot test the connectome, because the latent variable enters the decision pool's evidence directly (Eq. 16) and the connectome reaches it only through the network state; it moves against the predicted direction in 15 of 17. We therefore confine the connectome-specificity claim to ERP generation. The lower block is the joint-fit test: the joint model exceeds an EEG-only post-hoc readout, while that readout is statistically equivalent to a behavior-only DDM baseline within a $\pm 0.02$ bound fixed in advance, so the advantage comes from generating both channels together rather than from the encoder alone. The Wilcoxon signed-rank $P$ for the joint contrast is $1.5 \times 10^{-5}$. In the upper block the own fit is the main fit and the shuffle arm a refit on a shorter schedule, so each $\overline{\Delta}$ there holds a training-budget gap as well as the connectome effect. The refit-against-refit connectome pairing leaves $+0.832$ ($[+0.709, +0.959]$, 17/17) for ERP $r$ and $-0.019$ ($[-0.038, -0.002]$, 6/17, $P = 0.332$) for $z_{\mathrm{within}}$. Fig. 3b draws the joint, EEG-only post-hoc, and DDM columns.

| Contrast | Channel | $\overline{\Delta}$ | 95% CI | $N^+$ | $P$ | MDE |
|---|---|---|---|---|---|---|
| *Own connectome vs Null-DTI shuffle* | | | | | | |
| own − shuffle | ERP $r$ | $+0.881$ | $[+0.764, +1.001]$ | 17/17 | $1.5 \times 10^{-5}$ | 0.175 |
| own − shuffle | $z_{\mathrm{within}}\ R^2$ | $-0.042$ | $[-0.065, -0.021]$ | 2/17 | 0.0023 | 0.032 |
| *Joint generation vs post-hoc pipelines* | | | | | | |
| joint − EEG-only post-hoc | $z_{\mathrm{within}}\ R^2$ | $+0.519$ | $[+0.479, +0.557]$ | 17/17 | $1.5 \times 10^{-5}$ | 0.058 |
| EEG-only post-hoc − DDM baseline | $z_{\mathrm{within}}\ R^2$ | $-0.002$ | $[-0.005, +0.001]$ | – | $5.5 \times 10^{-10}$ [a] | 0.004 |

[a] Two one-sided tests against a $\pm 0.02$ equivalence bound; small $P$ means equivalent, not different.

Supplementary Table 2: **Per-participant face validity.** The values behind Fig. 2e–h. For each participant: the number of usable trials; the element-wise correlation $r$ between predicted and observed condition-mean ERP; that participant's split-half noise ceiling, the square root of the Spearman–Brown reliability (Methods Face-validity checks), and the ratio of the two (Fig. 2e); the reaction-time slow-tail bias, predicted minus observed $0.9$ quantile (Fig. 2g); whether an emergent global-field-power peak fell in each of the P1, N1, and P3 windows within $\pm 30$ ms, with the latencies themselves plotted in Fig. 2f; and the single-trial EEG reconstruction $r^2$ against the condition-mean baseline. Cohort medians over the 17 analyzed participants: $r = 0.858$, ceiling $0.969$, ratio $0.881$, tail $+0.1$ ms, single-trial $r^2 = -0.047$; emergent-peak counts P1 $5$ / N1 $9$ / P3 $14$, median one window per participant. All 18 fitted participants clear the $r > 0.60$ criterion. †sub-16 clears the criterion but has too few usable trials, and is excluded from every cohort statistic and figure, which run at $N = 17$. The 14 participants with a P3 hit are exactly the subset analyzed in How speed is set: network and readout routes. Expands Supplementary Fig. 1.

| Participant | trials | $r$ | ceiling | ratio | tail (ms) | P1 | N1 | P3 | single-trial $r^2$ |
|---|---|---|---|---|---|---|---|---|---|
| sub-01 | 569 | 0.632 | 0.973 | 0.649 | $+48.4$ | – | – | ✓ | $-0.070$ |
| sub-02 | 497 | 0.755 | 0.969 | 0.779 | $+0.1$ | ✓ | ✓ | ✓ | $-0.052$ |
| sub-03 | 571 | 0.925 | 0.989 | 0.936 | $-28.0$ | – | ✓ | – | $-0.085$ |
| sub-04 | 554 | 0.878 | 0.989 | 0.887 | $-20.0$ | ✓ | ✓ | – | $-0.116$ |
| sub-06 | 571 | 0.677 | 0.912 | 0.743 | $-42.2$ | – | – | ✓ | $-0.016$ |
| sub-07 | 520 | 0.800 | 0.935 | 0.856 | $-17.8$ | – | – | – | $-0.016$ |
| sub-08 | 561 | 0.877 | 0.984 | 0.891 | $+4.0$ | ✓ | ✓ | ✓ | $-0.083$ |
| sub-09 | 498 | 0.854 | 0.961 | 0.889 | $+17.4$ | ✓ | ✓ | ✓ | $-0.037$ |
| sub-10 | 520 | 0.858 | 0.974 | 0.881 | $+43.4$ | – | – | ✓ | $-0.071$ |
| sub-11 | 559 | 0.791 | 0.964 | 0.821 | $+7.8$ | – | ✓ | ✓ | $-0.031$ |
| sub-12 | 471 | 0.783 | 0.967 | 0.810 | $-35.1$ | – | ✓ | ✓ | $-0.047$ |
| sub-13 | 581 | 0.941 | 0.989 | 0.951 | $-19.7$ | – | – | ✓ | $-0.057$ |
| sub-14 | 550 | 0.880 | 0.961 | 0.916 | $-6.0$ | – | ✓ | ✓ | $-0.029$ |
| sub-16† | 216 | 0.830 | 0.954 | 0.870 | $-18.4$ | ✓ | – | – | $-0.042$ |
| sub-17 | 577 | 0.877 | 0.971 | 0.904 | $+10.4$ | – | – | ✓ | $-0.027$ |
| sub-18 | 557 | 0.859 | 0.952 | 0.903 | $+48.3$ | – | – | ✓ | $-0.010$ |
| sub-19 | 524 | 0.805 | 0.929 | 0.866 | $+10.3$ | ✓ | ✓ | ✓ | $-0.010$ |
| sub-20 | 565 | 0.860 | 0.984 | 0.875 | $-15.0$ | – | – | ✓ | $-0.066$ |

Supplementary Table 3: **Held-out reaction-time $R^2$ by EEG window, for the joint encoder.** Cohort means over $N = 17$ participants, in percent of within-condition reaction-time variance, with participant-level bootstrap $95\%$ CIs. The encoder is refit within each fold, and the rows are the windows it was allowed to see. $N^+$ counts participants above zero in the left block and participants moving in the predicted direction in the right block, where the quantity is the whole epoch minus that row's window, so a positive value means the whole epoch reads better. $P$ is the two-sided sign test on that paired difference. Every window starts at the start of the epoch. Fig. 3a draws every row.

| Window | $R^2$ [95% CI] | $N^+$ | Whole − window [95% CI] | $N^+$ | $P$ |
|---|---|---|---|---|---|
| $\leq 200$ ms | $-12.6$ $[-15.4, -9.9]$ | 0/17 | $+22.8$ $[+18.4, +27.5]$ | 17/17 | $1.5 \times 10^{-5}$ |
| $\leq 300$ ms | $-8.0$ $[-12.1, -4.1]$ | 4/17 | $+18.2$ $[+13.3, +23.5]$ | 17/17 | $1.5 \times 10^{-5}$ |
| $\leq 600$ ms | $0.0$ $[-3.8, +3.7]$ | 8/17 | $+10.3$ $[+7.2, +13.2]$ | 15/17 | $0.0023$ |
| whole epoch | $+10.2$ $[+6.2, +14.4]$ | 14/17 | – | – | – |
| whole epoch [a] | $+0.5$ $[+0.1, +0.8]$ | 12/17 | $+9.7$ $[+5.8, +13.7]$ | 14/17 | $0.013$ |

[a] The same encoder refit on the whole epoch without reaction time in its objective.

Supplementary Table 4: **Within-participant reaction-time variance partition, and the fixed and permuted latent-variable ablations.** Left block: each participant's single-trial reaction-time variance split into a stimulus component, a recoverable latent-variable component carried by the latent variable $z_n$, and irreducible diffusion noise, in percent. Every participant is quantifiable on the four premises fixed in advance, so the partition is defined at $N = 17$. Diffusion dominates throughout, but the latent-variable share is larger than the stimulus share in every participant (cohort means $0.27\%$ stimulus, $10.22\%$ latent variable, $89.51\%$ diffusion), with a range of $2.40\%$ (sub-10) to $26.40\%$ (sub-09). *leak* is the Fokker–Planck mass not absorbed inside the integration window, all within the pre-set tolerance. *model/obs* is the ratio of predicted to observed reaction-time variance; values above one show the model over-dispersing, most strongly at sub-13 ($1.77$). This partition is descriptive and in-sample, and it inherits the marginalized likelihood: it partitions reaction-time variance only. Right block: the two ablations, each one forward Fokker–Planck pass on the already-fitted model with no re-training. Fixing replaces each trial's evidence with its within-condition mean; permuting keeps every trial's evidence and shuffles which trial it belongs to, using 200 within-condition draws at seed $0$. $r_{\text{full}}$ and $r_{\text{permuted}}$ are the correlations between predicted and observed within-condition reaction-time residuals for the intact and the permuted model, and *null* is the $5$–$95\%$ band of the permutation draws. The fixed arm has no correlation to report. Its prediction is constant inside a condition, so the within-condition residual it would be scored on is zero to double precision (largest s.d. across the 17 participants, $1.1 \times 10^{-16}$ s), and $r$ has no value rather than a small one, so the two pairing panels Fig. 4d and e do not draw the arm at all and no column is given for it. Its distance from the intact reaction-time density is a real measurement, the total-variation distance $\mathrm{TV}(\text{full}, \text{fixed-}z)$ in the last column, median $0.0432$ and drawn for one participant in Fig. 4c. The intact model clears each participant's own band in 17/17 and the permuted arm falls outside in 3/17, close to the $\approx 1.7$ a $5$–$95\%$ band excludes by construction, so that count is not a score. $\mathrm{TV}(\text{full}, \text{permuted})$ is not tabulated: a within-condition row permutation commutes with the per-trial average, so the quantity is identically zero and what the artifact stores is a floating-point residual. The ablations are descriptive and in-sample, with no criterion fixed in advance.

| | | variance partition (%) | | | | | latent-variable ablations | | | |
|---|---|---|---|---|---|---|---|---|---|---|
| Participant | trials | stimulus | latent | diffusion | leak | model/obs | $r_{\text{full}}$ | $r_{\text{permuted}}$ | null $5$–$95\%$ | TV(fixed-$z$) |
| sub-01 | 569 | 0.06 | 9.36 | 90.57 | 0.50 | 1.26 | $+0.773$ | $+0.032$ | $[-0.084, +0.067]$ | 0.0446 |
| sub-02 | 497 | 0.02 | 13.36 | 86.63 | 0.12 | 1.16 | $+0.684$ | $+0.069$ | $[-0.074, +0.065]$ | 0.0600 |
| sub-03 | 571 | $< 0.01$ | 7.26 | 92.73 | 2.62 | 1.55 | $+0.670$ | $+0.091$ | $[-0.070, +0.073]$ | 0.0407 |
| sub-04 | 554 | $< 0.01$ | 3.76 | 96.24 | 0.72 | 1.49 | $+0.586$ | $+0.066$ | $[-0.071, +0.067]$ | 0.0165 |
| sub-06 | 571 | 1.73 | 7.58 | 90.69 | 4.49 | 1.14 | $+0.738$ | $+0.017$ | $[-0.067, +0.073]$ | 0.0350 |
| sub-07 | 520 | 0.05 | 8.55 | 91.40 | 0.26 | 1.26 | $+0.642$ | $-0.028$ | $[-0.079, +0.070]$ | 0.0316 |
| sub-08 | 561 | 0.71 | 2.83 | 96.47 | 3.14 | 1.11 | $+0.614$ | $-0.015$ | $[-0.074, +0.072]$ | 0.0198 |
| sub-09 | 498 | 0.22 | 26.40 | 73.38 | 0.30 | 1.21 | $+0.813$ | $-0.018$ | $[-0.068, +0.077]$ | 0.1075 |
| sub-10 | 520 | 0.03 | 2.40 | 97.57 | 2.74 | 1.13 | $+0.651$ | $-0.002$ | $[-0.073, +0.084]$ | 0.0156 |
| sub-11 | 559 | 0.81 | 13.20 | 85.99 | 1.01 | 1.12 | $+0.753$ | $-0.034$ | $[-0.076, +0.069]$ | 0.0616 |
| sub-12 | 471 | 0.06 | 6.07 | 93.87 | 4.22 | 1.29 | $+0.708$ | $+0.047$ | $[-0.073, +0.068]$ | 0.0269 |
| sub-13 | 581 | 0.06 | 11.39 | 88.54 | 0.04 | 1.77 | $+0.677$ | $-0.001$ | $[-0.071, +0.064]$ | 0.0432 |
| sub-14 | 550 | $< 0.01$ | 17.01 | 82.99 | 4.47 | 1.03 | $+0.692$ | $+0.022$ | $[-0.073, +0.069]$ | 0.0763 |
| sub-17 | 577 | 0.44 | 14.65 | 84.91 | 0.04 | 1.15 | $+0.664$ | $+0.013$ | $[-0.065, +0.075]$ | 0.0578 |
| sub-18 | 557 | 0.15 | 10.26 | 89.59 | 0.28 | 1.27 | $+0.652$ | $+0.022$ | $[-0.061, +0.067]$ | 0.0599 |
| sub-19 | 524 | 0.18 | 9.17 | 90.64 | 0.81 | 1.27 | $+0.708$ | $-0.001$ | $[-0.073, +0.070]$ | 0.0393 |
| sub-20 | 565 | 0.02 | 10.57 | 89.41 | 1.32 | 1.09 | $+0.506$ | $-0.098$ | $[-0.064, +0.061]$ | 0.0442 |
| mean | | 0.27 | 10.22 | 89.51 | 1.59 | 1.25 | | | | |
| median | | | | | | | $+0.677$ | $+0.013$ | | 0.0432 |

Supplementary Table 5: **Lausanne-2018 scale-1 parcel index.** The 83 regions of the Wilson–Cowan network (Methods Dataset and task), with the indices that name them in Supplementary Tables 7 and 13. The parcellation is a fixed atlas, so an index denotes the same region in every participant. Indices are 0-based and run right hemisphere, then left, then brainstem, so the left-hemisphere index of a region is its right-hemisphere index plus $41$. Region names translate the FreeSurfer label. The grouping is by lobe rather than by functional network because scale-1 parcels are large enough to span more than one network. ‡Five parcels that the main text and Methods call by a functional shorthand: V1 (pericalcarine and cuneus, the visual injection sites of Eq. 16), MT/V5 (middle temporal), pre-SMA/SMA (superior frontal), and M1 (precentral). Each shorthand names an area that occupies part of the parcel carrying it, not the whole parcel, so the parcels are proxies, as stated for the decision pool in Methods Decision readout and first-passage likelihood.

| Parcellation label | Region | R | L |
|---|---|---|---|
| *Frontal* | | | |
| lateralorbitofrontal | lateral orbitofrontal cortex | 0 | 41 |
| parsorbitalis | inferior frontal gyrus, pars orbitalis | 1 | 42 |
| frontalpole | frontal pole | 2 | 43 |
| medialorbitofrontal | medial orbitofrontal cortex | 3 | 44 |
| parstriangularis | inferior frontal gyrus, pars triangularis | 4 | 45 |
| parsopercularis | inferior frontal gyrus, pars opercularis | 5 | 46 |
| rostralmiddlefrontal | rostral middle frontal gyrus | 6 | 47 |
| superiorfrontal | superior frontal gyrus‡ | 7 | 48 |
| caudalmiddlefrontal | caudal middle frontal gyrus | 8 | 49 |
| precentral | precentral gyrus (primary motor)‡ | 9 | 50 |
| paracentral | paracentral lobule | 10 | 51 |
| *Cingulate* | | | |
| rostralanteriorcingulate | rostral anterior cingulate cortex | 11 | 52 |
| caudalanteriorcingulate | caudal anterior cingulate cortex | 12 | 53 |
| posteriorcingulate | posterior cingulate cortex | 13 | 54 |
| isthmuscingulate | isthmus of the cingulate (retrosplenial) | 14 | 55 |
| *Parietal* | | | |
| postcentral | postcentral gyrus (primary somatosensory) | 15 | 56 |
| supramarginal | supramarginal gyrus | 16 | 57 |
| superiorparietal | superior parietal cortex | 17 | 58 |
| inferiorparietal | inferior parietal cortex | 18 | 59 |
| precuneus | precuneus | 19 | 60 |
| *Occipital* | | | |
| cuneus | cuneus‡ | 20 | 61 |
| pericalcarine | pericalcarine cortex (primary visual)‡ | 21 | 62 |
| lateraloccipital | lateral occipital cortex | 22 | 63 |
| lingual | lingual gyrus | 23 | 64 |
| *Temporal and insular* | | | |
| fusiform | fusiform gyrus | 24 | 65 |
| parahippocampal | parahippocampal gyrus | 25 | 66 |
| entorhinal | entorhinal cortex | 26 | 67 |
| temporalpole | temporal pole | 27 | 68 |
| inferiortemporal | inferior temporal gyrus | 28 | 69 |
| middletemporal | middle temporal gyrus‡ | 29 | 70 |
| bankssts | banks of the superior temporal sulcus | 30 | 71 |
| superiortemporal | superior temporal gyrus | 31 | 72 |
| transversetemporal | transverse temporal gyrus (primary auditory) | 32 | 73 |
| insula | insula | 33 | 74 |
| *Subcortical* | | | |
| thalamus_proper | thalamus | 34 | 75 |

*continued on the next page*

Supplementary Table 5, continued.

| Parcellation label | Region | R | L |
|---|---|---|---|
| caudate | caudate nucleus | 35 | 76 |
| putamen | putamen | 36 | 77 |
| pallidum | globus pallidus | 37 | 78 |
| accumbens_area | nucleus accumbens | 38 | 79 |
| hippocampus | hippocampus | 39 | 80 |
| amygdala | amygdala | 40 | 81 |
| *Brainstem* | | | |
| brainstem | brainstem | 82 | |

Supplementary Table 6: **Region-by-region cohort table of the intervention maps.** All 83 Lausanne-2018 scale-1 parcels in the anatomical order of Fig. 5b, with the region's role in the model (decision pool, injection site, or none), its membership in the literature set, the number of participants its cut robustly slowed and sped, the number placing it among their eight strongest regions, the cohort median $\Delta\mathrm{RT}$ (ms; positive slows), the number of participants in which its scalp change exceeds the participant's own ERP band, and the cohort class (slowing, speeding, mixed, or sparse; Methods Sensitivity maps).

| Index | Region | Role | Literature set | Participants slowed | sped | top-8 | median $\Delta\mathrm{RT}$ (ms) | scalp robust | Class |
|---|---|---|---|---|---|---|---|---|---|
| *Frontal* | | | | | | | | | |
| 0 | R lat. orbitofrontal | – | – | 5 | 5 | 1 | $+0.0$ | 17 | mixed |
| 41 | L lat. orbitofrontal | – | – | 8 | 3 | 0 | $+1.0$ | 15 | mixed |
| 1 | R pars orbitalis | – | – | 2 | 3 | 0 | $-0.1$ | 10 | sparse |
| 42 | L pars orbitalis | – | – | 3 | 2 | 0 | $-0.2$ | 14 | sparse |
| 2 | R frontal pole | – | – | 5 | 2 | 0 | $-0.1$ | 13 | sparse |
| 43 | L frontal pole | – | – | 4 | 0 | 0 | $+0.6$ | 8 | sparse |
| 3 | R med. orbitofrontal | – | – | 5 | 3 | 0 | $+0.1$ | 16 | mixed |
| 44 | L med. orbitofrontal | – | – | 1 | 5 | 1 | $-0.5$ | 15 | sparse |
| 4 | R pars triangularis | – | – | 4 | 3 | 0 | $-0.4$ | 15 | mixed |
| 45 | L pars triangularis | – | – | 3 | 2 | 0 | $-0.2$ | 14 | sparse |
| 5 | R pars opercularis | – | – | 1 | 4 | 0 | $-0.6$ | 14 | sparse |
| 46 | L pars opercularis | – | – | 4 | 3 | 0 | $+0.1$ | 14 | mixed |
| 6 | R rostral mid. frontal | literature | control | 5 | 6 | 2 | $-0.9$ | 15 | mixed |
| 47 | L rostral mid. frontal | literature | control | 8 | 5 | 0 | $+0.0$ | 15 | mixed |
| 7 | R sup. frontal | literature | threshold | 12 | 3 | 4 | $+3.8$ | 16 | slowing |
| 48 | L sup. frontal | literature | threshold | 11 | 2 | 2 | $+4.1$ | 17 | slowing |
| 8 | R caudal mid. frontal | literature | control | 3 | 3 | 0 | $-0.3$ | 12 | mixed |
| 49 | L caudal mid. frontal | literature | control | 0 | 7 | 0 | $-1.3$ | 13 | sparse |
| 9 | R precentral | literature | motor | 9 | 2 | 4 | $+1.3$ | 17 | slowing |
| 50 | L precentral | literature | motor | 12 | 1 | 2 | $+2.9$ | 17 | slowing |
| 10 | R paracentral | – | – | 2 | 8 | 0 | $-0.6$ | 14 | sparse |
| 51 | L paracentral | – | – | 4 | 6 | 0 | $-0.5$ | 15 | mixed |
| *Cingulate* | | | | | | | | | |
| 11 | R rostral ant. cingulate | – | – | 2 | 4 | 0 | $-0.3$ | 17 | sparse |
| 52 | L rostral ant. cingulate | – | – | 6 | 3 | 0 | $+0.1$ | 14 | mixed |
| 12 | R caudal ant. cingulate | – | – | 2 | 8 | 0 | $-1.4$ | 13 | sparse |
| 53 | L caudal ant. cingulate | – | – | 1 | 7 | 1 | $-0.5$ | 15 | sparse |
| 13 | R post. cingulate | – | – | 6 | 5 | 0 | $-0.4$ | 15 | mixed |
| 54 | L post. cingulate | – | – | 5 | 5 | 1 | $+0.7$ | 15 | mixed |
| 14 | R isthmus cingulate | – | – | 6 | 6 | 0 | $+1.1$ | 14 | mixed |
| 55 | L isthmus cingulate | – | – | 3 | 7 | 0 | $-0.7$ | 15 | mixed |
| *Parietal* | | | | | | | | | |
| 15 | R postcentral | – | – | 10 | 3 | 1 | $+1.9$ | 17 | slowing |
| 56 | L postcentral | – | – | 15 | 1 | 4 | $+8.6$ | 16 | slowing |
| 16 | R supramarginal | pool | accumulation | 9 | 5 | 1 | $+1.9$ | 16 | slowing |
| 57 | L supramarginal | pool | accumulation | 9 | 8 | 1 | $+1.6$ | 16 | slowing |
| 17 | R sup. parietal | pool | accumulation | 8 | 6 | 11 | $-0.1$ | 17 | mixed |
| 58 | L sup. parietal | pool | accumulation | 9 | 7 | 8 | $+5.8$ | 17 | slowing |
| 18 | R inf. parietal | pool | accumulation | 7 | 8 | 4 | $+0.6$ | 17 | mixed |
| 59 | L inf. parietal | pool | accumulation | 9 | 7 | 6 | $+1.8$ | 17 | slowing |
| 19 | R precuneus | pool | accumulation | 6 | 8 | 5 | $+0.2$ | 17 | mixed |
| 60 | L precuneus | pool | accumulation | 7 | 8 | 2 | $+0.6$ | 16 | mixed |
| *Occipital* | | | | | | | | | |
| 20 | R cuneus | injection | – | 3 | 7 | 0 | $-0.8$ | 17 | mixed |
| 61 | L cuneus | injection | – | 7 | 7 | 2 | $-0.1$ | 16 | mixed |
| 21 | R pericalcarine | injection | – | 9 | 5 | 5 | $+1.7$ | 17 | slowing |
| 62 | L pericalcarine | injection | – | 11 | 4 | 2 | $+2.1$ | 17 | slowing |
| 22 | R lat. occipital | – | – | 6 | 10 | 7 | $-4.9$ | 17 | speeding |
| 63 | L lat. occipital | – | – | 7 | 6 | 2 | $+0.3$ | 17 | mixed |
| 23 | R lingual | – | – | 12 | 3 | 2 | $+3.0$ | 17 | slowing |
| 64 | L lingual | – | – | 10 | 7 | 2 | $+4.0$ | 17 | slowing |
| *Temporal* | | | | | | | | | |
| 24 | R fusiform | – | – | 7 | 9 | 2 | $-1.2$ | 17 | speeding |
| 65 | L fusiform | – | – | 9 | 6 | 6 | $+2.5$ | 17 | slowing |
| 25 | R parahippocampal | – | – | 7 | 3 | 1 | $+0.9$ | 16 | mixed |

| | | | | | | | | | |
|---|---|---|---|---|---|---|---|---|---|
| 66 | L parahippocampal | – | – | 7 | 5 | 1 | +0.5 | 17 | mixed |
| 26 | R entorhinal | – | – | 9 | 1 | 0 | +1.0 | 14 | slowing |
| 67 | L entorhinal | – | – | 2 | 5 | 1 | −0.2 | 16 | sparse |
| 27 | R temporal pole | – | – | 4 | 3 | 0 | −0.4 | 15 | mixed |
| 68 | L temporal pole | – | – | 4 | 6 | 0 | −0.3 | 14 | mixed |
| 28 | R inf. temporal | – | – | 4 | 9 | 3 | −2.6 | 16 | speeding |
| 69 | L inf. temporal | – | – | 6 | 8 | 3 | +0.2 | 16 | mixed |
| 29 | R mid. temporal | pool | evidence | 1 | 14 | 3 | −5.4 | 17 | speeding |
| 70 | L mid. temporal | pool | evidence | 5 | 11 | 7 | −6.7 | 17 | speeding |
| 30 | R banks STS | – | – | 5 | 9 | 2 | −1.7 | 16 | speeding |
| 71 | L banks STS | – | – | 4 | 9 | 2 | −1.5 | 16 | speeding |
| 31 | R sup. temporal | – | – | 9 | 3 | 1 | +1.4 | 17 | slowing |
| 72 | L sup. temporal | – | – | 5 | 9 | 4 | −1.6 | 17 | speeding |
| 32 | R transverse temporal | – | – | 5 | 4 | 0 | +0.2 | 15 | mixed |
| 73 | L transverse temporal | – | – | 5 | 3 | 0 | −0.1 | 14 | mixed |
| *Insula* | | | | | | | | | |
| 33 | R insula | – | – | 5 | 10 | 2 | −1.6 | 16 | speeding |
| 74 | L insula | – | – | 4 | 10 | 2 | −1.4 | 17 | speeding |
| *Subcortical* | | | | | | | | | |
| 34 | R thalamus | – | – | 9 | 5 | 0 | +3.3 | 15 | slowing |
| 75 | L thalamus | – | – | 11 | 2 | 1 | +3.7 | 16 | slowing |
| 35 | R caudate | – | – | 5 | 6 | 1 | −0.0 | 16 | mixed |
| 76 | L caudate | – | – | 5 | 5 | 0 | −0.0 | 15 | mixed |
| 36 | R putamen | – | – | 12 | 4 | 2 | +5.9 | 17 | slowing |
| 77 | L putamen | – | – | 9 | 3 | 2 | +1.7 | 16 | slowing |
| 37 | R pallidum | – | – | 2 | 5 | 0 | −0.4 | 16 | sparse |
| 78 | L pallidum | – | – | 5 | 4 | 2 | −0.5 | 14 | mixed |
| 38 | R accumbens | – | – | 1 | 3 | 0 | +0.1 | 11 | sparse |
| 79 | L accumbens | – | – | 2 | 1 | 0 | −0.4 | 10 | sparse |
| 39 | R hippocampus | – | – | 5 | 6 | 2 | −0.9 | 15 | mixed |
| 80 | L hippocampus | – | – | 4 | 4 | 0 | +0.1 | 17 | mixed |
| 40 | R amygdala | – | – | 3 | 3 | 1 | −0.6 | 15 | mixed |
| 81 | L amygdala | – | – | 3 | 3 | 0 | −0.2 | 15 | mixed |
| *Brainstem* | | | | | | | | | |
| 82 | brainstem | – | – | 5 | 7 | 2 | −0.1 | 17 | mixed |

Supplementary Table 7: **Per-participant strongest region.** For each participant: the intact predicted reaction time, the noise band (the 95th percentile of the absolute deviation from the median of the readout over 100 re-simulations with fresh noise and no cut), the number of the 83 regions whose cut moves the reaction time beyond that band, and the strongest region with its role (decision pool, injection site, literature set, or other), its $\Delta\mathrm{RT}$, and the region's cohort class (Fig. 5b,c). Positive $\Delta\mathrm{RT}$ means the cut slows the participant. Regions are Lausanne-2018 scale-1 parcels (Supplementary Table 5); all values are under the fitted pool readout (Methods Sensitivity maps). All 17 pass the existence criterion fixed in advance, though the criterion is a weak lower bound: the bands ($0.7$–$2.7$ ms) sit far below the top effects ($34$–$138$ ms). Ten distinct regions take the top slot, the cut slows 11 participants and speeds 6, and 6 of the 17 top regions are of the mixed class, so the same cut moves other participants the other way. The class describes the region across the cohort, so it can disagree with the sign in the participant shown (sub-01, sub-19).

| Participant | base RT (ms) | band (ms) | robust | Strongest region | Role | $\Delta\mathrm{RT}$ (ms) | Class |
|---|---|---|---|---|---|---|---|
| sub-01 | $677$ | $0.92$ | $70$ | L mid. temporal (70) | pool | $+67.2$ | speeding |
| sub-02 | $643$ | $2.75$ | $43$ | R banks STS (30) | other | $-44.8$ | speeding |
| sub-03 | $634$ | $1.11$ | $65$ | brainstem (82) | other | $+52.7$ | mixed |
| sub-04 | $541$ | $1.06$ | $65$ | R pericalcarine (21) | injection | $+56.1$ | slowing |
| sub-06 | $696$ | $0.73$ | $69$ | R sup. parietal (17) | pool | $+43.5$ | mixed |
| sub-07 | $694$ | $1.57$ | $49$ | R sup. parietal (17) | pool | $-39.0$ | mixed |
| sub-08 | $770$ | $1.07$ | $61$ | R sup. parietal (17) | pool | $+92.2$ | mixed |
| sub-09 | $527$ | $1.87$ | $48$ | L inf. parietal (59) | pool | $+37.0$ | slowing |
| sub-10 | $680$ | $0.77$ | $62$ | L sup. parietal (58) | pool | $+58.6$ | slowing |
| sub-11 | $666$ | $1.09$ | $50$ | L mid. temporal (70) | pool | $+34.5$ | speeding |
| sub-12 | $822$ | $2.64$ | $39$ | L inf. temporal (69) | other | $-137.6$ | mixed |
| sub-13 | $606$ | $1.39$ | $59$ | R precentral (9) | literature | $+84.7$ | slowing |
| sub-14 | $716$ | $2.10$ | $34$ | L inf. temporal (69) | other | $-77.4$ | mixed |
| sub-17 | $636$ | $1.39$ | $39$ | L inf. parietal (59) | pool | $+54.3$ | slowing |
| sub-18 | $688$ | $1.84$ | $54$ | L fusiform (65) | other | $+38.4$ | slowing |
| sub-19 | $688$ | $1.08$ | $64$ | L sup. parietal (58) | pool | $-41.8$ | slowing |
| sub-20 | $784$ | $1.63$ | $44$ | R banks STS (30) | other | $-65.3$ | speeding |

Supplementary Table 8: **Time-gated silencings per participant.** Two blocks, one for each participant's strongest region and one for the parietal accumulator set (Fig. 7b,c): the signed $\Delta\mathrm{RT}$ (ms) when the silencing is applied for the whole trial, only during $0$–$200$ ms, or only during $300$–$600$ ms; the late-window scalp change $S_{300\text{-}600}$ under the three gates; and the fraction of the participant's decisions the intact model has made by $300$ ms (Fig. 7a). In the first block, sub-18's late-gate $\Delta\mathrm{RT}$ ($2 \times 10^{-4}$ ms) and $S_{300\text{-}600}$ ($3 \times 10^{-5}$) are below the printed precision.

| Participant | Silenced | $\Delta\mathrm{RT}$ (ms) | | | $S_{300\text{-}600}$ | | | decided |
|---|---|---|---|---|---|---|---|---|
| | | whole | early | late | whole | early | late | by $300$ ms |
| *Strongest region* | | | | | | | | |
| sub-01 | L mid. temporal (70) | $+67.2$ | $+41.0$ | $+5.5$ | $0.035$ | $0.002$ | $0.041$ | $0.66$ |
| sub-02 | R banks STS (30) | $-44.8$ | $-38.1$ | $-1.3$ | $0.231$ | $0.245$ | $0.005$ | $0.41$ |
| sub-03 | brainstem (82) | $+52.7$ | $+52.2$ | $+0.7$ | $0.365$ | $0.352$ | $0.086$ | $0.49$ |
| sub-04 | R pericalcarine (21) | $+56.1$ | $+47.7$ | $-0.3$ | $0.182$ | $0.165$ | $0.293$ | $0.66$ |
| sub-06 | R sup. parietal (17) | $+43.5$ | $+29.0$ | $+7.2$ | $0.080$ | $0.036$ | $0.042$ | $0.31$ |
| sub-07 | R sup. parietal (17) | $-39.0$ | $-28.4$ | $+6.8$ | $0.054$ | $0.015$ | $0.024$ | $0.31$ |
| sub-08 | R sup. parietal (17) | $+92.2$ | $+55.7$ | $+9.4$ | $0.333$ | $0.072$ | $0.135$ | $0.47$ |
| sub-09 | L inf. parietal (59) | $+37.0$ | $+16.3$ | $+7.0$ | $0.116$ | $0.012$ | $0.080$ | $0.71$ |
| sub-10 | L sup. parietal (58) | $+58.6$ | $+27.9$ | $+11.5$ | $0.086$ | $0.007$ | $0.054$ | $0.34$ |
| sub-11 | L mid. temporal (70) | $+34.5$ | $+14.1$ | $+8.8$ | $0.048$ | $0.009$ | $0.051$ | $0.41$ |
| sub-12 | L inf. temporal (69) | $-137.6$ | $-105.2$ | $-24.6$ | $0.037$ | $0.046$ | $0.046$ | $0.23$ |
| sub-13 | R precentral (9) | $+84.7$ | $+62.3$ | $+0.8$ | $0.570$ | $0.382$ | $0.212$ | $0.83$ |
| sub-14 | L inf. temporal (69) | $-77.4$ | $-35.2$ | $-6.9$ | $0.139$ | $0.013$ | $0.139$ | $0.21$ |
| sub-17 | L inf. parietal (59) | $+54.3$ | $+32.2$ | $+7.2$ | $0.075$ | $0.010$ | $0.035$ | $0.66$ |
| sub-18 | L fusiform (65) | $+38.4$ | $+10.7$ | $+0.0$ | $0.194$ | $0.075$ | $0.000$ | $0.48$ |
| sub-19 | L sup. parietal (58) | $-41.8$ | $-27.5$ | $-3.7$ | $0.057$ | $0.010$ | $0.057$ | $0.32$ |
| sub-20 | R banks STS (30) | $-65.3$ | $-30.7$ | $-14.0$ | $0.086$ | $0.080$ | $0.094$ | $0.23$ |
| *Parietal accumulator set* | | | | | | | | |
| sub-01 | accumulator set | $-131.5$ | $-123.1$ | $-13.7$ | $0.078$ | $0.016$ | $0.035$ | $0.66$ |
| sub-02 | accumulator set | $+7.8$ | $+20.3$ | $-10.5$ | $0.133$ | $0.096$ | $0.031$ | $0.41$ |
| sub-03 | accumulator set | $+100.0$ | $+46.7$ | $+20.5$ | $0.304$ | $0.018$ | $0.293$ | $0.49$ |
| sub-04 | accumulator set | $-48.8$ | $-42.5$ | $+4.3$ | $0.507$ | $0.077$ | $0.326$ | $0.66$ |
| sub-06 | accumulator set | $+7.2$ | $-7.3$ | $-3.3$ | $0.096$ | $0.039$ | $0.045$ | $0.31$ |
| sub-07 | accumulator set | $-22.2$ | $-41.7$ | $+23.9$ | $0.117$ | $0.006$ | $0.118$ | $0.31$ |
| sub-08 | accumulator set | $+42.4$ | $+21.3$ | $+12.2$ | $0.358$ | $0.080$ | $0.192$ | $0.47$ |
| sub-09 | accumulator set | $+31.8$ | $+34.7$ | $+7.1$ | $0.214$ | $0.017$ | $0.169$ | $0.71$ |
| sub-10 | accumulator set | $+76.0$ | $+41.2$ | $+10.9$ | $0.121$ | $0.008$ | $0.087$ | $0.34$ |
| sub-11 | accumulator set | $-32.5$ | $-18.3$ | $-7.9$ | $0.128$ | $0.005$ | $0.125$ | $0.41$ |
| sub-12 | accumulator set | $-20.1$ | $-11.5$ | $-34.0$ | $0.146$ | $0.105$ | $0.079$ | $0.23$ |
| sub-13 | accumulator set | $+132.6$ | $+66.3$ | $+7.4$ | $0.434$ | $0.185$ | $0.199$ | $0.83$ |
| sub-14 | accumulator set | $+26.7$ | $+19.3$ | $+8.0$ | $0.085$ | $0.023$ | $0.066$ | $0.21$ |
| sub-17 | accumulator set | $+57.2$ | $+37.9$ | $+3.3$ | $0.227$ | $0.014$ | $0.049$ | $0.66$ |
| sub-18 | accumulator set | $+12.3$ | $+16.4$ | $+9.2$ | $0.187$ | $0.131$ | $0.077$ | $0.48$ |
| sub-19 | accumulator set | $-138.6$ | $-123.9$ | $-10.5$ | $0.202$ | $0.018$ | $0.150$ | $0.32$ |
| sub-20 | accumulator set | $-53.7$ | $-36.0$ | $+15.9$ | $0.130$ | $0.050$ | $0.119$ | $0.23$ |

Supplementary Table 9: **Notation and dimensions.** Indices, the trial-specific latent variable, dynamical states, the decision readout, and the frozen anatomical operators. The logistic function is $\varphi(x) = (1 + e^{-x})^{-1}$.

| Symbol | Meaning | Dimension / values |
|---|---|---|
| $i, j$ | region indices | $1, \ldots, R,\ R = 83$ (Lausanne-2018 scale 1) |
| $c$ | EEG channel index | $1, \ldots, C,\ C = 128$ |
| $n$ | trial index (batch $B$) | $1, \ldots, N$ |
| $t$ | dynamics step | $1, \ldots, T_{\text{dyn}},\ T_{\text{dyn}} = 200$ |
| $s$ | observation sample | $1, \ldots, N_{\text{obs}},\ N_{\text{obs}} = 100$ (100 Hz) |
| $z_n$ | trial-specific latent variable | $\mathbb{R}^1$ ($z$-dim $= 1$) |
| $\tilde{z}_n$ | within-condition residual of $z_n$ | $\mathbb{R}^1$, zero-mean per condition |
| $E_i, I_i$ | WC excitatory / inhibitory activity | scalar in $[0, 1)$ |
| $\text{in}_{E,i}$ | net pyramidal current (EEG source = evidence) | scalar |
| $\mathcal{D}$ | decision pool | 10 parcels |
| $e_n(t)$ | pooled evidence (Eq. 20) | scalar |
| $v_n(t)$ | accumulator drift (Eq. 21) | scalar |
| $C_{ij}$ | structural connectome (SC), frozen | $(R, R)$, normalized to $[0, 1]$ |
| $L_{ij}$ | mean fiber length, frozen | $(R, R)$, symmetric, zero diagonal (mm) |
| $\mathbf{L}$ | leadfield, frozen | $(C, R)$, unit Frobenius norm |
| $D_{ij}$ | conduction-delay steps | $(R, R),\ \text{round}(L_{ij}/(v\, \text{dt}_{\text{dyn}}))$ |

Supplementary Table 10: **Cohort and inclusion.** Twenty participants were acquired. Two (sub-05, sub-15) were dropped before fitting because the dataset did not process their motion-task EEG, citing excessive noise[5], leaving 18 fitted participants, every one of which clears the criterion fixed in advance on condition-mean ERP correlation ($r > 0.60$; Methods Dataset and task, Face-validity checks). One of them, sub-16, clears that criterion at $r = 0.830$ but contributes only 216 usable trials, below the $\geq 250$-trial criterion (this participant inverted the response keys on several trials[5]), and is excluded from every cohort statistic and figure. Fig. 2 and every analysis after it therefore run at $N = 17$; sub-16's face-validity values are kept in Supplementary Table 2. $n$, analyzed single trials.

| Participant | $n$ trials | ERP $r$ | Cohort |
|---|---|---|---|
| sub-01 | 569 | 0.632 | face validity + downstream |
| sub-02 | 497 | 0.755 | face validity + downstream |
| sub-03 | 571 | 0.925 | face validity + downstream |
| sub-04 | 554 | 0.878 | face validity + downstream |
| sub-05 | – | – | excluded (EEG not processed) |
| sub-06 | 571 | 0.677 | face validity + downstream |
| sub-07 | 520 | 0.800 | face validity + downstream |
| sub-08 | 561 | 0.877 | face validity + downstream |
| sub-09 | 498 | 0.854 | face validity + downstream |
| sub-10 | 520 | 0.858 | face validity + downstream |
| sub-11 | 559 | 0.791 | face validity + downstream |
| sub-12 | 471 | 0.783 | face validity + downstream |
| sub-13 | 581 | 0.941 | face validity + downstream |
| sub-14 | 550 | 0.880 | face validity + downstream |
| sub-15 | – | – | excluded (EEG not processed) |
| sub-16 | 216 | 0.830 | face validity only (too few trials) |
| sub-17 | 577 | 0.877 | face validity + downstream |
| sub-18 | 557 | 0.859 | face validity + downstream |
| sub-19 | 524 | 0.805 | face validity + downstream |
| sub-20 | 565 | 0.860 | face validity + downstream |

Supplementary Table 11: **Parameter ledger, freeze/train schedule, and fixed constants.** How every quantity is handled, the step at which it is trained, its constraint transform, and its initial or fixed value (Methods Frozen anatomical scaffold through Fitting protocol). "Adam" is updated by Adam[9] at the steps listed under *trained at*, which are the warm-start (WS), the curriculum stages A, B, and C, the re-anchor (RA), and the recalibration (RC), and is held fixed at every other step; *never* marks a quantity that enters no gradient. The inhibitory weights $w_{IE,i}$ are set by offline feedback-inhibition control before warm-start (Supplementary Note 4) and never enter any gradient. ClipT$[a, b]$ is a clipped reparameterization onto $[a, b]$; SoftplusT$(\epsilon)$ a positive softplus with floor $\epsilon$; an empty transform cell means the raw parameter is fit directly. The last two blocks collect the fixed constants and the fitting hyperparameters used for every participant, with the meaning of each in the second column; anatomy ($C$, $L$, leadfield $\mathbf{L}$, delays $D$) is calibrated once and never updated by gradients.

| Quantity | Handling | Trained at | Transform | Init / value |
|---|---|---|---|---|
| *Frozen anatomy* | | | | |
| leadfield $\mathbf{L}$ | fixed | never | unit Frobenius | forward model |
| connectome $C$ | fixed | never | norm. $[0, 1]$ | DWI connectome |
| fiber length $L$ | fixed | never | | CMP parcellation |
| *Offline-calibrated, then frozen* | | | | |
| $w_{IE,i}$ (FIC) | fixed-point solve on $W{=}G_0C$ | before WS | clip $[0.05, 10]$ | coupling-aware |
| $v$ (conduction speed) | frozen | never | | 3 m/s |
| $G_0$ ($W^0$ anchor) | frozen | never | | 0.5 |
| *Trained at warm-start and re-anchor, frozen for the curriculum (TOPO)* | | | | |
| $s_i, r_j$ (out/in gains) | Adam $+\mathcal{R}_{\text{ec}}$ | WS, RA | | $\mathbf{1}$ |
| $U_1, V_1$ (rank-4, masked $C \odot U_1 V_1^\top$) | Adam $+\mathcal{R}_{\text{ec}}$ | WS, RA | | $0.01\,\mathcal{N}$ |
| $\tau_m$ (neuromod. time const.) | Adam | WS, RA | | 20 ms |
| $G$ (global coupling) | Adam | WS, RA | | 0.5 |
| $a_{\text{vis}}, a_{\text{mot}}$ (stimulus gains) | Adam | WS, RA | | 1.0, 1.0 |
| $q_i$ (distributed input field) | Adam $+\mathcal{R}_{\text{ec}}$ | WS, RA | | $\mathbf{0}$ |
| *Substrate (Param, frozen across the curriculum)* | | | | |
| $\tau_{E,i}, \tau_{I,i}$ | left at init | never | ClipT$[2, 50]$ | 10, 15 ms |
| $\sigma_E, \sigma_I$ | left at init | never | SoftplusT$(10^{-3})$ | 0.05 |
| *Curriculum: latent / observation (Stage A, C)* | | | | |
| $\lambda$ ($z \to$pool scale) | Adam | A, C | | 0.5 |
| encoder $\phi$ | Adam | A, C | | amortized net |
| $g$ (EEG gain) | Adam | A, C | | 1.0 |
| $\log \sigma_{\text{obs}}$ | Adam | A, C | | 0.0 |
| *Curriculum: decision (Stage B, C)* | | | | |
| $\beta$ (condition-mean drift gain) | Adam | B, C, RC | | 0.5 |
| $\beta_{\text{trial}}$ (trial drift gain) | Adam | C | | 0.5 |
| $\eta$ (pool readout logits) | Adam | B, C, RC | softmax | $\mathbf{0}$ (uniform) |
| $b$ (drift bias) | Adam | B, C, RC | | $-0.1$ |
| $a$ (boundary) | Adam | B, C, RC | ClipT$[0.3, 5]$ | 1.0 |
| $w$ (start ratio) | Adam | B, C, RC | ClipT$[0.05, 0.95]$ | 0.5 |
| $t_0$ (non-decision, s) | Adam | B, C, RC | ClipT$[0.05, 0.6]$ | 0.15 |
| $\sigma_a$ (accumulator noise) | fixed (diffusion gauge) | never | SoftplusT$(0.1)$ | 1.0 |
| *Fixed constants (never trained)* | | | | |
| epoch window | pre / post stimulus | | | $[-200, +800]$ ms |
| $\text{dt}_{\text{dyn}}$ | dynamics step | | | 5 ms |
| $T_{\text{dyn}}$ | dynamics steps / epoch | | | 200 |
| EEG rate | observation sampling | | | 100 Hz ($\text{dt}_{\text{obs}} = 10$ ms) |
| $N_{\text{obs}}$ | observation samples / epoch | | | 100 |
| stride | dynamics-to-observation | | | $T_{\text{dyn}}/N_{\text{obs}} = 2$ |
| stim. duration | sensory square wave | | | 200 ms |
| $w_{EE}, w_{EI}, w_{II}$ | recurrent coupling weights | | | $1.5,\ 1.0,\ 0.5$ |
| $a_E, \theta_E, a_I, \theta_I$ | sigmoid gains, thresholds | | | $1.2,\ 2.8,\ 1.0,\ 1.0$ |
| $b_E$ | tonic background to E | | | 2.5 |
| $\tau_{\text{psp}}$ | EEG post-synaptic constant | | | 50 ms |
| $n_{\text{grid}}$ | Fokker–Planck interior bins | | | 128 |
| $\lambda_{\text{fb}}$ | free-bits floor | | | 1.0 |
| $\gamma_{\text{wp}}, \gamma_{\text{reg}}, \gamma_{\text{rt}}$ | regularizer weights | | | $0.10,\ 5 \times 10^{-3},\ 100$ |

*continued on the next page*

Supplementary Table 11, continued.

| Quantity | Handling | Trained at | Transform | Init / value |
|---|---|---|---|---|
| *Fitting hyperparameters* | | | | |
| FIC | inhibition calibration | | | $600$ steps, $\eta = 0.05$ |
| warm-start | topology pre-fit | | | lr $10^{-2}$, $5{,}000$ epochs |
| curriculum | stages A/B/C epochs | | | lr $2 \times 10^{-3}$, $(300, 200, 200)$ |
| re-anchor | topology refit | | | lr $10^{-2}$, $800$ epochs |
| recalibration | decision refit | | | lr $2 \times 10^{-3}$, $100$ epochs |
| optimizer | | | | Adam, grad-norm clip $1.0$ |

Supplementary Table 12: **Synthetic-data recovery test.** Recovery on trials generated by the frozen model and refit by the same curriculum. Two participants, sub-14 and sub-09, were chosen in advance by the size of the forward EEG imprint of their latent variable, and each was swept over an $11 \times 11$ $\beta \times \lambda$ grid, giving $121$ refits per participant. $|r|$ is the within-condition latent correlation at the base cell against the $0.20$ threshold, whose information ceiling is $\approx 0.46$; separability is a bootstrap partial correlation of each recovered parameter against its own true value, holding the other fixed, and clears when the interval excludes zero. Both participants clear both criteria. The reseed column counts how often the base cell cleared $0.20$ across ten fresh master seeds, each resetting the synthetic keys and the curriculum's random state; this is a readout of the sampling distribution and not the statistic fixed in advance, which is defined on the single base cell. Only the product $\beta\lambda$ scales per-trial evidence, so its recovery is reported alongside. sub-10 and sub-18 were swept on the same grid as reference points on the recovery floor, by their smaller imprint, and are not part of the test. Supplementary Fig. 7 draws the two test participants on the same sweep, giving the sampling distribution behind the reseed column and the intervals behind the separability criterion. Methods Identifiability and synthetic-data recovery.

| | | | separability (partial $r$ [95% CI]) | | |
|---|---|---|---|---|---|
| Participant | $\lvert r\rvert$ latent | reseed pass | $\beta$ | $\lambda$ | $\beta\lambda$ |
| *Test participants* | | | | | |
| sub-14 | $0.233$ | 9/10 | $0.958$ $[0.946, 0.968]$ | $0.400$ $[0.246, 0.536]$ | $0.642$ |
| sub-09 | $0.340$ | 9/10 | $0.946$ $[0.929, 0.963]$ | $0.533$ $[0.421, 0.645]$ | $0.187$ |
| *Swept, not part of the test* | | | | | |
| sub-10 | $0.121$ | – | $0.952$ $[0.941, 0.963]$ | $-0.008$ $[-0.188, 0.214]$ | $0.159$ |
| sub-18 | $0.118$ | – | $0.974$ $[0.964, 0.983]$ | $0.173$ $[0.020, 0.330]$ | $0.479$ |

Supplementary Table 13: **Focus regions and the literature region set.** The 19 regions of Fig. 5c with their standing before the scan: *known (axis)*, in the literature set, with its functional axis; *model prior*, a V1 injection site, outside the literature set, so that an effect there follows from the specification; *new*, in neither. Then the number of participants placing the region among their eight strongest regions, the number it robustly slowed and sped, the cohort median $\Delta\mathrm{RT}$ (ms; positive slows), and the cohort class. Every entry is a hypothesis generated inside the model, not a tested effect. The literature set comprises 18 regions in 9 bilateral pairs implicated by causal-perturbation studies of perceptual decisions, given here as region (Lausanne-2018 scale-1 parcellation label; parcel indices; axis): MT/V5[10] (middletemporal; 29, 70; evidence), intraparietal / superior parietal[11] (superiorparietal; 17, 58; accumulation), inferior parietal lobule (inferiorparietal; 18, 59; accumulation), supramarginal gyrus (supramarginal; 16, 57; accumulation), posterior parietal / precuneus (precuneus; 19, 60; accumulation), dorsolateral prefrontal cortex[12] (rostralmiddlefrontal; 6, 47; control), frontal eye field / premotor, taken as the caudal middle-frontal proxy[13] (caudalmiddlefrontal; 8, 49; control), pre-SMA / SMA[14] (superiorfrontal; 7, 48; threshold), and primary motor cortex[15,16] (precentral; 9, 50; motor). The first five pairs make up the model's decision pool.

| Region | Standing | Role | top-8 | slowed | sped | median $\Delta\mathrm{RT}$ (ms) | Class |
|---|---|---|---|---|---|---|---|
| R sup. frontal (7) | known (threshold) | literature | 4 | 12 | 3 | $+3.8$ | slowing |
| L sup. frontal (48) | known (threshold) | literature | 2 | 11 | 2 | $+4.1$ | slowing |
| R precentral (9) | known (motor) | literature | 4 | 9 | 2 | $+1.3$ | slowing |
| L precentral (50) | known (motor) | literature | 2 | 12 | 1 | $+2.9$ | slowing |
| L postcentral (56) | new | – | 4 | 15 | 1 | $+8.6$ | slowing |
| R sup. parietal (17) | known (accumulation) | pool | 11 | 8 | 6 | $-0.1$ | mixed |
| L sup. parietal (58) | known (accumulation) | pool | 8 | 9 | 7 | $+5.8$ | slowing |
| R inf. parietal (18) | known (accumulation) | pool | 4 | 7 | 8 | $+0.6$ | mixed |
| L inf. parietal (59) | known (accumulation) | pool | 6 | 9 | 7 | $+1.8$ | slowing |
| R precuneus (19) | known (accumulation) | pool | 5 | 6 | 8 | $+0.2$ | mixed |
| R cuneus (20) | model prior | injection | 0 | 3 | 7 | $-0.8$ | mixed |
| L cuneus (61) | model prior | injection | 2 | 7 | 7 | $-0.1$ | mixed |
| R pericalcarine (21) | model prior | injection | 5 | 9 | 5 | $+1.7$ | slowing |
| L pericalcarine (62) | model prior | injection | 2 | 11 | 4 | $+2.1$ | slowing |
| R lat. occipital (22) | new | – | 7 | 6 | 10 | $-4.9$ | speeding |
| L fusiform (65) | new | – | 6 | 9 | 6 | $+2.5$ | slowing |
| R mid. temporal (29) | known (evidence) | pool | 3 | 1 | 14 | $-5.4$ | speeding |
| L mid. temporal (70) | known (evidence) | pool | 7 | 5 | 11 | $-6.7$ | speeding |
| L sup. temporal (72) | new | – | 4 | 5 | 9 | $-1.6$ | speeding |

Supplementary Table 14: **Which model produced which number.** Every fit shares the frozen anatomy and forward pass of the main model; they differ in the latent variable, in what is refit, and in which trials the score reads. *In sample*: scored on the trials the fit used. *Held out*: every scored trial was absent from the fit that predicts it.

| Model or estimator | What it is | Where its numbers appear |
|---|---|---|
| Main fit | One-dimensional latent variable $z_n$ per trial, read by a linear separable encoder from the whole epoch; joint EEG and reaction-time objective; all trials. In sample. | Figs. 2b–h, 4–7; the in-sample $51.6\%$ of Fig. 3b |
| Strict five-fold refit | The whole pipeline refit without each fifth of the trials; every condition mean from the training trials; slope-1 $R^2$. Held out. | Fig. 3a (first column), c, d; the $9.6\%$ of EEG-inferred latent variables predict held-out reaction times when behavior is in the fit |
| Held-out encoder | Two-dimensional recognition encoder with a slow first-order autoregressive state across trials, refit per fold; decision parameters from the main fit. Held out. | Fig. 3a (encoder columns); Supplementary Table 3 |
| EEG-only and behavior-only baselines | EEG-only fit with a post hoc readout; per-condition drift-diffusion model. In sample. | Fig. 3b; Supplementary Table 1 |
| Model variance budget | Law-of-total-variance partition of the main fit's own reaction-time variance (Eq. S1). Neither in sample nor held out: not a prediction score. | Supplementary Fig. 3; Supplementary Note 2 |

### 7 Supplementary Figures

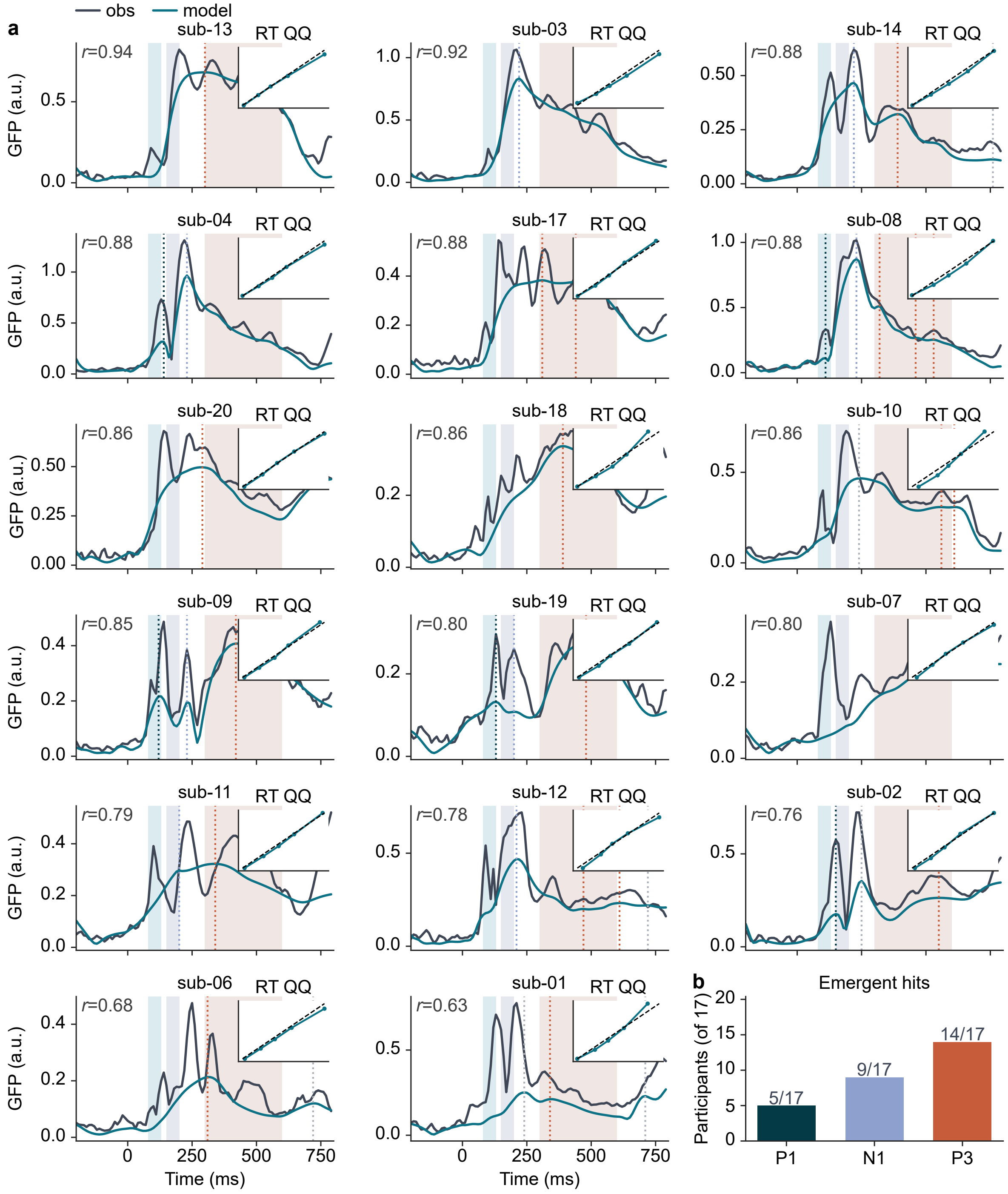

Supplementary Figure 1: **Face validity of each participant in the cohort ($N = 17$). a,** One panel per participant, expanding Fig. 2e, g, and h from cohort quantiles to individuals: observed (black) and model (blue) condition-mean global field power from $-200$ to $800$ ms, with the P1, N1, and P3 search windows shaded, detected peaks marked by dotted lines, and a reaction-time quantile–quantile plot inset. Panels are ordered by ERP correlation from highest to lowest, from sub-13 ($r = 0.941$) to sub-01 ($r = 0.632$), each printing its own value to two decimals; unrounded values are in Supplementary Table 2. A peak falling in two windows is drawn in the color of the earlier one, and the hit counts of **b** take precedence; each panel scales its own $y$ axis, so amplitudes are compared in Fig. 2e. **b,** Number of participants with an emergent global-field-power peak in each window within $\pm 30$ ms: P1 $5/17$, N1 $9/17$, P3 $14/17$.

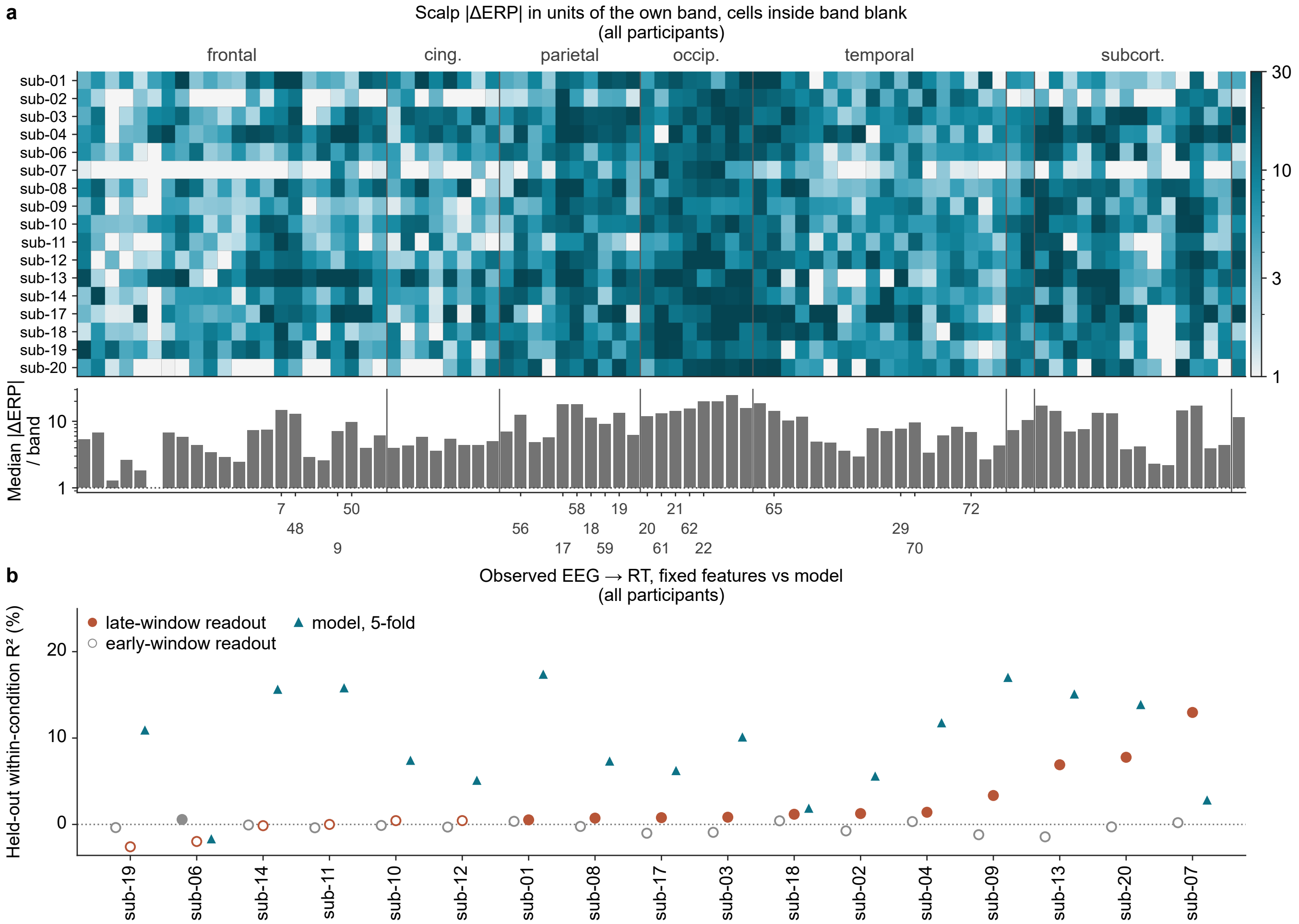


Supplementary Figure 2: **The scalp side of the intervention map for the whole cohort, and the observed-EEG readout. a,** The scalp magnitude of $\Delta\mathrm{ERP}$ for every region and participant, in units of the participant's own reseeded ERP band, in the anatomical order of Fig. 5b so that the two matrices read column for column; cells inside the band are blank (logarithmic color, clipped at 30 times the band). $1{,}280$ of $1{,}411$ cells are robust on the scalp and $896$ on both readouts; and the ratio of the strongest region's effect to the band runs from $29\times$ to $72\times$. In sub-08, 60 regions are robust on both readouts, 21 on the scalp only, 1 on behavior only, and 1 on neither. Below: the cohort median of the same ratio per region (logarithmic axis; dotted line, the band), largest for the right lingual gyrus (23; 25), the two lateral occipital parcels (63, 22; 20), the right fusiform gyrus (24; 19), and the two superior parietal parcels (17, 58; 18); the index labels mark the focus regions of Fig. 5c. **b,** Held-out within-condition $R^2$ of reaction time from observed EEG, per participant, under the folds of Fig. 3: a two-feature linear readout of the early window and of the late window (centro-parietal amplitude and training-fold template projection), against the whole model refit within each fold (Fig. 3a, first column). Cohort means $-0.3\%$, $2.0\%$, and $9.6\%$; the two-window readout reaches $2.4\%$ (not drawn); filled markers pass a within-condition permutation null at $P < 0.05$ (late: 11 of 17; early: 1 of 17). The late-window slope is negative in 16 of 17.

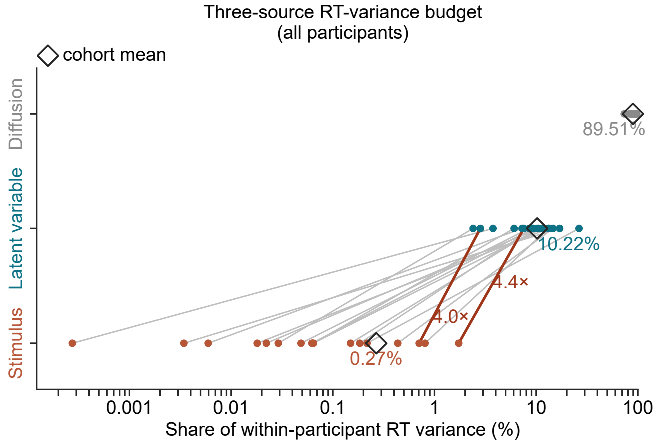


Supplementary Figure 3: **The variance budget.** The main fit's variance budget behind Fig. 4, over all of a participant's trials (logarithmic axis, one row per source; the stimulus row is the part that differs between conditions): filled points are each participant's share and hollow diamonds the cohort mean. Diffusion $89.51\%$, latent variable $z_n$ $10.22\%$, stimulus $0.27\%$. Shares rather than absolute variances are shown because the diffusion term receives the remaining unattributed variance and absorbs the bias from truncating the computational grid (*leak* column of Supplementary Table 4). Connectors pair each participant's stimulus share with its latent-variable share: latent-variable shares run from $2.40\%$ in sub-10 to $26.40\%$ in sub-09, exceed the stimulus share in 17/17, and do so by at least tenfold in 15/17; the other two are drawn in a different color with their ratios printed.

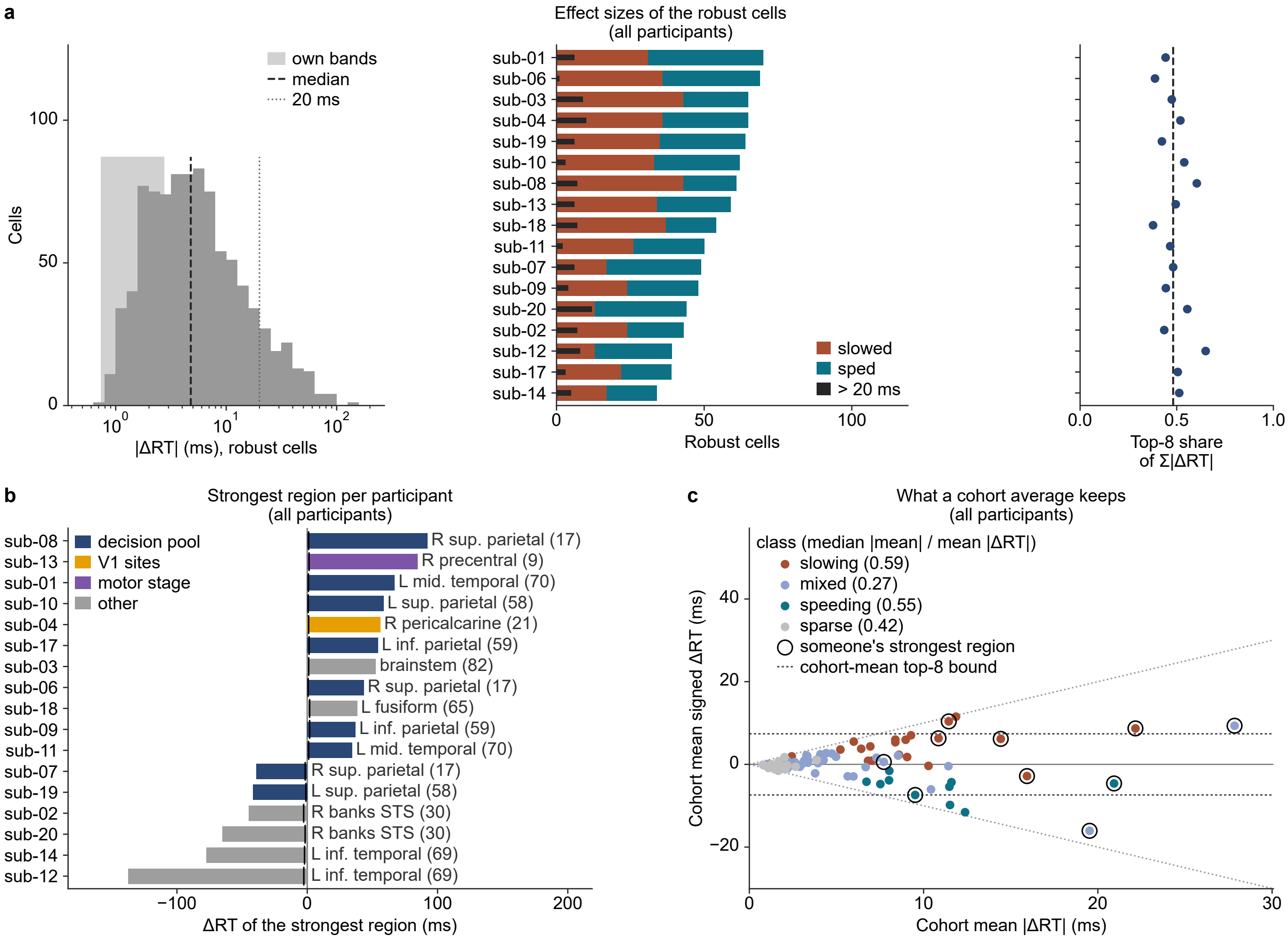


Supplementary Figure 4: **The cohort behind the intervention map.** All panels read the forward runs of Fig. 5; a cell is robust when its $|\Delta\mathrm{RT}|$ exceeds the participant's own reseeded noise band. **a,** Effect sizes of the $915$ robust cells. Left: their $|\Delta\mathrm{RT}|$ on a logarithmic axis, the range of the 17 noise bands shaded, the cohort median dashed ($4.8$ ms; interquartile range $2.4$–$10.0$ ms), and $20$ ms dotted ($102$ cells beyond it). Middle: per participant, ordered by the number of robust cells, the cells that slowed ($484$ in all) and sped ($431$), with the cells beyond $20$ ms in black (median 6 per participant, range 1–12). Right: the share of each participant's summed $|\Delta\mathrm{RT}|$ over all 83 regions carried by its eight strongest regions (median $0.48$, range $0.38$–$0.65$). **b,** The strongest region of each participant (Supplementary Table 7): its signed $\Delta\mathrm{RT}$ as a bar colored by the region's stage as in Fig. 5a, the region named, and the participant's own noise band as a black tick beside zero. Ten distinct regions take the top slot; the silencing slows 11 participants and speeds 6, with $|\Delta\mathrm{RT}|$ from $34$ to $138$ ms; by stage, 9 are decision-pool regions, 1 an injection site, 1 a motor-stage proxy, and 6 outside all three sets. **c,** What a cohort average keeps: for every region, the cohort mean of the signed $\Delta\mathrm{RT}$ against the cohort mean of $|\Delta\mathrm{RT}|$, colored by cohort class; a region on either dotted diagonal keeps all of its magnitude in the average, one on the zero line none. The key gives each class's median ratio of $|$mean signed $\Delta\mathrm{RT}|$ to mean $|\Delta\mathrm{RT}|$: $0.59$ slowing, $0.27$ mixed, $0.55$ speeding, and $0.42$ sparse. Rings mark the regions that are some participant's strongest region, and the dashed rules the eighth-largest $|$cohort mean$|$: the eight regions with the largest signed cohort mean hold the strongest region of 10 of the 17 participants (12 of 17 when ranked by the cohort mean of $|\Delta\mathrm{RT}|$).

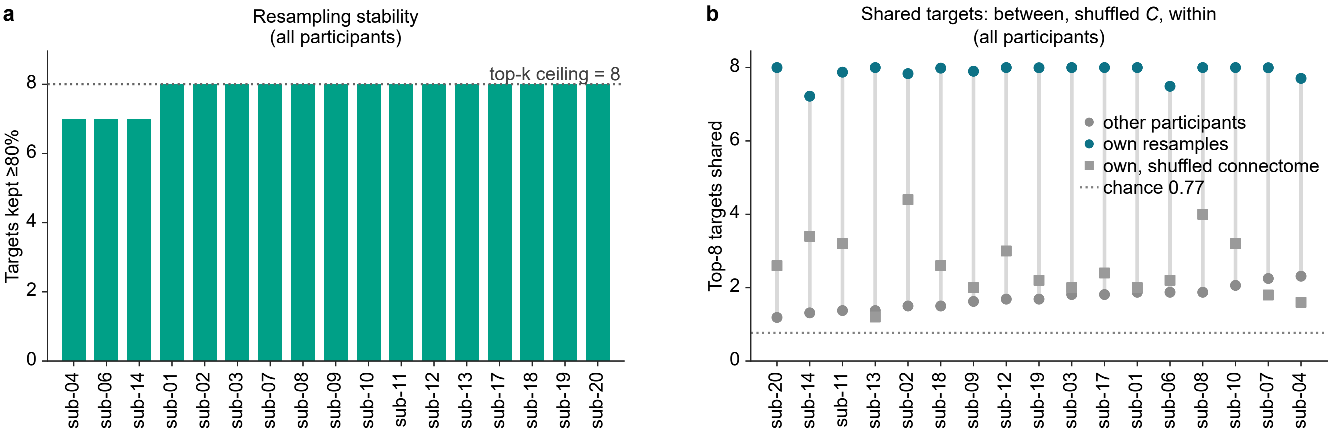


Supplementary Figure 5: **What bounds the intervention maps. a,** Resampling stability: the number of each participant's eight pre-fixed targets with inclusion frequency $\geq 0.8$ over 200 trial resamples, in practice $7$–$8$; the criterion fixed in advance requires at least 9 of 17 participants to retain robust targets (decision threshold $8.5$), and 17/17 do. **b,** Overlap of the eight strongest regions: for each participant, the mean overlap of its eight pre-fixed targets with the other 16 participants' targets (gray; mean $1.71$, at most $5$; dotted line, the expected overlap of two random 8-subsets of 83 regions, $0.77$), with the eight strongest regions of the same participant's model refit on a degree-preserving shuffle of its own connectome (squares; five shuffles, mean $2.58$), and with the eight strongest of 200 trial resamples of the same participant (blue; the sum of inclusion frequencies, $7.2$–$8$, all eight in 8 participants).

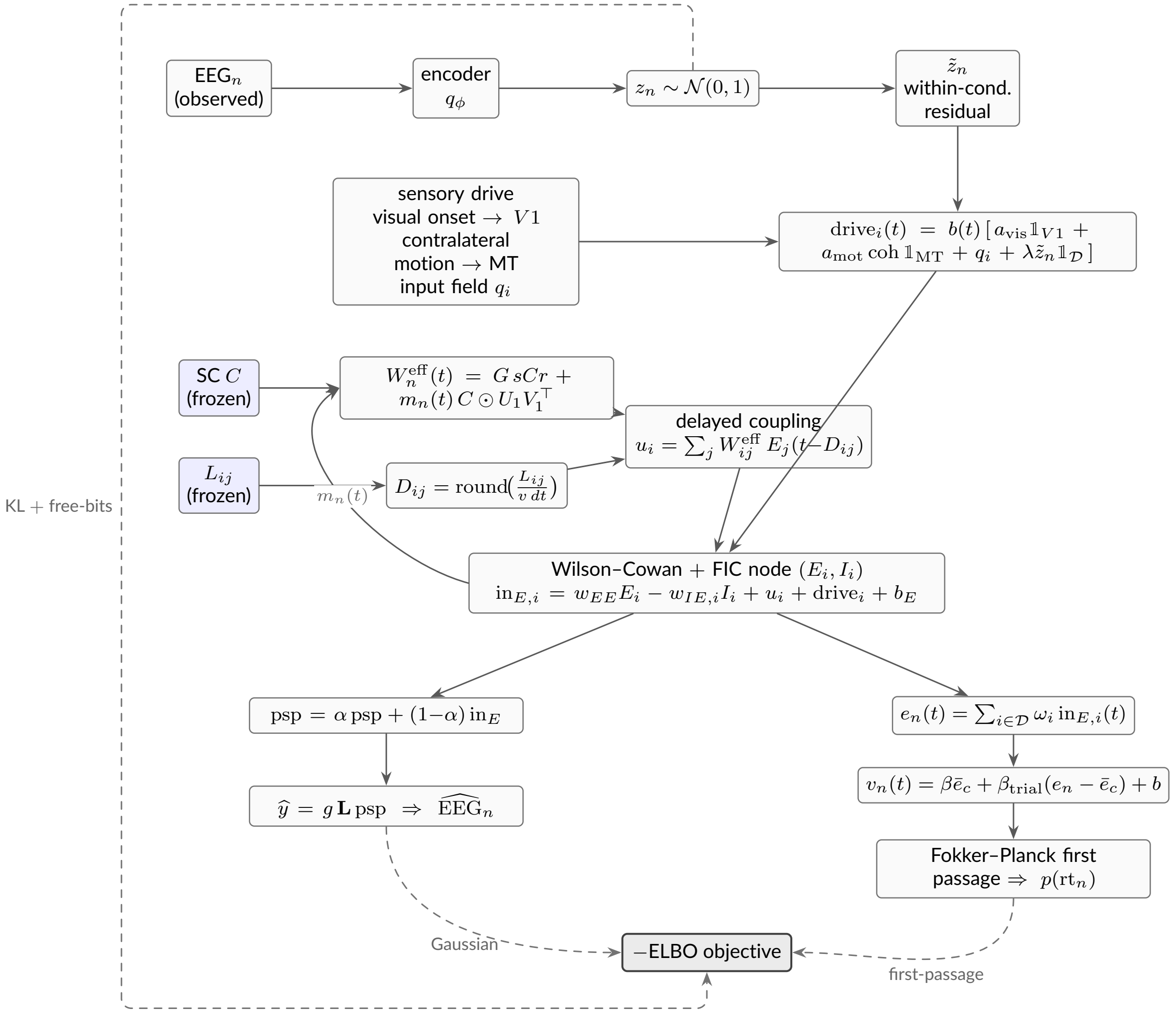


Supplementary Figure 6: **Joint generative model and data flow.** Nodes are tensors or states; solid arrows are the generative forward pass, dashed arrows are the likelihood and regularizer terms entering the negative ELBO. Shaded boxes are frozen anatomical operators. The EEG-only encoder $q_\phi$ infers the latent variable $z_n$; its within-condition residual $\tilde{z}_n$ is injected additively into the decision pool $\mathcal{D}$, while the sensory channels ($V1$, MT, $q$) are $z$-invariant. Regional net current $\text{in}_{E,i}$ is read out as EEG (slow PSP through the frozen leadfield) and as decision evidence (weighted pool readout driving a Fokker–Planck first-passage process). The neuromodulatory scalar $m_n(t)$ feeds back from mean activity to the effective connectivity $W_n^{\text{eff}}(t)$.

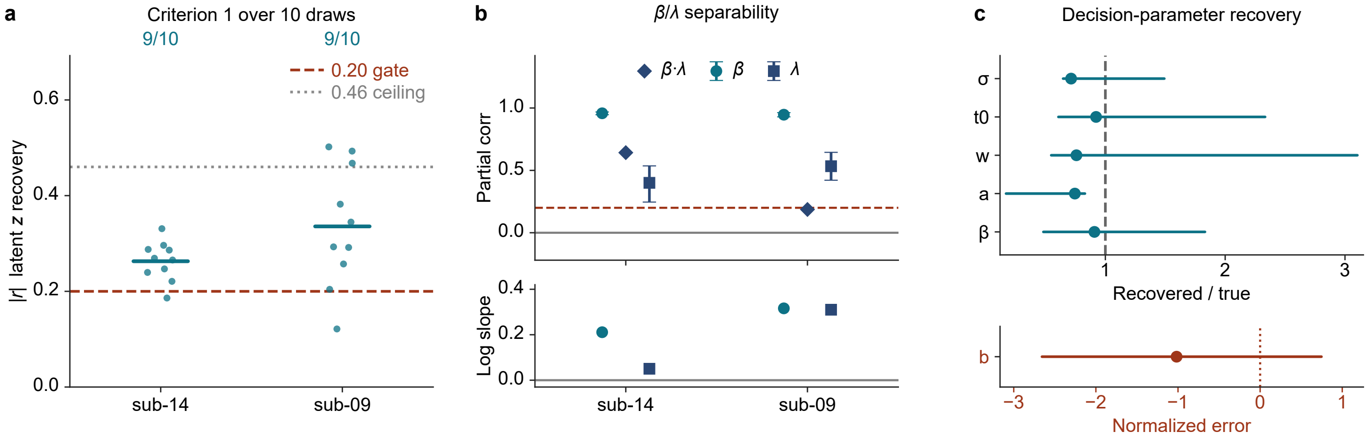


Supplementary Figure 7: **Identifiability criteria and their dependence on sampling.** Backs the synthetic-data recovery test of Supplementary Note 5. All three panels show the two test participants, sub-14 and sub-09, on the $11 \times 11$ sweep from which the criteria were read, so a value may be compared across panels as well as within one; the outcomes are in Supplementary Table 12. $\beta$ and $\lambda$ each carry identifiable information, but the partial correlation and the recovery slope do not ask the same question. **a,** Sampling distribution of the latent-recovery criterion: each participant re-drawn under ten fresh master seeds against the $0.20$ threshold, with the mean marked and the pass count printed, $9/10$ for both. The statistic fixed in advance is the single base cell, $|r| = 0.233$ and $0.340$, and is reported in Supplementary Table 12 rather than marked here, so that the panel is read as spread and the table as outcome. **b,** Separability and slope. Top, each participant's partial correlations with bootstrap intervals and the $\beta\lambda$ product printed; bottom, recovery slopes on log–log axes. $\lambda$ clears the criterion in both participants, $+0.400$ for sub-14 and $+0.533$ for sub-09, while their log–log slopes are $+0.050$ and $+0.310$ against $+0.211$ and $+0.316$ for $\beta$. With 121 cells an interval excludes zero above $r = 0.18$, so the plotted $0.20$ line and the interval both bind. **c,** Decision-parameter recovery pooled over the two participants and 121 cells each: the positive parameters $(\beta, a, w, t_0, \sigma)$ share a recovered-to-true ratio axis with a reference at $1$, while the drift bias $b$, which is $\leq 0$ and near-degenerate, uses a normalized error on a separate row. Marker, median; whisker, $2.5$–$97.5\%$. $\beta$, $w$, $t_0$, and $\sigma$ have intervals covering $1$ ($\sigma_a$ is free in the configuration this test used; Methods Identifiability and synthetic-data recovery); the bound $a$ falls below $1$ across its whole interval, and the medians of $a$, $w$, and $\sigma$ all sit near $0.75$; $b$ is weakly identifiable. Two participants rather than the whole cohort is a property of the question: this is a self-recovery test of the model and pipeline, not a population claim.
